# Discovery and functional interrogation of SARS-CoV-2 protein-RNA interactions

**DOI:** 10.1101/2022.02.21.481223

**Authors:** Joy S. Xiang, Jasmine R. Mueller, En-Ching Luo, Brian A. Yee, Danielle Schafer, Jonathan C. Schmok, Frederick E. Tan, Katherine Rothamel, Rachael N. McVicar, Elizabeth M. Kwong, Krysten L. Jones, Hsuan-Lin Her, Chun-Yuan Chen, Anthony Q. Vu, Wenhao Jin, Samuel S. Park, Phuong Le, Kristopher W. Brannan, Eric R. Kofman, Yanhua Li, Alexandra T. Tankka, Kevin D. Dong, Yan Song, Aaron F. Carlin, Eric L. Van Nostrand, Sandra L. Leibel, Gene W. Yeo

## Abstract

The COVID-19 pandemic is caused by severe acute respiratory syndrome-coronavirus-2 (SARS-CoV-2). The betacoronvirus has a positive sense RNA genome which encodes for several RNA binding proteins. Here, we use enhanced crosslinking and immunoprecipitation to investigate SARS-CoV-2 protein interactions with viral and host RNAs in authentic virus-infected cells. SARS-CoV-2 proteins, NSP8, NSP12, and nucleocapsid display distinct preferences to specific regions in the RNA viral genome, providing evidence for their shared and separate roles in replication, transcription, and viral packaging. SARS-CoV-2 proteins expressed in human lung epithelial cells bind to 4773 unique host coding RNAs. Nine SARS-CoV-2 proteins upregulate target gene expression, including NSP12 and ORF9c, whose RNA substrates are associated with pathways in protein N-linked glycosylation ER processing and mitochondrial processes. Furthermore, siRNA knockdown of host genes targeted by viral proteins in human lung organoid cells identify potential antiviral host targets across different SARS-CoV-2 variants. Conversely, NSP9 inhibits host gene expression by blocking mRNA export and dampens cytokine productions, including interleukin-1α/β. Our viral protein-RNA interactome provides a catalog of potential therapeutic targets and offers insight into the etiology of COVID-19 as a safeguard against future pandemics.

## Introduction

COVID-19 is caused by the novel severe acute respiratory syndrome coronavirus 2 (SARS-CoV-2), a positive-sense single-stranded (+ss)RNA virus. The viral genome encodes 29 proteins^1^, which include the four structural proteins, membrane or matrix (M), nucleocapsid (N), envelope (E), and spike (S) proteins. In addition, there are 16 non-structural proteins, NSP1-16, and 9 accessory proteins ORF3a-ORF10, though the expression of some of the accessory factors are still debated. Identification of conserved viral RNA processes, viral protein-host RNA interactions, and understanding how the virus hijacks these processes will enable the discovery of new antiviral targets and strategies.

Recent transcriptome-wide and proteome-wide studies in viral protein-host protein interactions^1^, viral protein and RNA interactions with host proteins^2, 3^, and viral RNA-host RNA interactions^4^ contribute to our understanding of host-virus interactions in SARS-CoV-2 infection. A recent study on the protein interactome with viral RNA shows that many of the SARS-CoV-2 proteins are RNA binding proteins that bind to its own RNA genome and mRNA transcripts^2^. As a (+ss)RNA virus, SARS-CoV-2 proteins are also found to associate with several host RNA binding proteins (RBPs)^1^, suggesting a possibility that SARS-CoV-2 proteins interact with the host transcriptome to a greater degree than previously anticipated. For example, several recent publications showed that NSP1 binds to the mRNA entry site on the host ribosomal RNA to inhibit host translation^5–7^. SARS-CoV-2 nucleocapsid protein interactome comprises many host RNA processing machinery proteins and stress granule proteins^1^, suggesting a potential role in interfering with host RNA processing and driving stress granule formation^8, 9^. One study showed that SARS-CoV-2 proteins bind to ∼140 host transcripts^10^. Focusing on non-coding RNAs, SARS-CoV-2 NSP16 was found to bind to U1/U2 snRNA to interfere with splicing, while NSP8 and NSP9 bind to the signal recognition ribonucleoprotein 7SK to block protein trafficking^10^. However, our knowledge of how viral and host coding RNAs interact with viral proteins and the functional implications of these interactions remain limited. Thus, a comprehensive interrogation of SARS-CoV-2 viral protein-RNA interactions is still needed for us to gain insights into viral RNA processing and how the virus hijacks host cellular machinery for its replication while simultaneously suppressing host gene expression.

Here, we investigated whether and how SARS-CoV-2 proteins, both in the context of authentic virus infection and exogenous expression, directly interact with the viral genome and host transcriptome using enhanced crosslinking and immunoprecipitation (eCLIP). More than 150 human RBPs have been profiled by eCLIP ^11, 12^ leading to insights into their regulatory roles^13^. Using this method, our systematic study of the RNA interactome of viral proteins infers new functionality in the largely unannotated viral proteome. Moreover, our results on the discovery of SARS-CoV-2 protein interactomes with host transcriptomes provides fundamental knowledge of host dependencies and viral mechanisms for hijacking the host cell, which will offer new opportunities to develop novel antiviral therapeutics.

## Results

### eCLIP elucidates SARS-CoV-2 protein-viral RNA interactions in virus infected cells

To investigate the RNA interactome of SARS-CoV-2 proteins, we performed eCLIP^11^ on SARS-CoV-2 infected African Green Monkey kidney (Vero E6) cells, which are an efficiently infected cell line **(Fig. 1a**). Infected cells were subject to UV irradiation, which covalently crosslinked interacting proteins to RNAs. This was followed by immunoprecipitation of non-structural proteins NSP8 and NSP12, which form part of the replication transcription complex (RTC), and N (nucleocapsid), using protein-specific antibodies to isolate the bound RNA. The RNA-bound proteins were resolved via SDS-PAGE and transferred to nitrocellulose membranes such that only the region spanning the expected protein size and 75 kDa larger were excised and purified in subsequent steps. The same size region of a non-immunoprecipitated input whole cell lysate was included as size-matched input to identify and remove non-specific, enriched sequences. RNA was converted to DNA libraries, sequenced to an average depth of ∼25 million reads, and mapped to the SARS-CoV-2 viral genome and African Green Monkey genome to determine SARS-CoV-2 protein RNA interactions. Thus, reads from the immunoprecipitation (IP) samples correspond to RNA crosslinked to the IP enriched proteins NSP8, NSP12, and N, while reads from the input (IN) samples correspond to RNA crosslinked to RBPs at a similar size to the IP protein in the cell milieu. Normalizing read density of IP to IN samples provide a measure of protein-specific RNA interaction.

**Figure 1.**
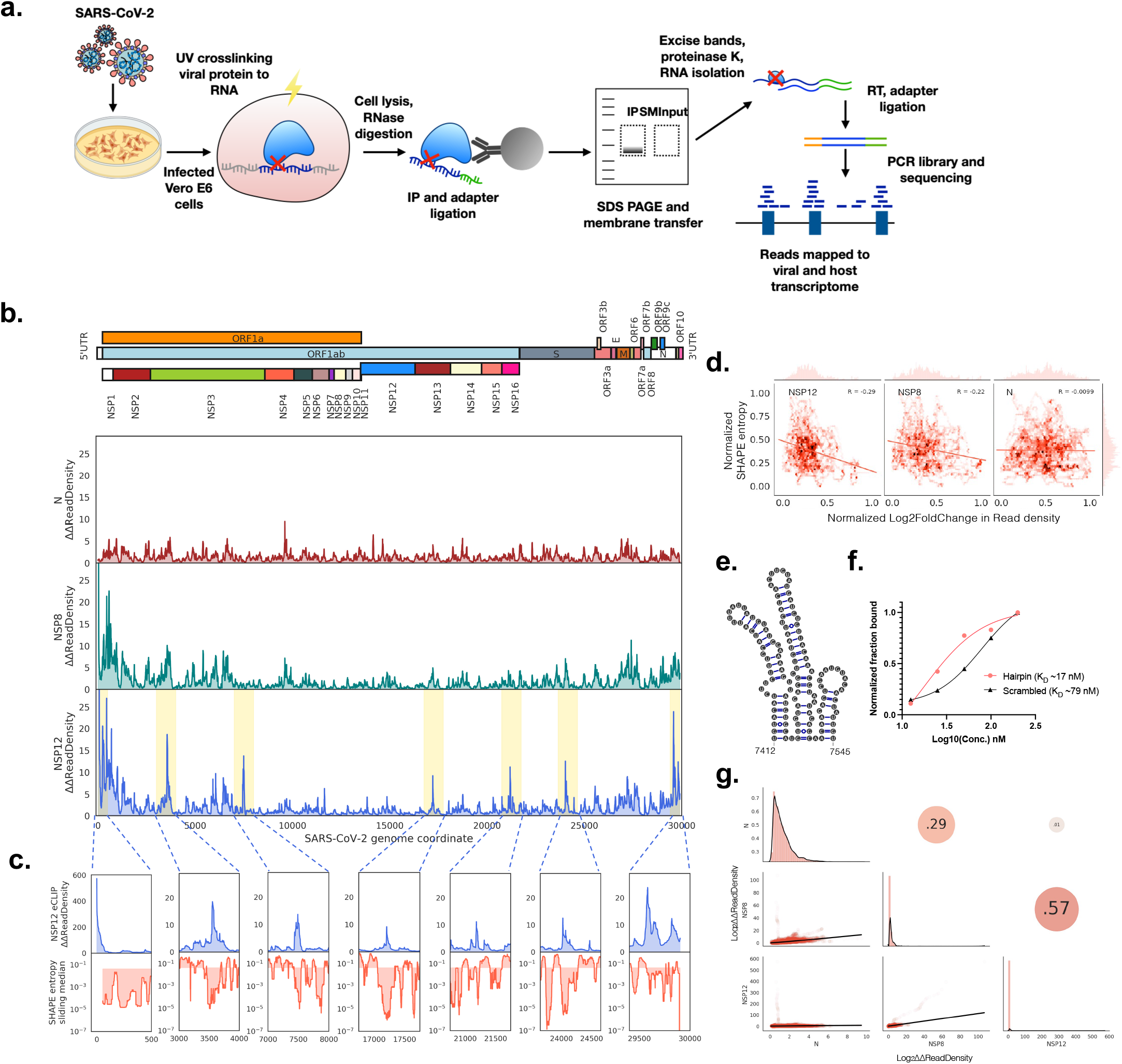
Genome maps of SARS-CoV-2 protein interactions with viral RNA. **a)** Schematic showing eCLIP performed on SARS-CoV-2 proteins in virus infected Vero E6 cells. Proteins in infected cells are UV crosslinked to bound transcripts, which are immunoprecipitated (IP) with antibodies that recognize NSP8 (primase), NSP12 (RNA dependent RNA polymerase, RdRp) and N (nucleocapsid) proteins. Protein-RNA IP product and Input lysate are resolved by SDS-PAGE and membrane transferred, followed by band excision at the estimated protein size to 75kDa above in both IP and Input lanes. Excised bands are subsequently purified, and library barcoded for Illumina sequencing. **b)** Mean fold change of eCLIP read density mapped to the positive sense SARS-CoV-2 genome in immunoprecipitated (IP) compared to input samples. Mean is taken from n = 2 independent biological samples. **c)** NSP12 eCLIP zoomed into yellow highlighted regions in b. Top row, NSP12 eCLIP; bottom row, SHAPE Shannon entropy^16^ with a sliding median of 55 nt. Shaded region in bottom row is partitioned at the global median entropy. **d)** Correlation between normalized SHAPE entropy^16^ and normalized log_2_(Fold Change) of IP over INPUT eCLIP read density i.e. eCLIP enrichment for NSP12 (left), NSP8 (middle) and N (right). R, Pearson’s coefficient. **e)** Secondary structure^16^ of the NSP12 eCLIP peak region from position 7412-7545. **f)** Fraction of RNA bound to NSP12 from filter binding assay, using hairpin RNA from position 7414-7555 and scrambled RNA as negative control. **g)** Correlation matrix of mean fold change of eCLIP read density: Bottom left panels, 2D density plots; diagonal, density plot corresponding to samples in bottom labels; top right panels, Pearson’s coefficient between samples.

Our eCLIP results provide the first genome-wide map of RNA interactions with viral proteins during an authentic SARS-CoV-2 infection. Reads are mapped across the entire genome in the positive sense for both IP and IN samples for all proteins, with greater than 96% coverage in all input and IP samples (**Supplementary Fig. 1a).** The near complete coverage implies that most of the viral RNA interacts with RNA binding proteins. Two biological replicates were performed for each protein, and read densities show strong replicate agreement in all samples (Pearson’s coefficient ≥0.87; **Supplementary Fig. 1b**). Input-normalized IP read densities also display strong replicate correlation (Pearson’s coefficient ≥0.88; **Supplementary Fig. 1c**).

To identify positions with particularly enriched RBP binding, we computed relative positional enrichment (ΔΔReadDensity) by dividing the fold change of read density of IP over IN at each position by the global median fold change for that sample (**Fig. 1b; Supplementary Fig. 1d**). We observed strong relative positional enrichment in NSP8 and NSP12 eCLIP read density at the 5′ end, at 573-fold and 103-fold at position 1 respectively, but only 0.75-fold for N (**Supplementary Fig. 1e-f**). High relative positional enrichment is also observed for the region before position 66, which marks the start of the leader transcription regulatory site (TRS), with >22-fold for NSP12, >4.9-fold for NSP8, and only >0.6-fold for N (**Supplementary Fig. 1d; Supplementary Table 1**). In both NSP12 and NSP8, there appear to be a drop in enrichment earlier, around position 33. This corresponds to the end of Stem Loop 1 (SL1)^4^ in the 5′ untranslated region (UTR), potentially implicating SL1 in recruiting, stabilizing or otherwise regulating the replicase proteins. At the 3′ end, NSP12 and NSP8 appear strongly enriched (>5-fold relative to global median) after the stop codon in N at position 29,533 up to the start of the S2M^4, 14^ structured region in the 3′ UTR at position 29,695 (**Supplementary Fig. 1f**). NSP12 and NSP8 continued to be strongly enriched again after the S2M region at position 29,809^4^. The lack of enrichment at the S2M structure may suggest that its function is unrelated to recruiting replicase proteins NSP12 and NSP8. Our eCLIP findings thus provide a map of the direct interaction between replicase proteins NSP8 and NSP12 with regions in the UTRs likely involved in regulating replication and transcription^15^.

We observed that only a small fraction of reads mapped to the negative sense strand in the input samples (0.00075 for N, 0.00076 for NSP12, 0.0043 for NSP8) and the IP samples for N (0.00046). In contrast, NSP12 and NSP8 IP samples enriched the fraction of negative sense reads to 0.067 and 0.039, about 100- and 10-fold from IN samples, respectively (**Supplementary Fig. 2a**). The negative sense strand coverage in IP samples was also lower in N (33%) than in NSP12 (80%) or NSP8 (58%) (**Supplementary Fig. 2b**). NSP12 and NSP8 IP reads are piled up in the 5′ and 3′ regions on the negative sense strand (**Supplementary Fig. 2c**), which is similar to the positive sense strand. The low fraction of reads in the input samples prevents further quantitative assessments. Nevertheless, the findings confirm the roles for NSP12 and NSP8, but not N, in transcribing negative sense RNA templates to generate mRNAs for translation, and the ability for N to selectively associate with positive sense genomic RNA over negative sense RNAs.

Besides regions around the 5′ and 3′ end, we observe several sharp peaks with high relative enrichment in read density (>5-fold, for a contiguous region of 10 nt or more) i.e. 22 peaks for NSP12, 37 peaks for NSP8, and 7 peaks for N (**Supplementary Table 1**). NSP12 eCLIP has especially strong peaks; five example major peaks are highlighted (**Fig. 1b-c**) at regions 3533-3635, 7436-7526, 17202-17222, 21177-21206, and 24018-24079 (**Fig. 1b**), which have maximum positional fold changes of 19, 14, 9.2, 11 and 13-fold (**Supplementary Table 1**), respectively. Intriguingly, when we lined up these regions with SHAPE reactivity data^16^, the peaks correspond to regions with low SHAPE Shannon entropy values, which represent regions that are rigid or structured (**Fig. 1c**). A closer inspection shows these stem-loop structures to have long, stable stems (**Supplementary Fig. 3a**). As no strong sequence motifs were observed, we hypothesized that structural elements in the SARS-CoV-2 genome likely facilitate protein-RNA interactions with NSP12. Recently, it was shown via RNA footprinting with SHAPE structure probing experiments that in addition to RNA-RNA base pairing, some nucleotides with low SHAPE reactivity may be due to direct hydrogen bond interactions with RNA binding proteins^16^. For NSP12 and NSP8, we further observe a slight negative correlation between log_2_ fold change in eCLIP read density and SHAPE entropy^16^, where the latter correlates inversely with structuredness. Our eCLIP findings can thus add a layer of functional information – interaction with NSP12 – to structural elements in SARS-CoV-2.

We noticed that the peak at position 7436-7526 is uniquely enriched in NSP12 eCLIP only. Located near the 3′ end of the gene encoding for NSP3, this peak overlaps with a structured region of 3 consecutive stem loops at region 7412-7545^16^. To validate the specific protein-RNA interaction, we performed a filter binding assay of NSP12 with *in vitro* transcribed RNA bearing the sequence in this region. We included a scrambled control with the same sequence composition but shuffled such that the structure is no longer preserved (**Supplementary Table 2**). The peak RNA shows a binding dissociation constant K_D_ of 17 nM, compared to a K_D_ of 79 nM by the scrambled control. The scrambled control likely represents the non-specific affinity of NSP12 for RNA (**Supplementary Fig. 3b**). The sequence and structure of the central hairpin, where the highest point of the peak fold change is located, are also highly conserved among related betacoronaviruses (**Supplementary Fig. 3c-d**). It also appears to be located ∼500 nt downstream of the steplike reduction in RNA reads extending from the 5′ side of the genome (**Supplementary Fig. 3e**), though the functional linkage between the two features is unknown.

Finally, we compared the relative eCLIP enrichment between NSP8, NSP12 and N to investigate any similarities in the positive sense RNA interaction between the three proteins. When comparing the relative positional enrichment, N and NSP12 show no correlation (R^2^ of 0.01), whereas N and NSP8 are slightly correlated (R^2^ of 0.27; **Fig. 1g**). As expected, NSP8 and NSP12 are the most highly correlated (R^2^ of 0.57), though there are still substantial differences, as 43% of the variation is unaccounted for by the other protein (**Fig. 1g**). However, when we transform the data logarithmically, we observe greater correlations among all three proteins (**Supplementary Fig. 4**). As logarithmic transformations shed light on signals at lower values, the greater correlations among the log-transformed data of different proteins may imply a greater similarity of more transient protein-RNA interactions. This invites future inquiry about the importance of transient protein-RNA interactions in the life cycle of SARS-CoV-2.

### SARS-CoV-2 proteins interact with host RNAs in virus-infected cells

As RNA virus infections have been shown to perturb host transcriptomes, such as via mRNA degradation^17^, mRNA export inhibition^18^, splicing interference^19^, 5′ cap stealing^20^, and other ways of host translation inhibition^21^, evidence for direct interaction between viral proteins host cell RNAs can shed light on the mechanism and function as a result of these interactions. Therefore, we investigated the extent to which NSP8, NSP12 and N interacted with host RNAs. Targeted transcripts were determined by having one or more peaks that meet the stringent IDR (irreproducible discovery rate^12^) threshold of overlapping peaks between two replicates for every protein, and satisfy statistical cutoffs of p<0.001, and more than 8-fold enrichment in the immunoprecipitated sample (IP) over the size-matched input sample. We found that NSP8, NSP12 and N interact with 457, 703 and 24 genes with 658, 1457 and 39 significant peaks, respectively (**Fig. 2a**). Interestingly, the number of RNA reads in Transcripts Per Kilobase Million (TPM) from both NSP8 and NSP12 immunoprecipitation (IP) samples were mapped more frequently to host transcripts than viral RNA **(Fig. 2b)**. Among the target genes, NSP12 and NSP8 shared 128 genes in common (18% of NSP12 targets, 26% of NSP8 targets), implying that NSP12 and NSP8 may interact with different host genes in both their individual and complexed states. In contrast, a majority of N immunoprecipitated RNA reads were mapped to viral RNA, consistent with its role in enclosing the viral genome during virion assembly^22^. The large number of peaks (2137 total) that map to the 1058 host genes further suggests a potential in perturbing host gene expression that may be required for viral replication.

**Figure 2.**
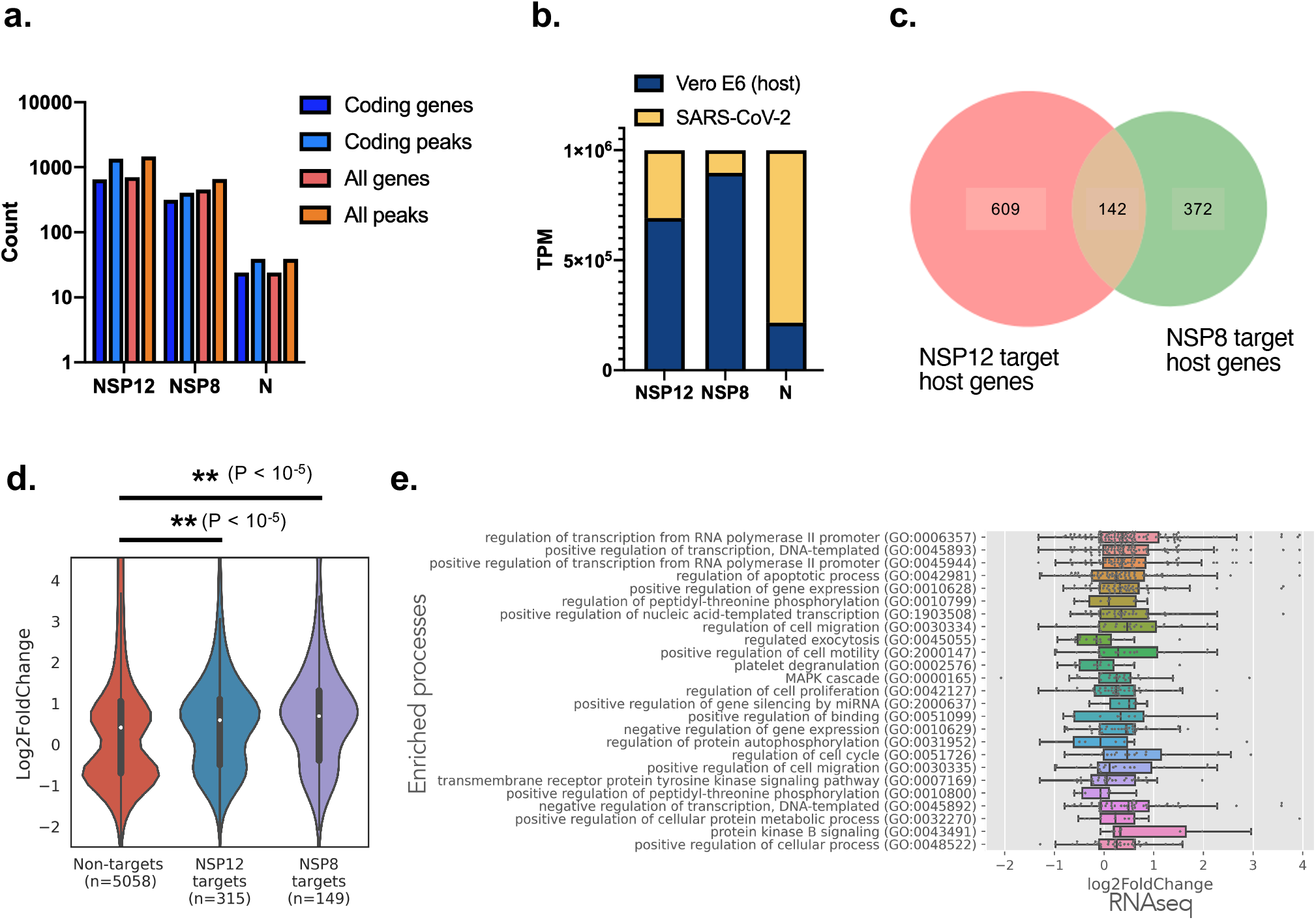
SARS-CoV-2 protein interactions with host cell RNAs in virus infected cells. **a)** Bar plot showing number of all genes, number of all peaks, number of coding genes and number of peaks mapping to coding genes from n = 2 biologically independent replicates of NSP12, NSP8 and N eCLIP of SARS-CoV-2 infected cells. Target genes have at least one reproducible peak (by IDR^12^) associated with each protein. **b)** Stacked bar plot showing TPM of reads mapped to the Vero E6 genome or SARS-CoV-2 genome in each of NSP12, NSP8 and N eCLIP. **c)** Venn diagram showing number of African Green Monkey (host) genes targeted by NSP8 and NSP12. **d)** Violin plot showing the distribution of Log_2_FoldChange in transcript levels in Vero E6 cells infected by SARS-CoV-2, for significantly differentially expressed genes (adjusted P < 0.05). Kolmogorov–Smirnov test p-values between eCLIP targets of NSP12 and NSP8 versus all differentially expressed genes are indicated above the plot. **e)** Top 25 Enriched Gene Ontology (GO) processes (adjusted p < 0.01) for NSP12 target host genes. Box plot indicates quartiles of differential expression (log_2_(Fold Change)) of target genes (grey dots).

To determine if there are differences in expression levels of the host target genes whose mRNAs are enriched in NSP12 and NSP8 eCLIP, we performed transcriptome-wide mRNA sequencing of SARS-CoV-2 infected Vero E6 cells and mapped the eCLIP target genes to differential expression levels. We found that NSP12 and NSP8 target mRNA levels are significantly increased than non-target genes (p<10^-5^, KS test; **Fig. 2d**). To understand the processes enriched by the target genes, we performed a Gene Ontology analysis. We found 54 GO processes that are significantly enriched by NSP12 target genes (P_adjusted_<0.01; **Supplementary Table 3**), whereas no significant GO processes are found in NSP8 target genes. Many of the GO processes fall into three broad categories related to regulating transcription and gene expression, cell cycle and apoptosis, and phosphorylation and signaling processes (**Fig. 2e**). Of the transcription regulation genes, many have antiviral response properties (e.g. *NF-κB, BATF, NR4A1, BMP2, SQSTM1, MAFF, MDM2*), while others have demonstrated proviral activities, such as *DDX5, SFPQ, FBXW11 and ATF-3*. Of the genes regulating cell proliferation, cell cycle and apoptosis, *PAK2* has been associated with anti-apoptotic signaling and promoting HIV survival^23^, whereas many other genes have overlapping annotation as the transcription regulation genes. A recent study elucidated global phosphorylation changes in cellular proteins upon SARS-CoV-2 infection^24^. Specifically, the p38/MAPK cascade activity is induced by viral infection, and treatment with p38 inhibitors has restrictive effects on viral proliferation.In our NSP12 eCLIP data, we also saw enrichment of the MAPK cascade and other signaling pathway genes (e.g. *MAPK1, MAP2K1/3, MAP4K4/5, PIM3, PAK2, EPHA2*).In the context of the whole virus infection where a multitude of viral proteins and host defense responses are at play, we cannot definitively conclude that the interactions between NSP12 and these mRNAs have a causative or inhibitive relationship. Nevertheless, the correlation of NSP12 protein-RNA interactions with these pathway genes, which are relevant to viral infection and host response, leads us to hypothesize a potential, albeit unknown, role, and our data represents a rich resource for subsequent mechanism studies. To understand the individual contribution of viral protein-host RNA interactions, we proceeded to profile the protein-RNA interactions of each SARS-CoV-2 protein.

### Exogeneously expressed SARS-CoV-2 proteins interact with one third of the transcriptome in lung epithelial cells

Even though we have performed NSP12, NSP8, and N eCLIPs in virus infected vero cells, in order to further investigate whether SARS-COV-2 proteins directly interact with the human host transcriptome, we performed eCLIP on the 29 proteins encoded in the SARS-CoV-2 genome and one mutant **(Fig. 2a)**. Due to the lack of antibodies specific for most of the viral proteins, the individual proteins were exogenously expressed in a lung epithelial cell line BEAS-2B, which is an immortalized primary bronchial cell line representative of normal lung physiology. Each protein was either fused with a 2xStrep tag and expressed stably via lentiviral transduction or fused with a 3xFLAG tag and expressed transiently via transfection. Following UV crosslinking, the tagged proteins were immunoprecipitated using anti-FLAG or anti-Strep antibodies. Subsequent RNA purification and library purification steps were performed as in the viral eCLIP experiments. Cells expressing only the 3xFLAG or 2xStrep tags and wildtype cells are used as controls to remove background peaks in subsequent analysis steps.

From our SARS-CoV-2 proteome-wide eCLIP results, SARS-CoV-2 proteins interacted with RNA represented by 4773 coding genes, which is about a third of the transcriptome of BEAS-2B cells. Nucleocapsid and non-structural proteins NSP2, NSP3, NSP5, NSP9 and NSP12 were found to target the greatest number of unique genes at 1339, 1647, 1199, 902, 863, and 865, respectively **(Fig. 2b)**. The large number of genes targeted by the viral proteins is consistent with the non-structural proteins from the replicase (ORF1ab) having a high affinity for its own RNA, though their potential for widespread interaction with host RNA has not been shown previously. The widespread interaction of Nucleocapsid with host RNAs when expressed in isolation is consistent with its capacity for nonspecific RNA binding, whereas it’s targeting the virus genome during RNA assembly occurs via interaction with the M protein^25^. For comparison, the extensively studied splicing factor RBFOX2 binds to 958 genes in HepG2 cells and 471 genes in K562 cells, the stress granule assembly factor G3BP1 binds to 561 genes in HepG2 cells, and the histone RNA hairpin-binding protein SLBP binds to 19 genes in K562 **(Fig. 3b)**. This suggests that viral proteins have similar capacities for interacting with RNA as endogenous human RBPs. Target genes with at least one significant eCLIP peak also appear highly distinct across the different SARS-CoV-2 proteins (**Fig. 3c**). Within individual targets, eCLIP reads also display different profiles, example include N eCLIP peak in 3′ UTR of *CXCL1*, NSP3 peaks found across all exons in *DYNCH1*, a NSP12 peak in 5′ UTR of *TUSC3* and a NSP2 peak in the intronic region upstream of 3′ splice site of *NAP1L4* (**Fig. 3d**). To cross validate the eCLIP findings that SARS-CoV-2 proteins interact with host cell RNAs, we validated a subset of these proteins as RBPs using crosslinking and solid phase purification (CLASP^26^), which stringently captures crosslinked protein-RNA interactions due to denaturing wash conditions. HEK293T cells transiently expressing NSP1, NSP2, NSP12, and ORF9c followed by pulldown of total RNA showed enrichment of these proteins (**Fig. 3e**). For comparison, we also included host RNA binding proteins ELAVL1, YTHDC1 and GAPDH as positive controls, and tubulin as negative controls. Furthermore, we performed RNA interactome capture (RIC^27^) of poly-A RNA (mRNA, lincRNA, and other POLII transcripts) pulldown using an oligo(dT) primer and found that NSP2 and NSP12 were enriched, but not NSP1 (**Fig. 3f**), which mostly enriched ribosomal RNAs in eCLIP (**Supplementary Fig. 5a-b**).

**Figure 3.**
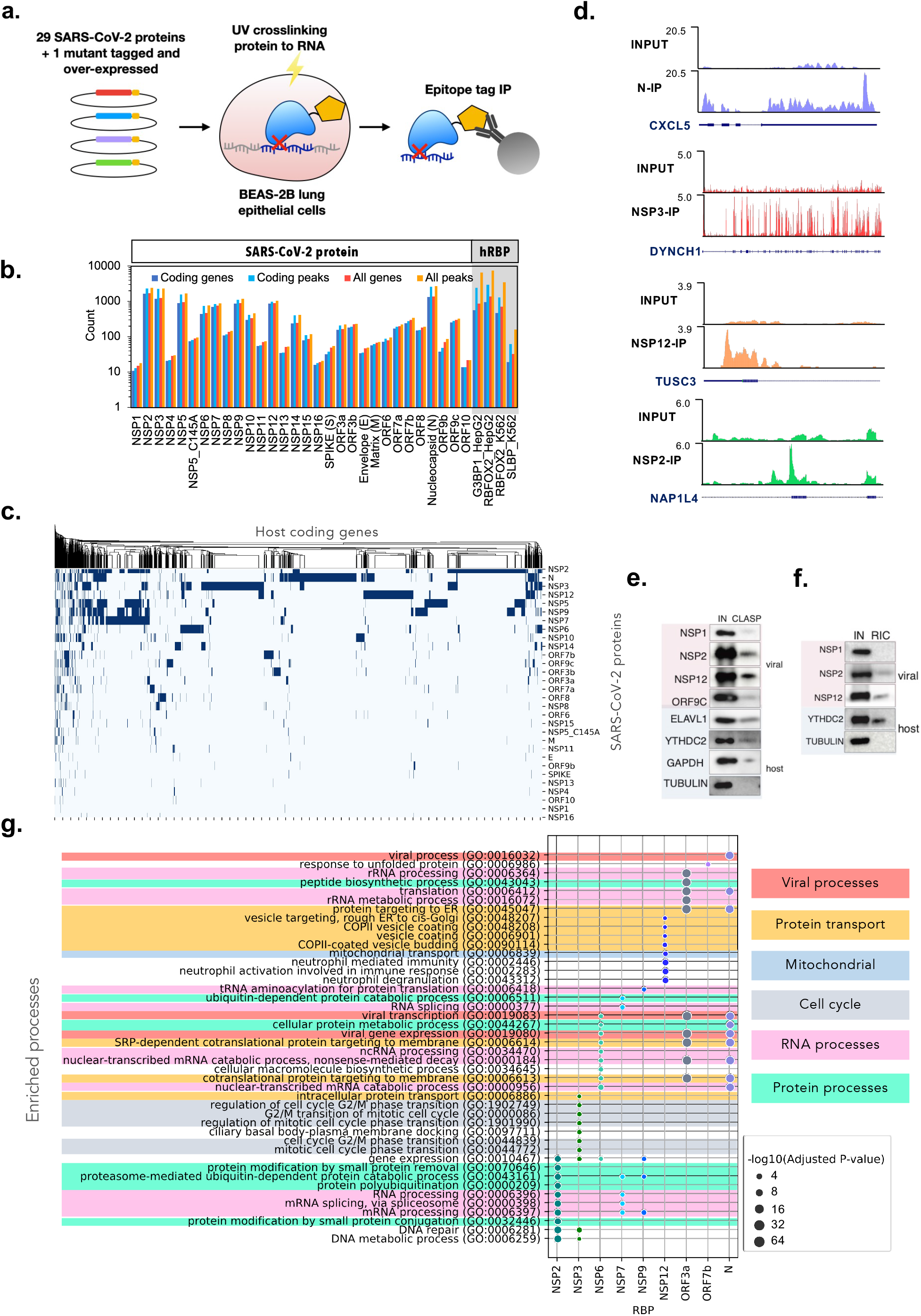
The SARS-CoV-2 proteome interacts with thousands of host transcripts. **a)** Schematic showing SARS-CoV-2 proteins individually tagged and expressed in human lung epithelial cells BEAS-2B to assay with eCLIP. **b)** Bar plot indicating number of all genes, number of all peaks, number of coding genes and number of coding peaks found to interact with each protein from n = 2 biologically independent experiments. In addition to SARS-CoV-2 proteins, ENCODE eCLIP data for example human RNA-binding proteins (hRBPs) are included for comparison. Target genes have at least one reproducible peak (by IDR) associated with each protein. **c)** Clustermap showing unique host coding genes (columns) targeted by each SARS-CoV-2 protein (rows). **d)** Example genome browser tracks for NSP3, NSP12, N and NSP2 mapping to DYNCH1, TUSC3, CXCL5 and NAP1L4 respectively. **e & f)** Western blots showing viral (pink background) and human (blue background) proteins enriched via CLASP (e) and RIC (f), with total cell lysate showed in input column (IN). **g)** Enriched Gene Ontology (GO) processes (adjusted p-value < 10-5) of unique eCLIP target coding genes for various SARS-CoV-2 proteins.

Distinct processes related to viral replication and host response are targeted by the viral proteins as shown by gene ontology (GO) analysis **(Fig. 3g, Supplementary Fig. 6)**. Many of the enriched GO terms are related to nucleic acid and protein synthesis, modification and transport, which is consistent with the primary objective of the virus hijacking host resources for its own biosynthesis and replication. Notably, several protein transport processes are enriched, namely SRP-dependent protein targeting to membrane as enriched by NSP6, ORF3a and N, and COPII vesicle budding and targeting from rough ER to Golgi as enriched by NSP12. These may be involved in viral vesicle formation to serve as replication organelles, as found in a number of positive sense RNA viruses^28^. Immune response processes are also enriched, including neutrophil mediated immunity targeted by NSP12 and platelet degranulation targeted by ORF9c. This supports our choice of lung epithelial cells as a model system that express the relevant cytokines for recruiting immune cells^29^. While the enriched GO terms are highly relevant to viral and host response processes, further analysis of binding patterns is needed to determine if there are any functional implications of viral proteins interacting with these genes.

To determine if there are sequence features that the viral proteins recognize, we generated sequence logos from 6-mers of eCLIP peaks. While some of the proteins display strong sequence preferences, most proteins appear to interact more non-specifically **(Supplementary Fig. 7)**. Some motifs resemble enrichments observed for human RBPs, where M, ORF7a and NSP10 appear to favor G-rich or GU rich motifs, and NSP5 has a motif (GNAUG). Other motifs may result from regional binding preferences **(Fig. 2e)**, as NSP2 and NSP9 have a strong preference for UC-rich polypyrimidine motifs (p values of 10^-96^ and 10^-41^ respectively), which may be a result of their binding to polypyrimidine tracts in intronic regions (discussed later), whereas N has an AU-rich motif likely because it preferentially binds to 3′ UTR which contain AU-rich elements^30^. NSP3, a large multifunctional protein, appears to coat entire exons and may not have a meaningful sequence motif. NSP12 primarily binds in the 5′ UTR, and a weakly enriched GUCCCG motif that resembles terminal oligopyrimidine (TOP) motifs^31^ hints at a possible role in translation perturbation.

Our systematic interrogation of SARS-CoV-2 protein-host RNA interactions demonstrates that a majority of SARS-CoV-2 viral proteins are RNA binding proteins that target roughly a third of the human transcriptome. Our analysis implies that these viral proteins may be involved in perturbing many essential cellular processes of the host. As eCLIP in virus infected cells are limited by IP-grade antibodies, we focus on the data obtained from the exogenous expression of individual proteins in BEAS-2B cells for systematic analysis of potential functional implications.

### SARS-CoV-2 proteins upregulate protein expression of target transcripts

By examining the regional binding preferences of each SARS-CoV-2 protein, we found that SARS-CoV-2 proteins are enriched at distinct regions of target mRNAs, which imply different regulatory functions because of the protein-RNA interaction. Aggregating the analysis of all targeted peaks for each SARS-CoV-2 protein identifies RNA regions that are preferentially bound (**Fig. 4a**). Of note, NSP12, ORF3b, ORF7b and ORF9c show the highest proportion of peaks in the 5′ UTR; NSP2, NSP3, NSP6 and NSP14 show the highest proportion of peaks in the coding region (CDS), NSP5, NSP7 and NSP9 display a high proportion of peaks in intronic regions, and N and NSP15 show the largest proportion of peaks in the 3′ UTR. We also performed a metagene analysis of read density across all target mRNA transcripts, where each of the 5′ UTR, CDS and 3′ UTR regions in an mRNA are scaled to standardized lengths (**Fig. 4b**). We found that even though NSP2 has a similar number and proportion of peaks in the CDS as NSP3, it mainly targets the region spanning the 5′ UTR and coding start. In contrast, NSP3 reads, along with that of NSP6 and NSP14, coat the entire CDS, with a slight bias towards the start of the coding sequence.

**Figure 4.**
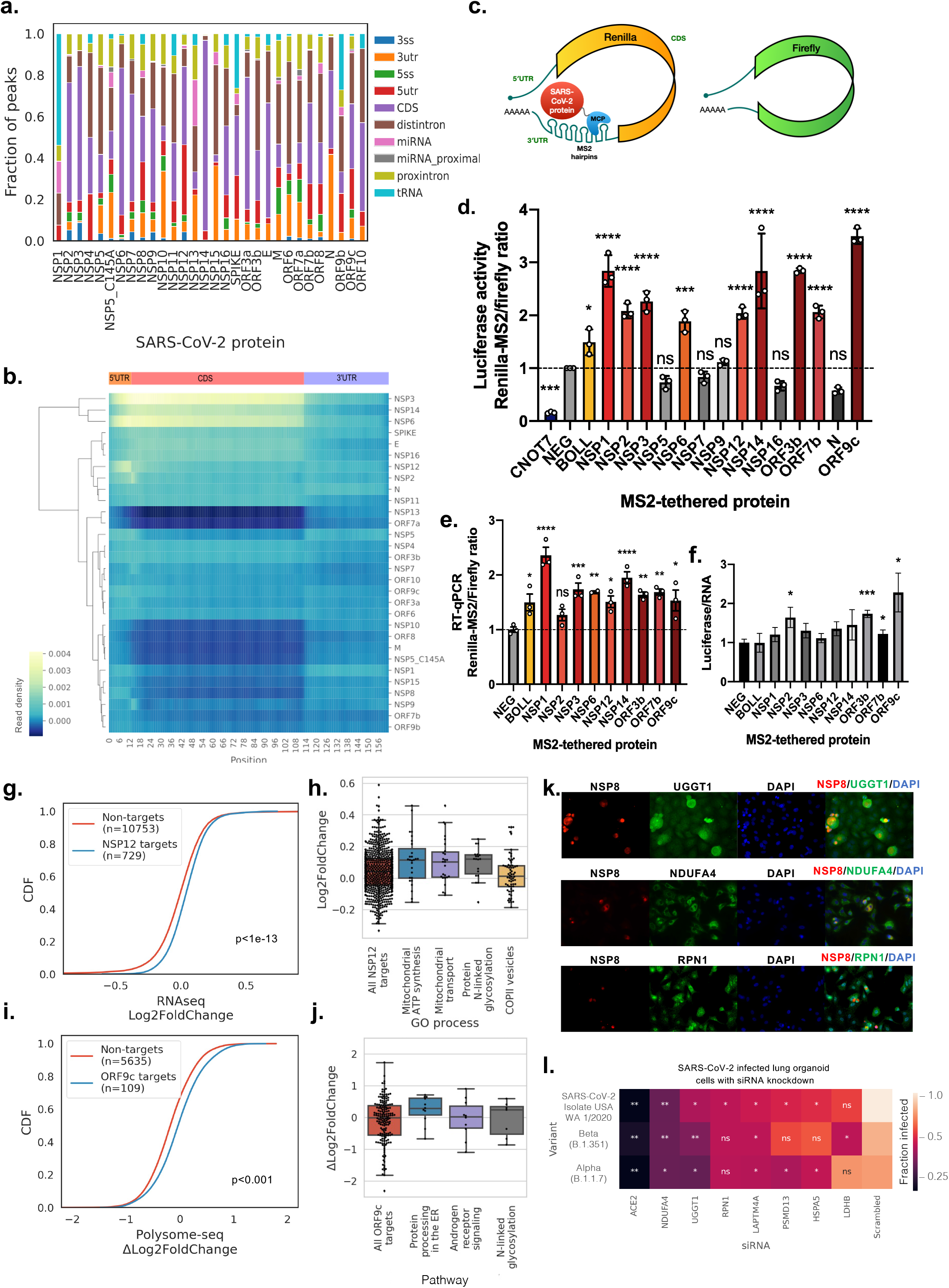
SARS-CoV-2 proteins specifically upregulate target gene expression. **a)** Stacked bar plot showing fraction of reproducible peaks (by IDR14) mapping to different regions of coding genes. 3ss, 3′ splice site; 3utr, 3′ untranslated region (UTR), 5ss, 5′ splice site; 5utr, 5′ UTR; CDS, coding sequence. **b)** Clustermap showing read density of target RNA by each SARS-CoV-2 protein scaled to a metagene profile containing 5′ UTR, CDS and 3′ UTR regions. **c)** Schematic showing the Renilla-MS2 and Firefly dual luciferase reporter constructs, where individual SARS-CoV-2 proteins fused to MCP are recruited to the Renillia-MS2 mRNA. **d & e)** Bar plot showing luciferase reporter activity ratios (d) and reporter RT-qPCR ratios (e) for the indicated coexpressed SARS-CoV-2 protein, known human regulators of RNA stability (CNOT7, BOLL) and negative control (FLAG peptide). Ratios are normalized to the negative control (mean ± s.e.m., n = 3 biologically independent replicate transfections; * p<0.05, ** p<0.005, *** p<0.0005, **** p<0.0001, two-tailed multiple t-test; ns, not significant). **f)** Bar plot showing the fold change of luciferase activity ratio and RT-qPCR ratio (mean ± s.e.m, n = 3; * p<0.05, *** p<0.001, two-tailed Welch’s t-test). **g)** Cumulative distributive plot (CDF) of log_2_(Fold Change) of gene expression in HEK293T cells transfected with a plasmid overexpressing NSP12 versus an empty vector plasmid. KS test p values indicate significance of difference in differential expression of NSP12 target genes versus non-eCLIP target genes. **h)** Enriched Gene Ontology (GO) processes (adjusted p < 10-4) of NSP12 target genes, with box plots indicating quartiles of differential expression (log_2_(Fold Change)) of target genes (black dots). **i)** CDF plot of Δlog_2_(Fold Change) of polysomal mRNA levels in BEAS-2B cells nucleofected with a plasmid overexpressing ORF9c versus an empty vector plasmid. KS test p values indicate significance of difference in differential expression of ORF9c target genes versus non-eCLIP target genes. **j)** Enriched BioPlanet pathways (adjusted p < 0.01) of ORF9c target genes, with box plots indicating quartiles of differential expression (Δlog_2_(Fold Change) of polysomal mRNA levels) of target genes (black dots). **k)** Immunofluorescence images (40X) of SARS-CoV-2 infected A549-ACE2 cells stained for SARS-CoV-2 NSP8 (red), endogenous genes (green), DNA content (blue). **l)** Heat map showing infection rate as measured by the integrated intensity of immunofluorescence staining of SARS-CoV-2 nucleocapsid protein in human iPSC derived lung organoid cells. Cells are treated with siRNAs targeting different host genes prior to viral infection by three different variants of SARS-CoV-2. Significant differences in infection rates are given by two-tailed t-test, * p<0.05, **p<0.01, ns, not significant, as compared to scrambled siRNA control for n = 3 biologically independent samples.

Since 8 of the SARS-CoV-2 proteins – NSP2, NSP3, NSP6, NSP12, NSP14, ORF3b, ORF7b and ORF9c – have binding preferences at the 5′ UTR and CDS, we hypothesized that their protein-RNA interactions could affect expression of the target mRNAs at the level of RNA turnover or translation. To evaluate the functional role of the specific protein-RNA interactions of SARS-CoV-2 proteins and target transcripts, we characterized 14 of the proteins with the highest number of unique target coding genes using our recently published tethered function reporter assays^32^ (**Fig. 4c**). We fused individual proteins with an MS2 phage coat protein (MCP), which localizes the tagged protein to MS2 aptamer hairpins inserted in the 3′ UTR of *Renilla* luciferase. A firefly luciferase without MS2 hairpins is included as a control for non-specific effects of the viral protein. Plasmids encoding the MCP-tagged proteins and reporter constructs are co-transfected into HEK293T cells. Changes in *Renilla* luciferase activity normalized to firefly luciferase activity measures up- or downregulation of protein expression via either translation or mRNA stability because of positioning the MCP tagged protein in the vicinity of the *Renilla* mRNA. The luciferase readout does not by itself distinguish between translational or mRNA stabilizing effects.

From our tethering experiments, we found that the ratio of Renilla-MS2 to firefly luciferase for 9 of the 14 SARS-CoV-2 proteins increase 1.9 (NSP6) to 3.5-fold (ORF9c) relative to FLAG-MCP control (p-value < 0.002, two tailed multiple *t*-test) (**Fig. 4d**). Interestingly, these SARS-CoV-2 proteins raise the target luciferase activity to greater extent than the tethering of BOLL (1.5-fold), which is a human RBP previously characterized to be amongst the strongest up-regulators from a screen of more than 700 human RBPs^32^. Even though NSP1 eCLIP enriched very few host mRNAs and its peaks are not mapped to the 5′ UTR and CDS, our results for NSP1 are consistent with its ability to enhance the translation of its own mRNA via interacting with the 5′ UTR of the genomic viral mRNA^6^. Of the remaining 5 SARS-CoV-2 proteins, only NSP5, NSP16 and N display slight (but not significant) down-regulation effects (0.73-fold to 0.58-fold) compared to the FLAG peptide control, but to a lesser extent than that of the known translation repressor CNOT7 (0.16-fold). NSP7 and NSP9 appear to have no effect on the relative luciferase activity of the target *Renilla* reporter. To understand if the increase in luciferase activity is occurring at the RNA or protein level, we performed RT-qPCR to measure the ratio of Renilla-MS2 to Firefly mRNAs. For all the proteins except for NSP2, the Renilla-MS2/Firefly mRNA ratio is significantly increased (p<0.05) compared to wildtype, albeit to different extents for different proteins (**Fig. 4e**). Of note, ORF9c shows the greatest enhancing effect (3.5-fold) in the dual luciferase assay, but its effect on the reporter RNAs is middling (1.5-fold). However, ORF9c displays the greatest fold change in luciferase activity ratio to RNA ratio (2.3-fold) **(Fig. 4f**), followed by NSP2 and ORF3b (1.6 and 1.7 fold respectively). The rest of the proteins range from 1.1-fold (NSP6) to 1.5-fold (NSP14), compared to 1.0-fold of BOLL, suggesting that the increase in abundance of the targeted reporter likely occurs at both the RNA and protein levels.

Based on the results of our reporter assay, we hypothesize that SARS-CoV-2 proteins that interact with the 5′ UTR and CDS of target genes could increase their abundance. Since NSP12 demonstrated targeted increase of reporter mRNA levels, we transiently overexpressed NSP12 and performed mRNA sequencing to determine if there are transcriptome-wide changes in gene expression. By comparing HEK293T cells transfected with NSP12 versus cells transfected with an empty plasmid vector, we observed that the eCLIP targets of NSP12 have greater log_2_ fold changes in mRNA levels than genes that are not eCLIP targets of any SARS-CoV-2 protein (p <10^-13^, KS test; **Fig. 4g**). Genes in the enriched GO processes, such as mitochondrial ATP synthesis and transport, protein N-linked glycosylation and COP II vesicle budding, are similarly upregulated by the overexpression of NSP12 (**Fig. 4h**). These observations provide support for the hypothesis that NSP12 has the capacity to increase the abundance of its eCLIP target RNAs.

To determine if SARS-CoV-2 proteins enhance the translation of endogenous genes, we performed polysome profiling on ORF9c, as it demonstrated the greatest ratio of changes in luciferase activity to changes in luciferase mRNA levels. We first determined the log_2_ fold changes of polysomal mRNA levels versus total mRNA levels in BEAS-2B cells overexpressing ORF9c and then compared it to wildtype BEAS-2B cells to obtain differential translation rates Δlog_2_FoldChange. We observed that the eCLIP targets of ORF9c have higher Δlog_2_FoldChange in translation rates than genes that are not eCLIP targets of any SARS-CoV-2 protein (p <10^-3^, KS test; **Fig. 4i**). Genes in the enriched pathways, such as protein processing in the ER, androgen receptor signaling, and protein N-linked glycosylation are similarly upregulated by the overexpression of ORF9c (**Fig. 4j**). Among the N-linked glycosylated GO term genes, Ribophorin I (RPN1) is part of an N-oligosaccharyl transferase complex that links high mannose oligosaccharides to asparagine residues found in the Asn-X-Ser/Thr consensus motif of nascent polypeptide chains, and UDP-Glucose Glycoprotein Glucosyltransferase 1 (UGGT1) is a soluble protein of the endoplasmic reticulum (ER) that selectively glucosylates unfolded glycoproteins. Represented in the mitochondrial ATP synthesis coupled electron transport and the respiratory electron transport chain GO processes, NDUFA4 is part of the enzyme cytochrome-c oxidase (or complex IV) and is important for its activity and biogenesis^33^. Consistent with our data showing that exogenous expression of ORF9c can interact with *RPNI, UGGT1 and NDUFA4* RNA and increase protein expression, we found that SARS-CoV-2 infection increases RPNI, UGGT1 and NDUFA4 protein levels specifically in infected cells (**Fig. 4h, i; Supplementary Fig. 8a**).

To determine if some of these host RNAs that interact with expression enhancing SARS-CoV-2 proteins are pro-viral or antiviral, we next investigated the impact of siRNA knockdown of these genes on viral infection or replication in human lung organoid cells. Human lung organoids are a physiologically relevant system to study infections and have been shown to be highly infectible by SARS-CoV-2^34^. siRNAs were selected from the target mRNAs of SARS-CoV-2 proteins with mRNA stabilization or translation enhancing activities, in addition to an anti-*ACE2* siRNA and a scrambled sequence as a negative control. We assayed for infected cells by immunofluorescence and determined infection rate by measuring the total integrated fluorescence intensity of the stained nucleocapsid protein. To control for cell viability, we divided the integrated intensity to the area stained by DAPI, and normalized the values to the scrambled control (**Fig. 4l**). We found that siRNA knockdown of *RPN1, UGGT1, NDUFA4, HSPA5, PSMD13, LAPTM4A, LAMP1,* and *LDHB* significantly (p<0.05, two-tailed *t*-test) reduced infection rates for at least one of the tested SARS-CoV-2 variants compared to a scrambled siRNA control (**Fig. 4l**). Of note, siRNA knockdown of *NDUFA4, UGGT1 and LAPTM4A* significantly reduced viral infection in all three variants.

Taken together, these results suggest that SARS-CoV-2 proteins with a preference for binding to 5′ UTR and CDS regions have a capacity for increasing the abundance of target mRNAs and/or translation rates. Furthermore, we found that eCLIP target genes are associated with enhanced RNA levels via NSP12 overexpression, and increased translation rates with ORF9c overexpression.

### NSP9 associates with the nuclear pore to block mRNA export

Since it was recently reported that several SARS-CoV-2 proteins are localized to the cell nucleus^35^, we were curious to find that the eCLIP peaks of NSP2, NSP5, NSP7, and NSP9 are enriched in intronic regions (**Fig. 5a**). To test whether these targets are implicated in infection induced alternative splicing, we performed deep sequencing (>50 million 100 nt reads per sample) of SARS-CoV-2 infected A549-ACE2 cells. We found a total of 1839 alternatively spliced genes across all five types of alternative splicing events i.e. alternative 5′ and 3′ splice site, skipped exons, skipped introns and mutually exclusive exons (false discovery rate < 0.1, |Inclusion level difference| > 0.05). By comparing genes with eCLIP peaks mapped to intronic regions or splice sites to genes not targeted by any of the SARS-CoV-2 proteins, we observed no significant differences in alternative splicing (significance level α = 0.01, KS test; **Supplementary Fig. 9a**). The little or no relationship with splicing led us to consider other potential ways intronic binding by these SARS-CoV-2 proteins may be affecting the host transcriptome.

**Figure 5.**
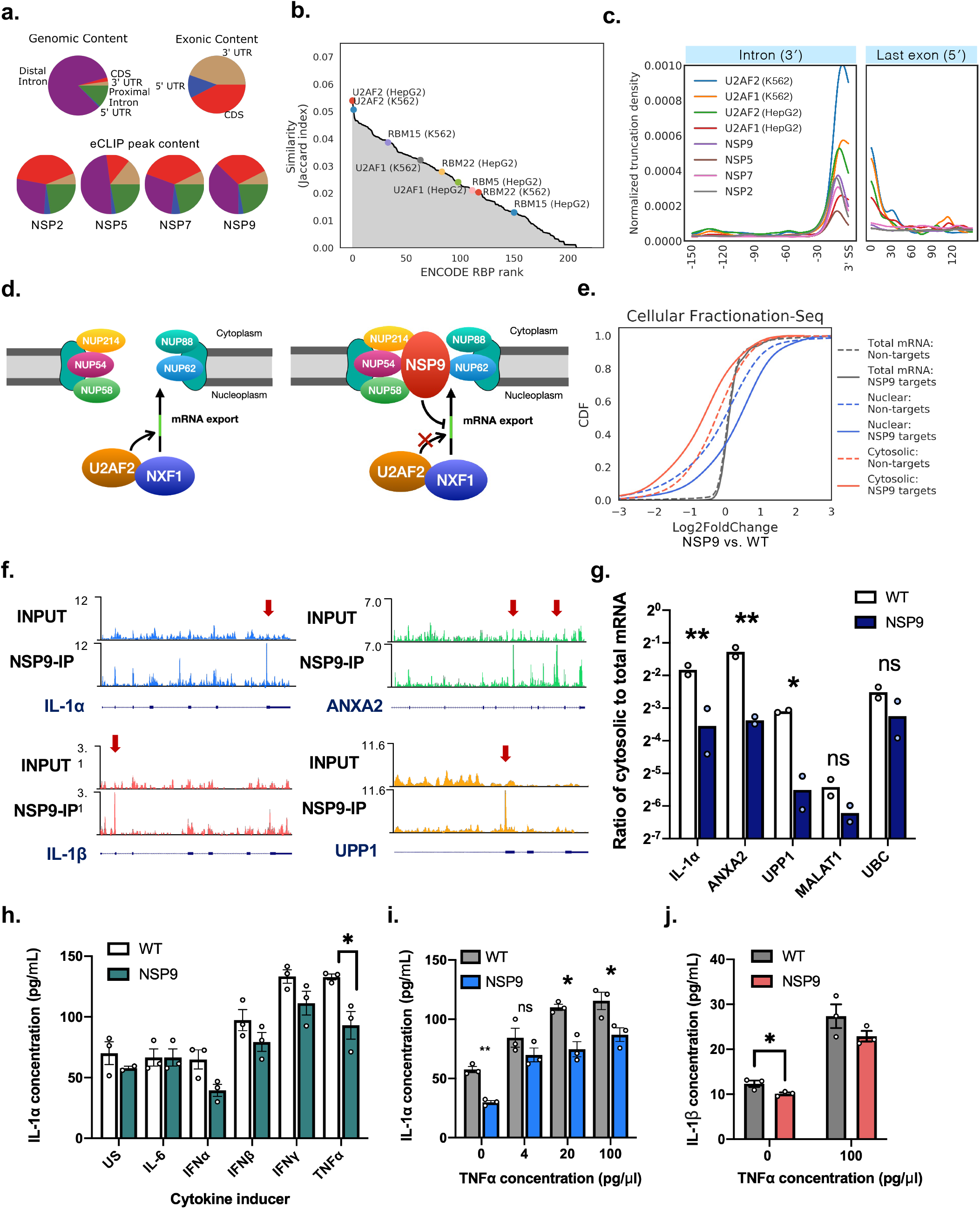
NSP9 interacts with U2AF2 substrates and inhibits mRNA export. **a)** Pie charts showing distribution of eCLIP peaks across different coding RNA regions for NSP2, NSP5, NSP7 and NSP9. Genomic content and exonic content are based on the hg19 human reference genome. **b)** Jaccard index similarity of NSP9 target genes as compared with all 223 ENCODE RBP datasets. **c)** Metadensity of eCLIP reads truncation sites averaged across all RNA targets by SARS-CoV-2 NSP2, NSP5, NSP7 and NSP9, and U2AF1/2 from the ENCODE consortium, zoomed into the region 150 nt upstream of 3′ splice sites, and the region 150 nt downstream of the 5′ end of the last exon. **d)** Schematic illustrating a model of NSP9 interacting with nuclear pore complex proteins NUP62, NUP214, NUP58, NUP88 and NUP54, and inhibiting U2AF2 substrate recognition in preventing NXF1 facilitated transport. **e)** Cumulative distributive plot (CDF) of log_2_(Fold Change) of BEAS-2B cells overexpressing NSP9 versus wildtype BEAS-2B cells in each of nuclear, cytosolic, and total mRNA fractions. Solid line indicate NSP9 target genes, dashed lines indicate genes that are not NSP9 targets. **f)** Genome browser tracks of NSP9 eCLIP target RNA mapped to IL-1α, IL-1β, ANXA2 and UPP1. g) Bar plot showing ratios of cytosolic to total fraction of mRNA levels measured by RT-qPCR, in wild type (WT) BEAS-2B cells, and BEAS-2B cells transduced to express NSP9 (*p<0.05, **p<0.0005, two-tailed multiple t-test with pooled variance, n = 2 biologically independent replicates). **h)** Bar plot showing mean concentration of IL-1α in culture media from WT and NSP9 expressing BEAS-2B cells, 48h after induction by cytokines indicated on the x-axis (US, unstimulated; mean ± s.e.m, n = 3 biologically independent replicates; *p < 0.05, Tukey’s multiple comparisons test). **i)** Bar plot showing mean concentration of IL-1α in culture media from WT and NSP9 expressing BEAS-2B cells, 48h after induction by different levels of TNFα (mean ± s.e.m, n = 3 biologically independent replicates, *p<0.05, **p<0.005, two-tailed t-test). **j)** Bar plot showing mean concentration of IL-1β in culture media from WT and NSP9 expressing BEAS-2B cells, 48h after induction by 0 or 100 ng/ml TNFα (mean ± s.e.m, n = 3 biologically independent replicates, *p<0.05, **p<0.005, two-tailed t-test).

To infer molecular function, we next compared the similarity between each of 223 ENCODE RBP datasets with NSP2, NSP5, NSP7, and NSP9 by computing the Jaccard Index of target genes. We found that U2AF2’s target gene set is most similar to NSP2, NSP7, and NSP9, and ranks highly for NSP5 (**Fig. 5b**; **Supplementary Fig. 9b**). However, since the cell lines used for ENCODE – HepG2 and K562 – and in this study – BEAS-2B – are different, the Jaccard indexes are low, at 0.050 for NSP7, 0.054 for NSP9, 0.057 for NSP5, and 0.074 for NSP2. To further ascertain similarity with U2AF2 protein-RNA interactions, we inspected the positional read density. The 5′ end of each eCLIP read can be used to approximate the crosslink site where reverse transcription is aborted or truncated when converting protein-bound RNA to cDNA. By taking the mean of the 5′ read truncation density across all target genes, we observe a strong enrichment for the truncation site at a median of 11 nt upstream of the 3′ splice site (**Fig. 5c; Supplementary Fig. 9c**). Furthermore, we observed a strong overlap between U2AF2 eCLIP performed in both HepG2 and K562 cells with NSP2, NSP5, NSP7, and NSP9, with median truncation site at 10 nt upstream of 3′ splice site, providing evidence of substrate similarity. Using affinity mass-spectrometry, a recent publication showed that NSP9 interacts with several nuclear pore complex proteins, including NUP62, NUP214, NUP88, NUP54 and NUP58^1^ (**Fig. 5d**). We confirmed that NUP62 indeed co-immunoprecipitated with NSP9 (**Supplementary Fig. 9d-e**). Even though U2AF2 was not found in the protein-protein interaction network of NSP9, it was previously reported to facilitate the binding of nuclear export factor TAP/NXF1 to its mRNA substrates^36^. From these observations, we hypothesize that NSP9 may interfere with mRNA export by associating with the nuclear pore and interfering with the U2AF/NXF1 complex for RNA substrate recognition (**Fig. 5e**). The significance of NSP2, NSP5, and NSP7 association with this intronic region may be less clear, and will benefit from future studies for clarification.

To determine if NSP9 inhibits mRNA export activity, we assayed for the mRNA levels of NSP9 target genes in cytosolic and nuclear fractions. Both NSP9 expressing BEAS-2B cells and the wild type BEAS-2B cells were fractionated, followed by RNA extraction and preparation for mRNA sequencing. We observed no difference in log_2_ fold changes of mRNA levels in NSP9 overexpressing cells versus wildtype cells between NSP9 eCLIP targets and non-targets, which agrees with the observation of lack of perturbation of target gene expression in the dual reporter assay. However, NSP9 eCLIP targets displayed greater log_2_ fold change of mRNA levels in the nuclear fraction and lower levels in the cytosolic fraction than non-target genes (**Fig. 5e**). To validate the sequencing results, we performed individual RT-qPCR on the subcellular fractionated RNAs for individual target mRNAs *IL-1α*, *ANXA2* and *UPP1* (**Fig. 5f**), and observed lower cytosolic to total mRNA ratios in NSP9-expressing versus parental cells, whereas the cytosolic mRNA levels of non-targeted controls *MALAT1* and *UBC* were not significantly lowered (**Fig. 5g**). Even though nuclear RNA fractions were purified at high yields (>1 µg/µl), the RT-qPCR CT values of the target genes were too high (>25 cycles) for accurate quantification.

Interleukin 1α (IL-1α) and interleukin 1β (IL-1β) are important inflammatory cytokines constitutively produced in epithelial cells and plays a central role in regulating immune responses, including being a master cytokine in acute lung inflammation^37^. To determine if NSP9 inhibiting the nucleocytoplasmic export of the mRNA of IL-1α has any impact on the production of this cytokine, we performed an ELISA on the growth media of BEAS-2B wild type and NSP9 expressing cells 48 hours after induction by several common cytokines. Interferon α, β and γ resulted in lowered IL-1α levels in NSP9 cells compared to wild type, though tumor necrosis factor alpha (TNFα) resulted in the greatest reduction (∼ 30%) (**Fig. 5h**). We reproduced the observation of reduced IL-1α produced at different concentrations of TNFα (**Fig. 5i**). In addition, we observed reduced IL-1β produced in NSP9 expressing cells than in wildtype BEAS-2B cells (**Fig. 5j**). Thus, NSP9 association with the nuclear pore complex proteins aligns with the observation of decreased cytoplasmic abundance of NSP9 target mRNAs, suggesting that NSP9 interaction may directly inhibit nuclear export. Further, NSP9 reduced the production of its target gene IL-1α/β, which suggests that the export inhibition mechanism may be a strategy that SARS-CoV-2 employs to dampen inflammatory host response.

Taken together, we observed similarities in intronic protein-RNA interactions by non-structural proteins 2, 5, 7, and 9, which resembles the binding profile of splicing factor U2AF2. We further showed that NSP9 reduces cellular mRNA export, likely by interfering with U2AF/NXF1 substrate recognition. Our findings suggest NSP9 may contribute to the viral effort in suppressing host gene expression.

## Discussion

In this study, we performed a systematic and comprehensive survey of the SARS-CoV-2 protein-host RNA interactions using eCLIP. First, we performed eCLIP on NSP8, NSP12 and N proteins in SARS-CoV-2 infected Vero E6 cells and identified both host and viral RNAs bound by these proteins. We found that NSP12 and NSP8 bound specifically to the 5′ UTR and 3′ UTR of the virus genome, signaling their role in genome replication, whereas N bound nonspecifically to the entire region of the virus genome. We also found that NSP12 and NSP8 strongly enriched regions upstream of leader transcription regulatory site and the negative sense strand of the virus genome, providing further evidence of their involvement, but not N’s, in generating mRNAs from negative sense RNAs. Several major peaks are found across the genome that overlap highly structured regions. A distinctly strong peak near the 3′ end of NSP3 on the positive sense strand was identified from the NSP12 eCLIP. NSP12 may be involved in RNA polymerase transcriptional stalling and recombination with co-infected viruses in the evolutionary history of SARS-CoV-2. Of the host proteins recently identified to interact with NSP12^1^, SLU7 is the only splicing regulator^38^, which may be recruited by NSP12 for virus genome splicing. However, without further evidence, the functional significance and mechanism of NSP12 binding to region 7436-7526 are unknown and await future investigations. As some of the protein-RNA interaction peaks appear highly conserved and structured, these RNA sequences could serve as potential targets for broadly neutralizing antiviral drugs such as RNA-targeting small molecules^39^ to protect against future coronavirus outbreaks.

Recent RNA interactome capture studies suggest that SARS-CoV-2 proteins interact with polyA RNAs in virus infected cells, which include host mRNAs^40^. In our virus eCLIP, many host transcripts were bound by the viral replicase proteins in the context of virus infected cells. These host target genes are generally upregulated upon viral infection, including both antiviral and pro-viral genes. However, the functional impact of the protein-RNA interactions was difficult to isolate based on whole virus infection data and led us to study the protein-RNA interactions in isolation next. Our findings also prompted us to hypothesize that more viral proteins are likely involved in interacting with host RNAs. Due to a lack of antibodies specific to SARS-CoV-2 proteins and the limited infectivity of large numbers of human cells needed for sufficient transcriptome coverage in eCLIP libraries, we next investigated the viral protein-host RNA interactions in lung epithelial cell lines expressing epitope tagged SARS-CoV-2 proteins.

We found a total of 4773 coding genes, or a third of the transcriptome of the human lung cell line BEAS-2B, targeted by individually expressed SARS-CoV-2 proteins and verified some of the viral proteins using orthogonal assays RNA interactome capture (RIC) and crosslinking and solid-phase purification (CLASP). Not only do the proteins interact with distinct target mRNAs, but the sequence motifs and regional preferences are also varied. The rich eCLIP dataset has enabled us to derive binding principles, from which we found 8 of the proteins with a strong preference for binding to the 5′ UTR (NSP12, ORF3b, ORF7b, ORF9c) and CDS (NSP2, NSP3, NSP6, NSP14) regions. Using MS2-tethering dual luciferase assays, we then functionally characterized these proteins to show that they significantly upregulate target mRNA levels, with a combination of mRNA stabilization and translation activation activities. NSP1 in SARS-CoV is known to induce endonucleolytic cleavage of host translated mRNAs^17, 41, 42^, and similarly in SARS-CoV-2, it has been demonstrated to reduce cytosolic transcripts^43, 44^. In our reporter assay, NSP1 upregulates the expression of the target reporter, when recruited there by MS2-hairpins. This agrees with previous findings that its interaction with viral 5′ leader RNA protects its own RNA from the global depletion of cytosolic RNA^10, 43, 45^. Our eCLIP and RIC findings indicate no direct interaction between NSP1 and cellular mRNAs. This may imply that global mRNA degradation is not facilitated by NSP1 interaction with host mRNAs, though transient nucleolytic events may not be captured by UV crosslinking and subsequent RNA preparation for sequencing.

Finally, we demonstrated by overexpression studies that NSP12 has enhancing effects on the RNA levels of endogenous target genes, while ORF9c displayed translation enhancing effects. We then showed that siRNA knockdown of host genes targeted by SARS-CoV-2 proteins with gene expression enhancing effects restricted viral infection or proliferation. Thus, we presented a potential for viral protein-host RNA interactions in upregulating host genes that are required for viral propagation.

We also found that NSP9 significantly interacts with >900 transcripts and, together with its association with the nuclear pore complex, inhibits mRNA export of its target RNAs. We further demonstrated that NSP9 inhibits mRNA export of the IL-1α/β cytokine and reduces its production. Our findings shed light on a direct RNA targeting mechanism that viral proteins may employ to disrupt host mRNA nucleocytoplasmic transport. Recently, NSP1 was shown to interact with NXF1 to prevent its binding with mRNA export factors^46^. Since cytosolic mRNA levels are depleted, the increase in intronic reads was found to be more likely driven by mRNA degradation and/or mRNA export inhibition than alternative splicing as a result of viral infection^43^. Thus, our results contribute further evidence that SARS-CoV-2 proteins could leverage nuclear mRNA export inhibition as a strategy to dampen host antiviral response^46–48^.

A recent report^10^ used a crosslinking and Halotag/Halolink resin pulldown method to investigate viral protein-host RNA interactions. They screened 26 of the 29 SARS-CoV-2 proteins, each cloned with an N-terminal Halotag fusion and expressed in HEK293 cells (in contrast to more relevant lung epithelial cells) and found a total of only ∼148 host RNAs targeted by only 10 of the SARS-CoV-2 proteins. Consistent with the HaloTag-based results, NSP1 was observed to be enriched at the mRNA entry channel of the 18S ribosomal subunit (**Supplementary Fig. 5b**). However, it is unclear why we observe such a dramatic increase in peaks identified with eCLIP versus Halotag-CLIP. It is possible that their assay conditions were overly stringent, as GAPDH, which was used as a negative control, is extensively annotated as an RNA binding protein^49^. Our results in eCLIP, RIC, and CLASP (performed under denaturing conditions) consistently showed that NSP2 interacts with host RNAs, even though the HaloTag method did not pulldown any. Although antibody-based immunoprecipitation as performed in our study is less specific than the purification of HaloTag-coupled proteins due to the denaturing washes possible during HaloTag-pulldown, our use of the same FLAG and STREP tags and antibodies for all experiments provides a control for the possibility of antibody-based background interactions, and we have previously observed limited background in anti-FLAG eCLIP experiments in wild-type cells^50^. Nevertheless, we applied stringent filtering to further remove potential background peaks: viral protein eCLIP peaks were filtered by eCLIP peaks found in the wild-type BEAS-2B cells and cells overexpressing only the 3xFLAG and 2xStrep peptides, where anti-FLAG and anti-Strep antibodies were used in the immunoprecipitation step. It is also possible that the large (∼33 kDa) size of the N-terminal HaloTag inhibits some interactions or proper localization of viral proteins that are better captured with the smaller (∼2.7 kDa) FLAG and (∼2.9 kDa) Strep tags. Finally, without extensive engineering of the SARS-CoV-2 genome, antibody-based immunoprecipitation was the only viable way of studying viral protein-RNA in a whole virus infection context.

Like many viruses, the host-viral interactions underlying SARS-CoV-2 infection is broadly understood in terms of the virus hijacking the host cell by globally shutting down the expression of host genes that are irrelevant or hostile to its replication^51^, while the host attempts to fight off the virus by mounting apoptotic and inflammatory responses. To add to this understanding, we propose that viral proteins interact with host RNAs to activate a subset of host genes for its own survival through targeted translation activation or mRNA stabilization. We show that NSP12 and ORF9c specifically upregulate genes in the processes of protein N-linked glycosylation, mitochondrial ATP synthesis and transport, and COPII vesicle formation. We also propose that NSP9 contributes another layer to dampening host gene expression by inhibiting mRNA export. Understanding specifically upregulated processes and pro-viral genes will enable the development of new antiviral strategies. Our extensive and comprehensive dataset of SARS-CoV-2 protein-host RNA interactions provide a rich resource for understanding host-virus interactions and to enable development of new therapeutic strategies for acute COVID-19 and potential future coronavirus pandemics.

## Methods

### Cell culture and cell line generation

BEAS-2B, HEK293T and Vero E6 cells were purchased from the American Type Culture Collection and were not further authenticated. Cells were routinely tested for mycoplasma contamination with a MycoAlert mycoplasma test kit (Lonza) and were found negative for mycoplasma. The ACE2-overexpressing A549 cell line (A549-ACE2) was clonally generated and a gift from Benjamin tenOever^52^. BEAS-2B cells were cultured on Matrigel (Corning) coated plates and maintained in the PneumaCult-Ex Plus Medium (Stem Cell Technologies), supplemented with 33 µg/ml hydrocortisone (Stem Cell Technologies). Growth media was replaced every two days, and the cells were passaged every four days. HEK293T, Vero E6 and A549-ACE2 cells were cultured in DMEM (ThermoFisher) supplemented with 10% FBS (ThermoFisher) and passaged every three days. All cell cultures were incubated at 37°C and 5% CO_2_.

BEAS-2B cells expressing 2xStrep tagged SARS-CoV-2 proteins were generated using lentiviral transduction and purified using 1 µg/ml puromycin for two days. Lentiviral particles were packaged and harvested from HEK293T cells. To prepare cells for eCLIP using the BEAS-2B cell lines, 2 million cells were seeded in 15 cm dishes and 20 ml of growth media, and crosslinked four days after seeding (∼20 million cells per plate for each eCLIP replicate sample). Plasmids with 3xFLAG tagged SARS-CoV-2 proteins were transiently expressed in BEAS-2B cells using Lipofectamine 3000 (ThermoFisher) transfection according to manufacturer instructions. Cells were seeded 24 hours before transfection, growth media was replaced 24 hours after transfection, and cells were UV crosslinked 3 days after transfection.

For human lung organoid generation, we used a previously published protocol^53^. In short, human ESCs (H9, WiCell) were dissociated into single cells, and then seeded onto Matrigel-coated plates (BD Biosciences) at a density of 1.75 x 105 cells/cm^2^ in Definitive Endoderm (DE) induction medium (RPMI1640, B27 supplement, 1% HEPES, 1% glutamax, 50 U/mL penicillin/streptomycin), supplemented with 100 ng/mL human activin A (R&D), 1 µM CHIR99021 (Stemgent), and 10µM ROCK inhibitor, Y-27632 (R&D Systems) on day 1 then only activin A on days 2 and 3. Anterior Foregut Endoderm (AFE) was generated by supplementing serum free basal medium (3:1 IMDM:F12, B27+N2 supplements, 50 U/mL penicillin/streptomycin, 0.25% BSA, 0.05 mg/mL L-ascorbic acid, 0.4 mM monothioglycerol) with 10 µM SB431542 (R&D) and 2 µM Dorsomorphin (StemGent) on days 4-6. On day 7, AFE cells were dissociated and embedded in matrigel. Lung Progenitor Cell (LPC) induction medium, containing serum free basal medium supplemented with 10 ng/mL human recombinant BMP4 (R&D), 0.1 µM all-trans retinoic acid (Sigma-Aldrich) and 3 mM CHIR99021 was added for 9-11 days. To generate 3D human lung organoids, LPCs were dissociated in Dispase (StemCellTech) and resuspended in Matrigel in a 12-well Transwell 0.4µm pore size Transwell (Corning) culture insert. 3D lung organoid induction medium (FGF7 (10 ng/mL), FGF10 (10 ng/mL), CHIR (3 mM), EGF (10 ng/mL) in serum free basal medium) was added to the lower chamber and changed every 2 days for 6 days. On day 25, media was changed to 3D lung branching medium consisting of FGF7 (10 ng/mL), FGF10 (10 ng/mL), CHIR (3 μM), RA (0.1 μM), EGF (10 ng/mL) and VEGF/PIGF (10 ng/mL) in serum free basal medium. Media was changed every 2 days for 6 days. Finally, 3D lung maturation media was added consisting of the 3D lung branching medium supplemented with Dexamethasone (50 nM), cAMP (100 μM) and IBMX (100 μM). Media was changed every 2 days for 7 days. For infections, the 3D organoids were dissociated into single cells using Dispase and TrypLE (Gibco). This study protocol was approved by the Institutional Review Board of UCSD’s Human Research Protections Program (181180).

### SARS-CoV-2 virus infection

SARS-CoV-2 isolates USA-WA1/2020 (BEI Resources, #NR-52281), hCoV-19/USA/CA_UCSD_5574/2020 (lineage B.1.1.7) and hCoV-19/South Africa/KRISP-K005325/2020 (lineage B.1.351 BEI Resources NR-54009) were propagated and infectious units quantified by plaque assay using Vero E6 cells for the WA1 variant, and Vero-TMPRSS2 cells for variants B.1.1.7 and B.1.351. For eCLIP assays, Vero E6 cells were seeded at 5 million cells 24 hours before infection. About an hour before infection, the culture media was changed from 10% FBS to 2% FBS in DMEM. Cells were infected with the WA1 variant at a multiplicity of infection (MOI) of 0.01 and incubated for 48 hours. Infected cells were then rinsed with 1XPBS and in a biosafety hood in the BSL3 laboratory. While still in the BSL3 laboratory, cells were UV crosslinked at 400 mJ•cm^−2^, 254 nm on a prechilled metal block, pelleted by centrifugation at 300 g for 3 min, and snap frozen before transferring out of BSL 3 to the BSL2 laboratory. Frozen cell pellets were stored at −80°C until ready for eCLIP processing in the same way as the BEAS-2B cells.

For immunofluorescence staining assays, A549-ACE2 cells were seeded at 20,000 cells per well of an 8-well chamber slide (Millipore), which was pre-coated with Matrigel (Corning) in DMEM media supplemented with 10% FBS. 24 hours after seeding, the growth media was changed to DMEM supplemented with 2% FBS before infecting the cells at an MOI of 3 for 48 hours. Viruses were inactivated by soaking entire slides/plates in 4% paraformaldehyde (PFA) for 30 min at room temperature, before transferring from BSL3 to BSL2 laboratory.

### eCLIP library preparation and sequencing

The eCLIP experiment was performed as previously described^12^. Confluent cells were rinsed with 1XPBS and UV-cross-linked (400 mJ•cm^−2^, 254 nm) on ice, before cell lysis. Lysates were sonicated and treated with RNase I to fragment RNA. Two percent volume of each lysate sample was stored for preparation of a parallel SMInput library. The remaining lysates were immunoprecipitated using 15 µl anti-Strep or 10 µl anti-FLAG antibody (depending on the epitope tag of the construct; **Supplementary Table 4**) on Sheep Anti-Mouse IgG Dynabeads M-280 (ThermoFisher) overnight at 4°C. Negative control samples are wild type (WT) BEAS-2B cells, and performed using both anti-Strep and anti-FLAG antibodies (separately). Bound RNA fragments in the immunoprecipitates were dephosphorylated and 3ʹ-end ligated to an RNA adaptor. Protein–RNA complexes from SMInputs and immunoprecipitates were run on an SDS– polyacrylamide gel and transferred to nitrocellulose membrane. Membrane regions comprising the exact RBP sizes to 75 kDa above were excised, and RNA was released from the complexes with proteinase K. For negative control WT samples, two sizes are cut: 10 kDa – 85 kDa, and 85 kDa – 225 kDa. SMInput samples were dephosphorylated and 3ʹ-end ligated to an RNA adaptor. All RNA samples (immunoprecipitates and SMInputs) were reverse transcribed with AffinityScript (Agilent). cDNAs were 5ʹ-end ligated to a DNA adaptor. cDNA yields were quantified by qPCR, and 100–500 fmol of library was generated with Q5 PCR mix (NEB).

### Analysis of eCLIP sequencing data

For eCLIP performed on SARS-CoV-2 infected Vero E6 cells, reads were adapter trimmed and mapped to the African Green Monkey GCA_000409795.2/ChlSab2 and SARS-CoV-2 MN908947.3 genome assemblies. PCR duplicate reads were removed using the unique molecular identifier sequences in the 5ʹ adaptor, and remaining reads were retained as ‘usable reads’. For reads mapped to the SARS-CoV-2 genome, bedgraph densities were generated using SAM tools v1.9 to obtain read densities at each nucleotide position. The eCLIP enrichment for each position *x* is computed as a ratio of read densities *R* in the IP versus INPUT samples.

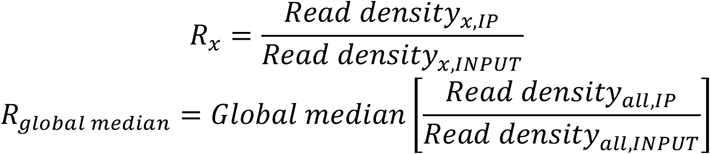

The relative positional enrichment was obtained by normalizing the positional enrichment by the global median ratio, i.e. *R_x_*/ *R_global median_*, and denoted ΔΔReadDensity. The mean of 2 biologically independent replicates was taken and referenced in this study.

For reads mapped to the African Green Monkey genome, peaks were called on the usable reads by CLIPper^54^ and assigned to gene annotations in Ensembl ChlSab1.1 release 102 and Sars_cov_2 ASM985889 v3.101 were used to annotate peaks mapped to the African Green Monkey and SARS-CoV-2 genome. Each peak was normalized to the size matched input (SMInput) by calculating the fraction of the number of usable reads from the IP sample relative to the usable reads from the SMInput sample. Reproducible peaks were defined as peaks that pass a cutoff of fold change of >8-fold and p-value of <0.001 using the irreproducible discovery rate (IDR) analysis, which is an analysis methodology^12, 55^ used to assess replicate agreement.

For eCLIP performed in BEAS-2B cells overexpressing epitope tagged proteins, Rreads were processed as described^11^. Briefly, reads were adapter trimmed and mapped to human-specific repetitive elements from RepBase (version 18.05) by STAR^56^, and ‘usable reads’ were obtained exactly as above. For identifying eCLIP peaks, reads mapped to repeat elements were removed, and remaining reads were mapped to human genome assembly hg19 with STAR. PCR duplicate reads were removed using the unique molecular identifier sequences in the 5ʹ adaptor, and remaining reads were retained as ‘usable reads’. Peaks were called on the usable reads by CLIPper^54^ and assigned to gene regions annotated in GENCODE v19 with the following order of descending priority: CDS, 5ʹ UTR, 3ʹ UTR, proximal intron and distal intron. Proximal intron regions are defined as extending up to 500 bp from an exon–intron junction. Peaks were normalized to size-matched input and IDR analysis was performed to identify reproducible peaks exactly as above. Peaks are also filtered for ≥20 bases in length, and not overlapping with WT negative control samples or samples expressing 3xFLAG and 2xStrep peptides. Target transcripts were defined as transcripts that contained at least one significant reproducible peak. Code is available on GitHub (https://github.com/YeoLab/eclip, and https://github.com/yeolab/merge_peaks). Gene Ontology analysis of eCLIP target genes was performed using ENRICHR (https://maayanlab.cloud/Enrichr/https://maayanlab.cloud/Enrichr/). Jaccard Index for comparing eCLIP target transcripts for SARS-CoV-2 proteins and ENCODE RBPs are computed as 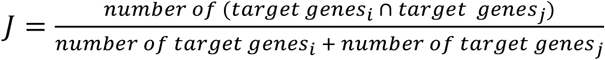 for viral protein *i* and ENCODE protein *j*. Cluster maps were visualized using Cytoscape version 3.8.1.

### RNA sequencing and analysis

A549-ACE2 cells infected with SARS-CoV-2 WA1 at MOI of 3 for 48 hours and Vero E6 cells infected at MOI of 0.1 for 48 hours were treated with TRIzol (Thermo Fisher), which inactivated the virus, and purified with Direct-zol RNA kits (Zymo). Uninfected cells were seeded and treated for RNA purification in parallel to serve as controls. Confluent 10 cm dishes of BEAS-2B cells were processed similarly. 500 ng of purified RNA in each replicate and condition was used for strand-specific RNA-seq library preparation using the Illumina Stranded mRNA Prep kit (Illumina, Cat. 20040534) and IDT Illumina RNA UD Indexes Set B (Illumina, Cat. 20040554). Libraries were sequenced on the NovasSeq at a depth of at least 50 million reads per sample in Paired End 100 mode. RNA-seq reads were trimmed of adaptor sequences using cutadapt (v1.144.0) and for A549-ACE2 and BEAS-2B cells, mapped to repetitive elements (RepBase v18.04) using STAR (v 2.5.2bv2.4.0i). Reads that did not map to repetitive elements were then mapped to the human genome (hg19) for the human cell lines. For SARS-CoV-2 infected Vero E6 cells, repeat element mapping was not performed, and reads were directly mapped to the African Green Monkey GCA_000409795.2/ChlSab2 and SARS-CoV-2 MN908947.3 genome assemblies. GENCODE v19 gene annotations and featureCounts (v.1.5.30) were used to create read count matrices. Code is available on GitHub (https://github.com/YeoLab/eclip).

RNA-seq read density was visualized using bed graph density values generated using SAM tools version 1.9. Splice junction arches were generated and visualized in the Integrated Genome Viewer version 2.8.13. using .bam files that were downsampled to 1 percent of the original .bam files using SAM tools.

### Polysome fractionation

Polysome fractionation was performed as described previously^57^. Briefly, to obtain crude lysates, cell cultures were washed once with PBS containing Cycloheximide (CHX; 100µg/ml), harvested by cell scraping and then lysed on ice using 20 mM Tris HCl pH 7.4, 150 mM NaCl, 5 mM MgCl2, 1 mM DTT with 1% Triton-X + Protease Inhibitors + RNase inhibitors + CHX (100 µg/ml). Nuclei and debris were separated from crude lysate by centrifugation at 15,000 g at 4°C for 5 min. Sucrose gradients (10%–50%) were prepared in 20 mM Tris HCl pH 7.4, 150 mM NaCl, 5 mM MgCl2, 1 mM DTT + RNase inhibitors + CHX (100ug/ml) using a Biocomp Model 108 gradient master. Crude cellular lysates were then loaded onto gradients and separated by centrifugation at 110,000 g, 3 hours at 4°C and fractionated into 0.5mL aliquots using a Biocomp Model 152 Piston Fractionator. Polysome fractions (typically fractions #13 through #24) were pooled and RNA extraction/purification was performed for the preparation of sequencing libraries using the Illumina Stranded mRNA Prep kit (Illumina, Cat. 20040534) and IDT Illumina RNA UD Indexes Set B (Illumina, Cat. 20040554). Libraries were sequenced on the NovaSeq at a depth of at least 25 million reads per sample in Paired End 100 mode. Sequencing reads are first processed as RNA-seq libraries, where RNA-seq reads were trimmed of adaptor sequences using cutadapt (v1.4.0) and mapped to repetitive elements (RepBase v18.04) using STAR (v2.4.0i). Reads that did not map to repetitive elements were then mapped to the human genome (hg19) for the human cell lines.

Only transcripts with read count >50 were considered. The change in polysome enrichment of any sample condition *i* relative to any control condition *0* can be represented by a ratio of ratios. More specifically, we have ratios representing the polysome enrichment in condition *P_i_* is normalized by polysome enrichment in control condition *P_0_*.

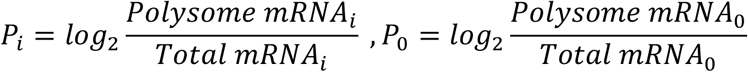

The change in polysome enrichment is the ratio *P*_i_ − *P*_0_ and denoted ΔLog_2_FoldChange.

### Filter binding assay

Filter binding assay was performed as described previously^57^. Double stranded templates were made from first performing PCR (Roche, KAPA HiFi HotStart ReadyMix) using primers Scov2_7431_7555_left + Scov2_7431_7555_right_revcomp for the sequence from region 7431_7555, and Scov2_7431_7555_Scrambled_left + Scov2_7431_7555_Scrambled_right_revcomp for the scrambled control. This was followed by another PCR using primers T7_fwd + Scov2_7431_7555_rev and T7_fwd + Scov2_7431_7555_Scrambled_rev, respectively, to prepend the T7 promoter. *In vitro* transcription was performed on the purified templates using the MegaShortScript kit (ThermoFisher), and column purified RNA was biotinylated using Pierce RNA 3’ End Biotinylation Kit (ThermoFisher). Recombinant His-tagged NSP12 (R&D Systems, catalog # 10686-CV) was incubated with *in vitro* transcribed and biotinylated RNA in 20 mM Tris, 200 mM KCl at room temperature for 1 hr. A sandwich of three membranes was assembled in a dot blot apparatus (Biorad) consisting of a top layer of Polyethersulfone (Millipore PES, 0.45 um pore size), middle layer 100% Nitrocellulose (GE, Hybond ECL Nitrocellulose) and bottom layer Nylon (GE, Hybond Nylon-N+). Membranes were washed twice with 20mM Tris, 200mM KCl before and after the application of samples. Membranes were crosslinked using a Stratalinker (4000 J). Blots were visualized with Streptavidin HRP (Chemiluminescent Nucleic Acid Detection Module, Thermo-Pierce).

### Multiple sequence alignment and phylogenetic analysis

Complete genomes of betacoronavirus reference sequences from NCBI were downloaded on April 5^th^, 2021, and bat and pangolin coronavirus complete genome sequences were downloaded from GISAID on April 6^th^, 2021. Sequence accession codes are displayed in Fig. 1h. Multiple sequence alignment was performed using MAFFT v7.453 and default parameters, and sequence alignment was visualized using Jalview (version 1.0). Consensus sequence score was generated in Jalview, and the consensus sequence and consensus RNA structure of region in the alignment corresponding to position 7470-7510 in SARS-CoV-2 was generated using the RNAAliFold prediction. The phylogenetic tree was constructed using the average distance algorithm from the multiple sequence alignment and visualized within Jalview.

### Plasmid construction

2xStrep-tagged plasmids in a pLVX vector expressing SARS-CoV-2 proteins were a gift from Nevan Krogan^1^. Plasmids containing NSP3, NSP4, NSP13, NSP14 and NSP16 fused with a 3XFLAG tag and cloned into a pcDNA3.4 vector (**Supplementary Table 5**) were codon optimized using the same protein sequence based off the reference sequence (NC_045512.2) and were synthesized by GeneArt (ThermoFisher). Plasmids containing NSP1, NSP5, NSP7, NSP8, NSP11, NSP12, Spike, ORF3b, ORF7b, ORF8, ORF9b, ORF9c, ORF10 fused with a 3XFLAG tag are cloned into a pcDNA3.4 vector using flanking primers (**Supplementary Table 6**) ordered from Integrated DNA Technologies by PCR amplifying from the pLVX plasmids (**Supplementary Table 5**). Cloning into the pcDNA3.4 was performed using FastDigest restriction enzymes EcoRI and BshT1 (Invitrogen) and Gibson assembly (NEB).

MCP-tagged SARS-CoV-2 expression plasmids for the MS2 tethering assay were generated by Gateway Assembly (ThermoFisher). SARS-CoV-2 ORFs were amplified by PCR (KAPA HiFi HotStart ReadyMix, Roche) from the 2xStrep-tagged or 3xFLAG-tagged plasmids with oligonucleotide primers containing attB recombination sites and recombined into pDONR221 using BP clonase II (ThermoFisher) (**Supplementary Table 6**). ORFs were then recombined into a custom pEF DEST51 destination vector^32^ (ThermoFisher) engineered to direct expression of the ORFs as fusion proteins with a V5 epitope tag and MCP appended C terminally and under the control of the EF1-alpha promoter to create ORF–V5–MS2BP constructs. The MCP-tagged BOLL and CNOT7 expressing plasmids were similarly taken from the previously reported large scale MS2-tethering screening assay^32^.

### Repeat-family-centric mapping

Binding to rRNA was analyzed using a family-aware repeat element mapping pipeline^12^. In the pipeline, reads were mapped to a database of 7,419 multicopy element transcripts, including the 5S, 5.8S, 18S and 28S rRNAs as well as tRNAs, retrotransposable elements and numerous other RNAs. Reads mapping to multiple element families were not considered for further analysis. Fold enrichment of reads mapped to IP samples are normalized by INPUT samples for individual replicates. Code is available on GitHub (https://github.com/YeoLab/repetitive-element-mapping).

### De novo motif analysis

HOMER was used to identify *de novo* motifs using reads from IDR peaks. The foreground was a bed file of significant IDR peaks; the background was randomly defined peaks within the same annotated region as the foreground peaks. Code is available on GitHub (https://github.com/YeoLab/clip_analysis_legacy).

### Crosslinking and solid-phase purification (CLASP)

CLASP was performed as previously described in Kim, Arcos et al., 2021 with slight variation described in brief below. For each experiment, one 10cm plate of HEK293T cells was used. To stabilize protein-RNA interactions, the growth media was removed and cells were washed twice in 1 x PBS and irradiated on a cold block with UV_254nm_ (400 mJ/cm^2^). Cells were then scrapped and collected and pelleted in cold PBS. Cells were then lysed and denatured in denaturation buffer (50 mM Tris-HC, pH 6.8, 10% glycerol, 2.5% SDS, 0.66% NP-40) and sonicated with Bioruptor Pico (Diagenode) for 30 seconds on and 30 seconds off for a total of 5 minutes. Lysate was then incubated for 10 mins at 95°C and moved to RT for additional 10 mins. To capture crosslinked protein-RNA complexes, 0.66x of SPRI beads (Hawkins et al., 1994) (1 mg/ml SPRI beads in 10 mM Tris–HCL, pH= 8.0, 1 M NaCl, 18% PEG-8000, 1 mM EDTA and 0.055% Tween-20) were added to lysate and incubated at RT for 10 minutes. SPRI beads were then washed 5 times in denaturing buffer (30 mM Tris–HCl, pH 6.8, 6% glycerol, 1.5% SDS, 0.4% NP-40, 1 M NaCl, 8% PEG-8000) to remove all non-specific interactions. The crosslinked RNA-protein complexes were then eluted from SPRI beads using denaturation buffer (50 mM Tris-HC, pH 6.8, 10% glycerol, 2.5% SDS, 0.66% NP-40) and underwent benzonase treatment to degrade all nucleic acid. Protein was then precipitated using methanol/chloroform extraction protocol. Extracted proteins was then resuspended in 2x NuPage LDS running buffer + DTT and run on SDS-page and transferred to nitrocellulose for immunoblotting.

### RNA Interactome Capture (RIC)

RIC was performed as previously described in Perez-Perri et al., 2013. In brief, crosslinked cell pellets were resuspended in ice-cold lysis buffer (20 mM Tris–HCl, pH 7.5, 500 mM LiCl, 1 mM EDTA, 5 mM DTT, and 0.5% (wt/vol) LiDS, 5 mM DTT and complete protease inhibitor cocktail) and incubated on ice for 5 minutes. Cells were then sonicated with Bioruptor Pico (Diagenode) for 30 seconds on and 30 seconds off for a total of 5 minutes. Insolubles were then removed by spinning lysate at 15,000 g for 5 minutes and supernatant moved to a new tube. Oligo dt beads (NEB) were added and incubated in lysate for 1 hour at 37°C with gentle rotation. Beads were collected with magnet, and supernatant was transferred to a new tube for a second round of capture. Beads were then subject to successive rounds of washes using wash buffers 1-3 (buffer 1: 20 mM Tris–HCl, 500 mM LiCl, 1 mM EDTA, 5 mM DTT, and 0.5% LiDs; buffer 2: 20 mM Tris–HCl, 500 mM LiCl, 1 mM EDTA, 5 mM DTT, and 0.1% LiDs; buffer 3: 20 mM Tris–HCl, 200 mM LiCl, 1 mM EDTA, 5 mM DTT, and 0.02% LiDs) with 5 minutes of gentle rotation. RNA-protein interactions were eluted off the beads using RNAse free water and combined with 10× RNase buffer, 1 M DTT, and 1% NP40 (final concentrations: 1× RNase buffer, 5 mM DTT, 0.01% NP40) and ∼200 U RNase T1 and RNase A (Sigma-Aldrich). RNA was digested for 60 min at 37 °C. Eluted proteins were resuspended in 2x NuPage LDS running buffer + DTT and run on SDS-page and transferred to nitrocellulose for immunoblotting.

### Metagene mapping analyses

Metagene plots were created using the intersection of eCLIP peaks and a set of mRNA regions. To generate the list of each CDS, 5ʹ UTR and 3ʹ UTR, non-overlapping gene annotations from GENCODE v19 were used. First, low-expression transcripts (TPM < 1) from BEAS-2B cells were removed. Then, transcripts with the highest TPM were selected, resulting in a single transcript per gene in the CDS. For each 5ʹ UTR, CDS and 3ʹ UTR in a gene, the entire set of exons making up the region was concatenated and overlapped with eCLIP peaks, resulting in a vector of positions across the transcript containing values of 1 if a peak was found at a given position or 0 otherwise. Plotted lines represent the number of total peaks found at each position divided by the total number of unique transcripts. The length of each region within the metagene was then scaled to 8%, 62% and 30%, corresponding to the average length of regions from the most highly expressed transcripts in ENCODE HepG2 RNA-seq control datasets. The peak density was calculated as the percentage of peaks at a given position (https://github.com/YeoLab/rbp-maps).

To visualize eCLIP read truncation densities on exon-intron-exon regions, we fetched reads for each pre-mRNA transcript. Truncation sites were defined as the 5ʹ end of read2 for pair-end eCLIP, and 5ʹ end of every read for single-end eCLIP. The number of truncation sites from the INPUT library were subtracted from the IP library. Then, the total subtracted signal was normalized with the total subtracted signal on the pre-mRNA transcript. Interval features such as introns and exons were extracted from the Gencode v19 coordinates. Since each region interval can have different lengths for different transcripts, two windows slicing from the 5ʹ end or the 3ʹ end were created, up to a length of 150 bases. No scaling was applied for this metagene visualization. Finally, the density in each feature was normalized across all IDR peak-containing transcripts, creating the average metagene density map. The density signal was smoothed using gaussian kernel density with sigma = 5 bases. To call alternative splicing (AS) events from deeply sequenced RNA-seq data, we used rMATs 3.2.5 with gencode v19 annotations. Significant AS events are defined with the threshold FDR < 0.1, inclusion level difference > 0.05.

### MS2-tethering dual luciferase assay

The Renilla–MS2 and Firefly reporter constructs were taken from a previously reported MS2-tethering dual luciferase assay^32^. A 3:1:1 mix of MCP-tagged SARS-CoV-2 protein expression plasmid, Renilla–MS2 and Firefly reporter constructs was mixed with Lipofectamine 3000 reagents following the manufacturer’s directions (ThermoFisher). The transfection mixture was added to PDL-coated 96-well plates, with a total of 100 ng DNA per well of a 96-well plate. HEK293T cells were added to each well at a count of 20,000 cells/well for a reverse transfection. Cells were lysed 48 hours post transfection, and luciferase activity was measured with the Dual-Luciferase Reporter Assay System (Promega), in a microplate reader (Spark, Tecan). Luciferase substrate was added to all wells, then reads with 10 second integration times were performed. Values were expressed as the ratio of the mean luciferase activity of MS2-tagged renilla luciferase over MS2-untagged firefly luciferase from three replicates and normalized to this ratio from the negative control – an MCP-tagged FLAG epitope plasmid.

### MS2-tethering dual reporter RT-qPCR

RT-qPCR validation was performed on cells transfected under the same conditions as the dual luciferase assay. Total RNA was isolated by lysing cells in TRIzol (Thermo Fisher) and purified with Direct-zol RNA kits (Zymo), following the manufacturers’ protocols. Reverse transcription of 50 ng total RNA was performed using Protoscript II First Strand cDNA Synthesis Kit with oligo(dT)23 primers (NEB). cDNA was undiluted, and target transcripts were quantified with Power SYBR Green Master Mix (Thermo Fisher) using gene specific primers (**Supplementary Table 6**). Three biological replicate samples from independently transfected cells were assayed, and RT–qPCR was carried out in three technical replicates. Mean Ct values were calculated from each triplicate set for each biological replicate. Biological replicates were averaged to generate mean fold changes, and values expressed as fold differences to control samples were calculated using the ΔΔCt method.

### Co-Immunoprecipitation

For each co-IP sample, one confluent 10 cm dish of cells was used. Cells were washed with PBS, scraped, and centrifuged at 200 g for 5 min to pellet. Cell pellets were snap frozen for storage at − 80°C until use. Dynabead M-280 Sheep Anti-Mouse IgG (Invitrogen) magnetic beads were washed three times using TBS+0.05% Tween-20 (TBST) before incubating with 5 µg anti-Strep antibody (ref antibody list) for 45 min with rotation at room temperature. Cell pellets were resuspended and lysed in 500 µl of gentle, non-denaturing lysis buffer (20 mM Tris-HCl pH 8.0, 137 mM NaCl, 1% NP-40 (Igepal), 2 mM EDTA, Protease Inhibitor Cocktail Set III (EMD Millipore)) on ice for 30 min. After cell lysis, lysates were centrifuged at 20,000 g at 4°C for 10 min. Antibody bound beads were washed three times with TBST before resuspending in 100 µl of the gentle lysis buffer. The remaining of the cleared lysate was added to the resuspended beads and incubated overnight at 4°C with rotation. After overnight incubation, About 20 µl or 4% of the cleared lysate was set aside as the INPUT sample to check for antibody integrity and protein expression. The remaining IP samples were washed in chilled lysis buffer three times, before resuspending in 60 µl lysis buffer. INPUT and IP samples were carried forward to Western blotting.

### Western blot

Cells were washed with PBS and lysed in lysis buffer (50 mM Tris-HCl, 100 mM NaCl, 1% NP- 40, 0.1% SDS, 0.5% sodium deoxycholate; pH 7.4) with Protease Inhibitor Cocktail Set III (EMD Millipore). Lysates were sonicated in a water bath sonicator (Diagenode) at 4 °C for 5 min with 30-s on/off pulses at the low setting. Protein extracts were denatured at 75 °C for 20 min and run at 150 V for 1.5 h on 4-12% NuPAGE Bis-Tris gels in NuPAGE MOPS running buffer (Thermo Fisher). Proteins were transferred to polyvinylidene difluoride membrane using NuPAGE transfer buffer (Thermo Fisher) with 10% methanol. Membranes were blocked in blocking buffer (TBS containing 5% (wt/vol) dry milk powder) for 30 min and probed with primary antibodies in blocking buffer for 16 h at 4 °C. Membranes were washed three times with TBST and probed with secondary HRP-conjugated antibodies in blocking buffer for 1 h at room temperature. Signal was detected by Pierce ECL substrate (Thermo Fisher) and imaged on an Azure Biosystems C600 imager.

### siRNA knockdown assay

Human PSC derived lung organoids were dissociated into single cells and seeded at 15,000 cells per well of a 96-well plate one day before transfection. siRNAs were ordered from Integrated DNA Technologies (**Supplementary Table 7-8**). 25 nM of siRNAs were transfected using the Lipofectamine RNAiMAX reagent (ThermoFisher). Growth media was replaced one day after transfection. Two days after siRNA transfection, the growth media was replaced with the base media for the lung organoid cells. Lung organoid cells were infected by SARS-CoV-2 at an MOI of 1 for 24 hours, respectively. Cells were fixed with 4% paraformaldehyde for 30 min, which inactivates the virus, before transferring from BSL3 to BSL2, and proceeding with immunofluorescence staining using anti-Nucleocapsid antibody (40143-R019, Sino Biological).

### Immunofluorescence

Fixed cells were permeabilized with PBS with 0.25% Triton X-100 (PBST) and blocked with blocking buffer (5% goat serum in PBST) for 1 h at room temperature. Next, cells were incubated with primary antibodies (**Supplementary Table 4**) at 1:250-2000 dilutions in blocking buffer for 16 h at 4 °C, washed with PBS+0.01% Triton X-100 three times for 5 min each at room temperature, and then incubated with secondary antibody (goat anti-rabbit secondary IgG (H+L) Superclonal Recombinant Secondary Antibody, Alexa Fluor 488 or Alexa Fluor 555 (Invitrogen)) in blocking buffer for 1 h. After staining, cells were washed again in PBST three times for 5 min each at room temperature. Staining of nuclei with 4ʹ,6-diamidino-2-phenylindole (DAPI) was performed with mounting solution (ProLong Diamond Antifade Mountant with DAPI (ThermoFisher)) or 50% glycerol in 1×PBS.

Chamber slide images were captured on a ZEISS Axiocam 503 epifluorescence microscope camera with a 40X objective. Images were collected via Zeiss ZEN software and converted to tiff for downstream analysis. Images were analyzed using a custom-developed pipeline in CellProfiler (v.3.1.09). First, cell nuclei were segmented using the Dapi channel. Cell boundaries were then identified using the watershed algorithm with identified nuclei as seed. Virus fluorescent signal (Alexa Fluor 555 staining for SARS-CoV-2 NSP8) was thresholded to identify virus infected cells. Finally, the relative fluorescent intensity of protein of interest of virus-infected and -noninfected cells were calculated. For imaging 96-well plates, the IncuCyte S3 was used to measure GFP fluorescence and its software was used to determine total integrated intensity. A Keyence BZ-X800 microscope was used to count the number of cells using the DAPI channel. The total integrated intensity of GFP fluorescence was divided by the cell count and normalized to the scrambled siRNA sequence control to determine infection rate.

### ELISA

Wildtype BEAS-2B and NSP9 expressing BEAS-2B cells were seeded at 100,000 cells per well of a 24 well plate (pre-coated with Matrigel). One day after seeding, cytokines (IL-6, IFNα, β and γ and TNFα) were added to a final concentration of 100 pg/µl, unless otherwise specified. 48 hours after stimulation, growth media was collected and stored at −80°C until use. The LEGEND MAX Human IL-1α ELISA Kit (Biolegend) was used to assay for IL-1α concentration, and The LEGEND MAX Human IL-1β ELISA Kit (Biolegend) was used to assay for IL-1β concentration. The sample absorbance was measured on a Tecan Infinite M200 Pro plate reader.

### Subcellular fractionation

Subcellular fractionation was performed as described previously^58^ with minor modifications. Briefly, one confluent 10 cm tissue culture plate (corresponding to ∼8 million cells) was used for each fractionation sample, and two independent replicates were performed. BEAS-2B wild type and NSP9 expressing cells were rinsed once with ice-cold PBS and then harvested by scraping and resuspension in 1ml of ice-cold PBS. Cells were centrifuged at 200g for 3min at 4°C, the supernatant removed and the pellets either processed directly or snap-frozen and stored at −80°C until use.

For fractionation, cell pellets were thawed on ice and resuspended in 1 ml of hypotonic lysis buffer (20 mM Tris HCl pH 7.5, 10 mM KCl, 1.5 mM MgCl, 5 mM EGTA, 1 mM EDTA, 1 mM DTT) supplemented with protease inhibitor and 20ul RNAse inhibitor (RNAseOUT). Cells were incubated on ice for 15min, transferred into a 2ml dounce homogenizer with a tight-fitting (type B) pestle and gently homogenized using 8 strokes to lyse the cells while keeping nuclei intact. This and all subsequent homogenization steps were performed on ice at all times. After homogenization, 1/10th volume (100-150 ul) was removed as the total input fraction and mixed with 3 volumes of Trizol LS (300-450 ul). The remaining lysate was transferred into a 1.5 ml tube and centrifuged at 1200g for 10min at 4°C to pellet cell nuclei. After the first spin, the supernatant was transferred into a fresh 1.5ml tube for two additional repeats of the 1200g spin. The nuclei pellets from the first 1200g spin were gently rinsed with 250 µl of hypotonic lysis buffer and resuspended in 1 ml 0.32 M sucrose buffer (0.32 M sucrose, 3 mM CaCl2, 2 mM MgOAc, 0.1 mM EDTA, 10mM Tris Cl pH8.0, 1mM DTT, 0.5% v/v NP-40) supplemented with protease and RNAse inhibitors. The nuclei pellets in 0.32M sucrose buffer were transferred into a clean 2ml dounce homogenizer and resuspended using 3 strokes of a tight-fitting pestle. After addition of 1 ml of 2 M sucrose buffer (2 M Sucrose, 5 mM MgOAc, 0.1 mM EDTA, 10 mM Tris pH8.0, 1 mM DTT) supplemented with protease and RNAse inhibitors, the nuclei suspension was mixed and gently transferred to create a layer on top of a 1ml cushion of 2M sucrose buffer in a 3ml ultracentrifuge tube. The tubes were transferred into a SW50.1 swinging bucket rotor and centrifuged at 30,000g for 30min at 4°C. After the spin, the supernatant was removed, and the pellet was rinsed twice with 500 µl of 0.32M sucrose buffer. The rinsed nuclear pellet was then resuspended by trituration in 250 µl of hypotonic lysis buffer and 750 µl of Trizol LS were added. This is the nuclear fraction.

To obtain the cytoplasmic fraction, 10 ul of TurboDNAse was added and mixed into the supernatant from the third 1200 g spin in 1.5 ml ultracentrifuge tubes. The samples were then centrifuged at 100,000 g for 1 h at 4°C in a tabletop ultracentrifuge using a TLA110 fixed-angle rotor. After the spin, the supernatant was transferred into a fresh 5ml tube and 3 volumes of Trizol LS were added. This is the cytoplasmic fraction. All fractions are stored at −80°C until use.

RNA was purified from the samples using the Direct-zol kit (Zymo Research). Reverse transcription was performed according to manufacturer instructions using the Superscript IV kit (ThermoFisher) using an oligo(dT) primer. Gene specific primers (**Supplementary Table 6**) were used in the qPCR, performed with the Power SYBR Green Master Mix (Thermo Fisher) on a BIO-RAD CFX 384-well qPCR thermocycler to quantify transcript levels in each fraction.

## Data availability

Plasmids and cell lines generated in this work are available upon request. All sequencing data are deposited in GEO with accession GSE173508.

## Acknowledgements

We would like to appreciate members of the Yeo lab for providing helpful discussions. The work has been supported by Emergency COVID-19 Research Seed Funding (#R00RG2636) from the University of California Office of the President. This publication includes data generated at the UC San Diego IGM Genomics Center utilizing an Illumina NovaSeq 6000 that was purchased with funding from a National Institutes of Health SIG grant (#S10 OD026929). J.S.X as a visiting fellow is partially supported by Agency for Science, Technology and Research (A*STAR) and Industrial Alignment Fund Pre-Positioning (IAF-PP) grant H17/01/a0/012. ELVN is supported by the NHGRI (R00HG009530). AFC is supported by an NIH grant K08 AI130381 and a Burroughs Wellcome Fund Career Award for Medical Scientists. The following reagents was deposited by the Centers for Disease Control and Prevention and obtained through BEI Resources, NIAID, NIH: SARS-Related Coronavirus 2, Isolate USA-WA1/2020, NR-52281. The following reagent was obtained through BEI Resources, NIAID, NIH: SARS-Related Coronavirus 2, Isolate hCoV-19/South Africa/KRISP-K005325/2020, NR-54009, contributed by Alex Sigal and Tulio de Oliveira.

## Author contributions

J.S.X. and G.W.Y. conceived of the project. J.S.X, J.R.M, E-C.L, D.S, J.C.S, F.E.T, K.R., K.W.B, R.N.M, A.T., A.F.C. and S.L.L designed and performed experiments. K.L.J, S.S.P, E.M.K, Y-H.L., K.D.D performed experiments. J.S.X, J.R.M, E-C.L, D.S, J.C.S, F.E.T, K.R., K.W.B, P.L., A.Q.V, Y.S, and S.L.L analyzed experimental results. J.S.X, E-C.L, B.A.Y, H-L.H, C-Y.C, W.J, E.K. and E.L.V.N performed bioinformatics and structural analysis. J.S.X, C-Y.C., S.L.L and G.W.Y wrote the manuscript with help from all authors. G.W.Y supervised the project.

