## Supplementary information for "Discovery and functional interrogation of SARS-CoV-2 protein-RNA interactions"

**Supplementary Figure Legends**

**Supplementary Figure 1.** **eCLIP results from SARS-CoV-2 infected cells mapped to the positive sense genome. a)** Tracks showing coverage in each sample across the genome in the positive strand. Gray lines indicate positions with no coverage. Values in parenthesis indicate fraction of coverage. **b)** Matrix showing correlation of read densities between every sample as indicated on the horizontal and vertical axes. Dot size and color scale with the Pearson’s coefficient of each comparison, which is indicated in each dot. **c)** 2D density plots comparing replicate correlation of fold change in read density, log_2_(IP/IN) for each of NSP12, NSP8 and N.

Line is a linear regression model, and R indicates Pearson’s coefficient. **d)** Read densities mapped across the SARS-CoV-2 genome in the positive sense for each of NSP12, NSP8 and N proteins in both immunoprecipitated (IP) and input samples. **e)** Relative positional enrichment at the 5′ end. SL1, Stem Loop 1; TRS, transcriptional regulatory site. f**)** Relative positional enrichment at the 3′ end. S2M, Stem loop 2 motif.

**Supplementary Figure 2.** **eCLIP results from SARS-CoV-2 infected cells mapped to the positive sense genome. a)** Stacked bar plots showing total read densities mapped to position and negative strands of the SARS-CoV-2 genome in each eCLIP sample. **b)** Tracks showing coverage in each sample across the genome in the negative strand. Gray lines indicate positions with no coverage. Values in parenthesis indicate fraction of coverage. **c)** Read densities mapped across the SARS-CoV-2 genome in the negative sense for each of NSP12, NSP8 and N proteins in both immunoprecipitated (IP) and input samples.

**Supplementary Figure 3.** **Structured regions enriched by NSP12 eCLIP of SARS-CoV-2 infected cells. a)** Secondary structures that overlap with the major peaks enriched in NSP12 eCLIP at positions 3533-3635,17202-17222, 21177-21206, and 24018-24079. **b)** Filter-binding assay showing RNA crosslinked to NSP12 on the nitrocellulose layer, and the free RNA on the nylon layer. **c)** Multiple sequence alignment of the region corresponding among the SARS-CoV-2 reference sequence, and homologous bat and pangolin sequences. Consensus sequence and structure are indicated underneath the aligned sequences. **d)** Phylogenetic tree constructed from c). **e)** RNA-seq read density plot from SARS-CoV-2 infected A549-ACE2 cells mapping sequenced reads to the sense strand of the SARS-CoV-2 genome.

**Supplementary Figure 4.** **Correlations of eCLIP results of SARS-CoV-2 infected cells** Matrix showing correlation of mean log_2_(fold change) of eCLIP read density between samples indicated on the horizontal and vertical axes (bottom left, 2D density plots with linear regression line; top right, dots showing Pearson’s coefficient); the diagonal panels show the histogram distribution of mean log_2_(fold change) values for samples indicated in the horizontal axis. Mean is taken for n = 2 biologically independent samples.

**Supplementary Figure 5.** **SARS-CoV-2 protein-host RNA interaction characteristics.**
**a)** Stacked bar plot showing number of reproducible eCLIP peaks (IDR) that target coding and non-coding RNAs for each SARS-CoV-2 protein individually expressed in BEAS-2B. **b)** Locations of NSP1 (blue) binding sites on rRNAs. Line plots show the fold enrichment of read coverage at each position (blue) and the mean of 446 other RBPs analyzed by the ENCODE consortium (grey; https://www.encodeproject.org/, accession code ENCSR456FVU) on 18S and 28S rRNAs (lightly shaded areas indicate 10–90% confidence intervals).

**Supplementary Figure 6. eCLIP target cluster maps.** Cluster maps of eCLIP target genes (blue boxes connected by red edges to yellow box at center), clustered by top GO terms, for SARS-CoV-2 proteins (yellow box at center). Grey edges are human protein-protein interaction data from Mentha downloaded on 2/17/2020.

**Supplementary Figure 7. Sequence motifs from IDR peaks from eCLIP performed in BEAS-2B cells.**

**Supplementary Figure 8.** **Immunofluorescence of SARS-CoV-2 infected cells.**
**a)** Segmentation performed in CellProfiler on nucleus and cytoplasm of A549-ACE2 cells infected by SARS-CoV-2. Virus marker is NSP8, and nuclei are stained by DAPI. **b)** An example image of lung organoid cells infected by SARS-CoV-2. Segmentation and quantification are performed in the Keyence software, using nucleocapsid as virus marker and DAPI to stain for nuclei. **c)** Bar plot showing mean relative fluorescence intensities of cells from h, dots represent segmented individual cells (mean ± s.d. * p<0.05, ** p<0.005, **** p<0.0001, two-tailed Welch’s t-test).

**Supplementary Figure 9.** **Characteristics of intron associated SARS-CoV-2 proteins**
**a)** Violin plot showing distribution of Include Level Difference of significantly, alternatively spliced genes (FDR < 0.1, |Include Level Difference| > 0.05) in A549-ACE2 cells infected with SARS-CoV-2. KS test p values indicate significance of difference in Include Level Difference of SARS-CoV-2 protein target genes versus all genes. **b)** Jaccard index similarity of NSP9 target genes as compared with all 223 ENCODE RBP datasets. **c)** Normalized read density of NSP9 enriched RNA mapped to averaged exonic and intronic regions for every transcript (rows). **d-e)** Uncropped Western blot images of NSP9 co-IP showing anti-Strep immunoprecipitation of 2xStrep tagged NSP9, and blotting for d) NUP62 and e) 2xstrep tagged NSP9.

**Supplementary Tables**

**Supplementary Table 1.** SARS-CoV-2 genome regions with >5-fold ∆∆ReadDensity

|  | **Chromosome** | **Start** | **End** | **MaxRel.Enrichment** | **PeakMaxPosition** | **PeakLength** | **Strand** | **Sample** |
| --- | --- | --- | --- | --- | --- | --- | --- | --- |
| **1** | MN908947.3 | 559 | 579 | 5.4 | 567 | 20 | + | N |
| **2** | MN908947.3 | 939 | 961 | 5.6 | 945 | 22 | + | N |
| **3** | MN908947.3 | 3688 | 3713 | 5.9 | 3701 | 25 | + | N |
| **4** | MN908947.3 | 6635 | 6650 | 5.6 | 6640 | 15 | + | N |
| **5** | MN908947.3 | 9602 | 9620 | 9.5 | 9605 | 18 | + | N |
| **6** | MN908947.3 | 14138 | 14158 | 6.4 | 14149 | 20 | + | N |
| **7** | MN908947.3 | 26896 | 26917 | 5.5 | 26902 | 21 | + | N |
| **1** | MN908947.3 | 1 | 133 | 573.2 | 1 | 132 | + | NSP12 |
| **2** | MN908947.3 | 211 | 867 | 27.0 | 391 | 656 | + | NSP12 |
| **3** | MN908947.3 | 932 | 1028 | 11.3 | 983 | 96 | + | NSP12 |
| **4** | MN908947.3 | 1237 | 1369 | 8.8 | 1270 | 132 | + | NSP12 |
| **5** | MN908947.3 | 3533 | 3635 | 18.6 | 3547 | 102 | + | NSP12 |
| **6** | MN908947.3 | 5906 | 5924 | 8.4 | 5908 | 18 | + | NSP12 |
| **7** | MN908947.3 | 6042 | 6067 | 9.7 | 6046 | 25 | + | NSP12 |
| **8** | MN908947.3 | 6427 | 6508 | 8.8 | 6435 | 81 | + | NSP12 |
| **9** | MN908947.3 | 6564 | 6602 | 6.8 | 6570 | 38 | + | NSP12 |
| **10** | MN908947.3 | 7436 | 7526 | 13.7 | 7489 | 90 | + | NSP12 |
| **11** | MN908947.3 | 17202 | 17222 | 9.2 | 17206 | 20 | + | NSP12 |
| **12** | MN908947.3 | 21177 | 21206 | 11.2 | 21181 | 29 | + | NSP12 |
| **13** | MN908947.3 | 24018 | 24079 | 12.5 | 24022 | 61 | + | NSP12 |
| **14** | MN908947.3 | 25541 | 25558 | 6.8 | 25547 | 17 | + | NSP12 |
| **15** | MN908947.3 | 27188 | 27280 | 9.0 | 27202 | 92 | + | NSP12 |
| **16** | MN908947.3 | 27576 | 27629 | 7.5 | 27581 | 53 | + | NSP12 |
| **17** | MN908947.3 | 27645 | 27733 | 6.9 | 27689 | 88 | + | NSP12 |
| **18** | MN908947.3 | 28391 | 28404 | 5.4 | 28398 | 13 | + | NSP12 |
| **19** | MN908947.3 | 28912 | 28933 | 5.8 | 28920 | 21 | + | NSP12 |
| **20** | MN908947.3 | 29035 | 29052 | 5.4 | 29041 | 17 | + | NSP12 |
| **21** | MN908947.3 | 29555 | 29693 | 23.8 | 29579 | 138 | + | NSP12 |
| **22** | MN908947.3 | 29794 | 29880 | 12.4 | 29819 | 86 | + | NSP12 |
| **1** | MN908947.3 | 1 | 68 | 106.8 | 1 | 67 | + | NSP8 |
| **2** | MN908947.3 | 226 | 323 | 12.6 | 273 | 97 | + | NSP8 |
| **3** | MN908947.3 | 332 | 380 | 6.8 | 348 | 48 | + | NSP8 |
| **4** | MN908947.3 | 382 | 513 | 21.3 | 435 | 131 | + | NSP8 |
| **5** | MN908947.3 | 520 | 592 | 22.5 | 565 | 72 | + | NSP8 |
| **6** | MN908947.3 | 603 | 892 | 15.1 | 642 | 289 | + | NSP8 |
| **7** | MN908947.3 | 922 | 1010 | 10.7 | 950 | 88 | + | NSP8 |
| **8** | MN908947.3 | 1215 | 1369 | 8.8 | 1242 | 154 | + | NSP8 |
| **9** | MN908947.3 | 2463 | 2473 | 5.5 | 2470 | 10 | + | NSP8 |
| **10** | MN908947.3 | 2568 | 2628 | 7.4 | 2599 | 60 | + | NSP8 |
| **11** | MN908947.3 | 3452 | 3489 | 7.1 | 3465 | 37 | + | NSP8 |
| **12** | MN908947.3 | 3534 | 3726 | 10.1 | 3548 | 192 | + | NSP8 |
| **13** | MN908947.3 | 4868 | 4886 | 6.2 | 4874 | 18 | + | NSP8 |
| **14** | MN908947.3 | 4932 | 4947 | 5.7 | 4932 | 15 | + | NSP8 |
| **15** | MN908947.3 | 5389 | 5419 | 9.0 | 5398 | 30 | + | NSP8 |
| **16** | MN908947.3 | 6016 | 6095 | 7.8 | 6041 | 79 | + | NSP8 |
| **17** | MN908947.3 | 6425 | 6473 | 8.9 | 6438 | 48 | + | NSP8 |
| **18** | MN908947.3 | 6478 | 6495 | 5.2 | 6482 | 17 | + | NSP8 |
| **19** | MN908947.3 | 6561 | 6602 | 7.4 | 6574 | 41 | + | NSP8 |
| **20** | MN908947.3 | 6626 | 6677 | 7.8 | 6647 | 51 | + | NSP8 |
| **21** | MN908947.3 | 9602 | 9625 | 7.2 | 9605 | 23 | + | NSP8 |
| **22** | MN908947.3 | 9669 | 9683 | 5.6 | 9675 | 14 | + | NSP8 |
| **23** | MN908947.3 | 23574 | 23590 | 5.6 | 23579 | 16 | + | NSP8 |
| **24** | MN908947.3 | 24020 | 24048 | 7.1 | 24026 | 28 | + | NSP8 |
| **25** | MN908947.3 | 24069 | 24079 | 6.0 | 24071 | 10 | + | NSP8 |
| **26** | MN908947.3 | 25551 | 25579 | 5.7 | 25557 | 28 | + | NSP8 |
| **27** | MN908947.3 | 26896 | 26927 | 6.4 | 26902 | 31 | + | NSP8 |
| **28** | MN908947.3 | 27152 | 27162 | 5.4 | 27156 | 10 | + | NSP8 |
| **29** | MN908947.3 | 27170 | 27218 | 6.8 | 27190 | 48 | + | NSP8 |
| **30** | MN908947.3 | 27221 | 27282 | 8.3 | 27248 | 61 | + | NSP8 |
| **31** | MN908947.3 | 27400 | 27430 | 11.3 | 27406 | 30 | + | NSP8 |
| **32** | MN908947.3 | 27674 | 27719 | 8.1 | 27691 | 45 | + | NSP8 |
| **33** | MN908947.3 | 28922 | 28932 | 5.3 | 28929 | 10 | + | NSP8 |
| **34** | MN908947.3 | 29025 | 29052 | 7.6 | 29035 | 27 | + | NSP8 |
| **35** | MN908947.3 | 29562 | 29614 | 7.6 | 29582 | 52 | + | NSP8 |
| **36** | MN908947.3 | 29626 | 29693 | 8.5 | 29656 | 67 | + | NSP8 |
| **37** | MN908947.3 | 29818 | 29868 | 6.9 | 29831 | 50 | + | NSP8 |

**Supplementary Table 2.** Nucleic acid sequences used in filter binding assay

| **Name** | **Type** | **Sequence 5ʹ —> 3ʹ** |
| --- | --- | --- |
| **T7_fwd** | Single stranded DNA oligonucleotide | TAATACGACTCACTATAGGGCGAAAACGCCC |
| **Scov2_7431_7555_left** | Single stranded DNA oligonucleotide | ACTATAGGGCGAAAACGCCC CATTTTATTATGTATGGAAAAGTTATGTGCATGTTGTAGACGGTTGTAATTCATCAACTTGTATGATG |
| **Scov2_7431_7555_right_revcomp** | Single stranded DNA oligonucleotide | AACACCATTAACAATAGTTGTACATTCGACTCTTGTTGCTCTATTACGTTTGTAACACATCATACAAGTTGATGAATTACAACCG |
| **Scov2_7431_7555_rev** | Single stranded DNA oligonucleotide | AACACCATTAACAATAGTTGTACATTCG |
| **Scov2_7431_7555_Scrambled_left** | Single stranded DNA oligonucleotide | ACTATAGGGCGAAAACGCCC GAATTTTATCATGACTGAACTGTTTTTGTGTCCCAAAAATAGTGCTACATTAGGAAAATAATTAAGATTA |
| **Scov2_7431_7555_Scrambled_right_revcomp** | Single stranded DNA oligonucleotide | CCCTCCTGATAAGTAAACCCTAGTACTAAATAACATATTAACTCCAATTCAGCCGTAATCTTAATTATTTTCCTAATGTAGCACTATTTT |
| **Scov2_7431_7555_Scrambled_rev** | Single stranded DNA oligonucleotide | CCCTCCTGATAAGTAAACCCTAG |
| **Scov2_7431_7555_PCR** | Double stranded DNA, PCR product | TAATACGACTCACTATAGGGCGAAAACGCCCCATTTTATTATGTATGGAAAAGTTATGTGCATGTTGTAGACGGTTGTAATTCATCAACTTGTATGATGTGTTACAAACGTAATAGAGCAACAAGAGTCGAATGTACAACTATTGTTAATGGTGTT |
| **Scov2_7431_7555_Scrambled_PCR** | Double stranded DNA, PCR product | TAATACGACTCACTATAGGGCGAAAACGCCCACTATAGGGCGAAAACGCCC GAATTTTATCATGACTGAACTGTTTTTGTGTCCCAAAAATAGTGCTACATTAGGAAAATAATTAAGATTACGGCTGAATTGGAGTTAATATGTTATTTAGTACTAGGGTTTACTTATCAGGAGGG |
| **Scov2_7431_7555_RNA** | In vitro transcribed RNA | GGGCGAAAACGCCCCAUUUUAUUAUGUAUGGAAAAGUUAUGUGCAUGUUGUAGACGGUUGUAAUUCAUCAACUUGUAUGAUGUGUUACAAACGUAAUAGAGCAACAAGAGUCGAAUGUACAACUAUUGUUAAUGGUGUU |
| **Scov2_7431_7555_Scrambled_RNA** | In vitro transcribed RNA | GGGCGAAAACGCCCGAAUUUUAUCAUGACUGAACUGUUUUUGUGUCCCAAAAAUAGUGCUACAUUAGGAAAAUAAUUAAGAUUACGGCUGAAUUGGAGUUAAUAUGUUAUUUAGUACUAGGGUUUACUUAUCAGGAGGG |

**Supplementary Table 3.** Enriched Gene Ontology processes in NSP12 eCLIP in SARS-CoV-2 infected cells

|  | **Gene Ontology Term** | **Adjusted P-value** |
| --- | --- | --- |
| **1** | regulation of transcription from RNA polymerase II promoter (GO:0006357) | 1.3E-08 |
| **2** | positive regulation of transcription, DNA-templated (GO:0045893) | 6.9E-08 |
| **3** | positive regulation of transcription from RNA polymerase II promoter (GO:0045944) | 2.8E-06 |
| **4** | regulation of apoptotic process (GO:0042981) | 2.8E-06 |
| **5** | positive regulation of gene expression (GO:0010628) | 5.7E-05 |
| **6** | regulation of peptidyl-threonine phosphorylation (GO:0010799) | 2.6E-04 |
| **7** | positive regulation of nucleic acid-templated transcription (GO:1903508) | 4.2E-04 |
| **8** | regulation of cell migration (GO:0030334) | 5.4E-04 |
| **9** | regulated exocytosis (GO:0045055) | 5.4E-04 |
| **10** | positive regulation of cell motility (GO:2000147) | 6.1E-04 |
| **11** | platelet degranulation (GO:0002576) | 6.2E-04 |
| **12** | MAPK cascade (GO:0000165) | 7.8E-04 |
| **13** | regulation of cell proliferation (GO:0042127) | 7.8E-04 |
| **14** | positive regulation of gene silencing by miRNA (GO:2000637) | 8.7E-04 |
| **15** | positive regulation of binding (GO:0051099) | 1.2E-03 |
| **16** | negative regulation of gene expression (GO:0010629) | 1.2E-03 |
| **17** | regulation of protein autophosphorylation (GO:0031952) | 1.9E-03 |
| **18** | regulation of cell cycle (GO:0051726) | 1.9E-03 |
| **19** | positive regulation of cell migration (GO:0030335) | 2.7E-03 |
| **20** | transmembrane receptor protein tyrosine kinase signaling pathway (GO:0007169) | 2.8E-03 |
| **21** | positive regulation of peptidyl-threonine phosphorylation (GO:0010800) | 3.0E-03 |
| **22** | negative regulation of transcription, DNA-templated (GO:0045892) | 3.0E-03 |
| **23** | positive regulation of cellular protein metabolic process (GO:0032270) | 3.0E-03 |
| **24** | protein kinase B signaling (GO:0043491) | 3.5E-03 |
| **25** | positive regulation of cellular process (GO:0048522) | 3.5E-03 |
| **26** | positive regulation of cell proliferation (GO:0008284) | 3.5E-03 |
| **27** | regulation of fibroblast proliferation (GO:0048145) | 3.7E-03 |
| **28** | negative regulation of programmed cell death (GO:0043069) | 3.7E-03 |
| **29** | negative regulation of apoptotic process (GO:0043066) | 3.7E-03 |
| **30** | regulation of transcription, DNA-templated (GO:0006355) | 3.7E-03 |
| **31** | stress-activated protein kinase signaling cascade (GO:0031098) | 3.7E-03 |
| **32** | negative regulation of transcription from RNA polymerase II promoter (GO:0000122) | 3.7E-03 |
| **33** | positive regulation of protein phosphorylation (GO:0001934) | 3.8E-03 |
| **34** | cell-cell junction organization (GO:0045216) | 4.4E-03 |
| **35** | positive regulation of protein kinase activity (GO:0045860) | 4.4E-03 |
| **36** | cellular response to transforming growth factor beta stimulus (GO:0071560) | 5.2E-03 |
| **37** | transmembrane receptor protein serine/threonine kinase signaling pathway (GO:0007178) | 5.3E-03 |
| **38** | mRNA 3'-end processing (GO:0031124) | 5.3E-03 |
| **39** | ERK1 and ERK2 cascade (GO:0070371) | 5.8E-03 |
| **40** | regulation of cell-matrix adhesion (GO:0001952) | 6.9E-03 |
| **41** | regulation of stress-activated MAPK cascade (GO:0032872) | 6.9E-03 |
| **42** | transcription, DNA-templated (GO:0006351) | 6.9E-03 |
| **43** | regulation of mitotic cell cycle (GO:0007346) | 6.9E-03 |
| **44** | positive regulation of protein autophosphorylation (GO:0031954) | 6.9E-03 |
| **45** | regulation of programmed cell death (GO:0043067) | 7.6E-03 |
| **46** | branching morphogenesis of an epithelial tube (GO:0048754) | 8.3E-03 |
| **47** | regulation of cell-substrate junction assembly (GO:0090109) | 8.3E-03 |
| **48** | negative regulation of RNA metabolic process (GO:0051253) | 8.3E-03 |
| **49** | cellular response to corticosteroid stimulus (GO:0071384) | 8.3E-03 |
| **50** | negative regulation of peptidyl-threonine phosphorylation (GO:0010801) | 8.3E-03 |
| **51** | positive regulation of intracellular signal transduction (GO:1902533) | 8.3E-03 |
| **52** | regulation of cell motility (GO:2000145) | 8.8E-03 |
| **53** | regulation of focal adhesion assembly (GO:0051893) | 8.9E-03 |
| **54** | transcription from RNA polymerase II promoter (GO:0006366) | 9.9E-03 |

**Supplementary Table 4.** Antibodies used in this work

| **Antibody** | **Cat No.** | **Species** | **Company** | **Purpose** |
| --- | --- | --- | --- | --- |
| **StrepMAB-Immo** | 2-1517-001 | Mouse | Iba life sciences | eCLIP, co-IP |
| **Monoclonal ANTI-FLAG® M2 antibody** | F3165 | Mouse | Sigma Aldrich | eCLIP |
| **SARS-CoV / SARS-CoV-2 (COVID-19) NSP8 antibody [5A10]** | GTX632696 | Mouse | GeneTex | virus eCLIP, immunofluorescence |
| **SARS-CoV-2 (COVID-19) RdRp (nsp12) antibody** | GTX135467 | Rabbit | GeneTex | virus eCLIP |
| **SARS-CoV-2 (2019-nCoV) Nucleocapsid Antibody** | 40143-R019 | Rabbit | Sino Biological | virus eCLIP, immunofluorescence |
| **NUP62 antibody** | A304-941A-M | Rabbit | Bethyl | Western blot |
| **NDUFA4 antibody** | GTX34093 | Rabbit | GeneTex | Western blot, immunofluorescence |
| **UGGT1 antibody** | A305-530A-M | Rabbit | Bethyl | Western blot, immunofluorescence |
| **RPN1/Ribophorin I Antibody** | A305-027A | Rabbit | Bethyl Laboratories | Western blot, immunofluorescence |
| **β-Actin (8H10D10) mAb** | # 3700 | Mouse | Cell Signaling Technology | Western blot |
| **GAPDH antibody** | MAB374 | Mouse | Millipore Sigma | CLASP |
| **Alpha-tubulin** | ab7291 | Mouse | abcam | CLASP, RIC |
| **YTHDC1** | A305-096A | Rabbit | Bethyl | CLASP, RIC |
| **ELAVL1** | 12582S | Rabbit | Cell Signaling Technology | CLASP |
| **TrueBlot Anti-Rabbit IgG HRP secondary antibody** | 18-8816-43 | Goat | Rockland Immunochemicals | Western blot |
| **TrueBlot Anti-Mouse IgG HRP secondary antibody** | 18-8817-33 | Goat | Rockland Immunochemicals | Western blot |
| **anti-Rabbit IgG (H+L), Superclonal Recombinant Secondary Antibody, Alexa Fluor 488** | A27034 | Goat | Invitrogen | Immunofluorescence |
| **anti-Mouse IgG (H+L), Superclonal Recombinant Secondary Antibody, Alexa Fluor 555** | A28180 | Goat | Invitrogen | Immunofluorescence |
| **IRDye 800CW Goat anti-Mouse IgG Secondary Antibody** | 926-32210 | Goat | Licor | Western blot |

**Supplementary Table 5.** Plasmids used in this work

| **Protein name** | **Construct name** | **Parent vector** | **Source** | **Experiment** |
| --- | --- | --- | --- | --- |
| **NSP1** | NSP1_3XFLAG | pcDNA3.4 | This work | eCLIP |
| **NSP2** | NSP2_2xStrep | pLVX | Gordon et al | eCLIP |
| **NSP3** | NSP3_3xFLAG | pcDNA3.4 | This work | eCLIP |
| **NSP4** | NSP4_3xFLAG | pcDNA3.4 | This work | eCLIP |
| **NSP5** | NSP5_3xFLAG | pcDNA3.4 | This work | eCLIP |
| **NSP5_C145A** | NSP5_C145A_2xStrep | pLVX | Gordon et al | eCLIP |
| **NSP6** | NSP6_3xFLAG | pcDNA3.4 | This work | eCLIP |
| **NSP7** | NSP7_3xFLAG | pcDNA3.4 | This work | eCLIP |
| **NSP8** | Nsp8_3XFLAG | pcDNA3.4 | This work | eCLIP |
| **NSP9** | NSP9_2xStrep | pLVX | Gordon et al | eCLIP |
| **NSP10** | NSP10_2xStrep | pLVX | Gordon et al | eCLIP |
| **NSP11** | NSP11_3xFLAG | pcDNA3.4 | This work | eCLIP |
| **NSP12** | NSP12_3xFLAG | pcDNA3.4 | This work | eCLIP |
| **NSP13** | NSP13_3xFLAG | pcDNA3.4 | This work | eCLIP |
| **NSP14** | N-3xFLAG_NSP14 | pcDNA3.4 | This work | eCLIP |
| **NSP15** | NSP15_2xStrep | pLVX | Gordon et al | eCLIP |
| **NSP16** | NSP16_3xFLAG | pcDNA3.4 | This work | eCLIP |
| **Spike** | Spike_3xFLAG | pcDNA3.4 | This work | eCLIP |
| **ORF3a** | ORF3a_2xStrep | pLVX | Gordon et al | eCLIP |
| **ORF3b** | ORF3b_3xFLAG | pcDNA3.4 | This work | eCLIP |
| **E** | E_2xStrep | pLVX | Gordon et al | eCLIP |
| **M** | M_2xStrep | pLVX | Gordon et al | eCLIP |
| **ORF6** | ORF6_2xStrep | pLVX | Gordon et al | eCLIP |
| **ORF7a** | ORF7a_2xStrep | pLVX | Gordon et al | eCLIP |
| **ORF7b** | N-3xFLAG_ORF7b | pcDNA3.4 | This work | eCLIP |
| **ORF8** | ORF8_3xFLAG | pcDNA3.4 | This work | eCLIP |
| **N** | N_2xStrep | pLVX | Gordon et al | eCLIP |
| **ORF9b** | ORF9b_3xFLAG | pcDNA3.4 | This work | eCLIP |
| **ORF9c** | N-3xFLAG ORF9c | pcDNA3.4 | This work | eCLIP |
| **ORF10** | ORF10_3xFLAG | pcDNA3.4 | This work | eCLIP |
| **3xFLAG** | 3xFLAG | pcDNA3.4 | This work | eCLIP |
| **2xStrep** | 2xStrep | pcDNA3.4 | This work | eCLIP |
| **NSP1** | NSP1–V5–MS2BP | pDONR221 | This work | MS2 tethering dual luciferase |
| **NSP2** | NSP2–V5–MS2BP | pDONR221 | This work | MS2 tethering dual luciferase |
| **NSP3** | NSP3–V5–MS2BP | pDONR221 | This work | MS2 tethering dual luciferase |
| **NSP5** | NSP5–V5–MS2BP | pDONR221 | This work | MS2 tethering dual luciferase |
| **NSP6** | NSP6–V5–MS2BP | pDONR221 | This work | MS2 tethering dual luciferase |
| **NSP9** | NSP9–V5–MS2BP | pDONR221 | This work | MS2 tethering dual luciferase |
| **NSP12** | NSP12–V5–MS2BP | pDONR221 | This work | MS2 tethering dual luciferase |
| **NSP14** | NSP14–V5–MS2BP | pDONR221 | This work | MS2 tethering dual luciferase |
| **NSP16** | NSP16–V5–MS2BP | pDONR221 | This work | MS2 tethering dual luciferase |
| **Spike** | Spike–V5–MS2BP | pDONR221 | This work | MS2 tethering dual luciferase |
| **ORF3b** | ORF3b–V5–MS2BP | pDONR221 | This work | MS2 tethering dual luciferase |
| **ORF7b** | ORF7b–V5–MS2BP | pDONR221 | This work | MS2 tethering dual luciferase |
| **N** | N–V5–MS2BP | pDONR221 | This work | MS2 tethering dual luciferase |
| **ORF9c** | ORF9c–V5–MS2BP | pDONR221 | This work | MS2 tethering dual luciferase |
| **FLAG** | FLAG–V5–MS2BP | pDONR221 | Luo et al | MS2 tethering dual luciferase |
| **BOLL** | BOLL–V5–MS2BP | pDONR221 | Luo et al | MS2 tethering dual luciferase |
| **CNOT7** | CNOT7–V5–MS2BP | pDONR221 | Luo et al | MS2 tethering dual luciferase |
|  | pEF DEST51 V5-MS2BP destination vector | pEF DEST51 | Luo et al | MS2 tethering dual luciferase |

**Supplementary Table 6.** Primers used in this work

| **Target** | **Primer name** | **Sequence 5ʹ —> 3ʹ** | **Experiment** | **Source** |
| --- | --- | --- | --- | --- |
| **Firefly luciferase** | Firefly_fwd | GCTAAGAAAGGCGGCAAGAT | RT-qPCR | Luo et al |
| **Firefly luciferase** | Firefly_rev | GGTGGATCACAAAGTGGCTTA | RT-qPCR | Luo et al |
| **Renilla luciferase** | Renilla_fwd | CGCTCCAGATGAAATGGGTA | RT-qPCR | Luo et al |
| **Renilla luciferase** | Renilla_rev | GGTGGATCACAAAGTGGCTTA | RT-qPCR | Luo et al |
| **IL1a** | IL1a_Keita_fwd | GAATGACGCCCTCAATCAAAGT | RT-qPCR | Keita et al |
| **IL1a** | IL1a_Keita_rev | TCATCTTGGGCAGTCACATACA | RT-qPCR | Keita et al |
| **UPP1** | UPP1_sahin_fwd | ACTGCCCAGGTAGAGACTATC | RT-qPCR | Sahin et al |
| **UPP1** | UPP1_sahin_rev | CTGCACCAGCTTCTTGTTAAG | RT-qPCR | Sahin et al |
| **ANXA2** | ANXA2_lu_fwd | TCTACTGTTCACGAAATCCTGTG | RT-qPCR | Lu et al |
| **ANXA2** | ANXA2_lu_rev | AGTATAGGCTTTGACAGACCCAT | RT-qPCR | Lu et al |
| **MALAT1** | MALAT1_yang_fwd | GAATTGCGTCATTTAAAGCCTAGTT | RT-qPCR | Yang et al |
| **MALAT1** | MALAT1_yang_rev | GTTTCATCCTACCACTCCCAATTAAT | RT-qPCR | Yang et al |
| **UBC** | NM_001111112(2) | GATTTGGGTCGCAGTTCTTG | RT-qPCR | IDT PrimeTime qPCR primers |
| **UBC** | NM_001111112(2) | CCTTATCTTGGATCTTTGCCTTG | RT-qPCR | IDT PrimeTime qPCR primers |
| **pLVX constructs** | pcDNA3.4_EcoRI_fwd | TCCGGACTCTAGAGGATCGAACCCTTGAATTCGCCGCCACCATG | Cloning 3XFLAG tagged constructs | This work |
| **pLVX constructs** | pLVX_1xstrep_rev | GTCTTTGTAATCTGAACCACCTCCCTCGAGGCCTTTCTCGAATTGTGGAT | Cloning 3XFLAG tagged constructs | This work |
| **pLVX constructs** | pcDNA3.4_XhoI_3XFLAG_BshTI_rev | GATTGTCGAGATATCAAACTCATTACTAACCGGTTTATTTATCGTCATCATCCTTGTAGTCGATGTCGTGATCCTTATAGTCGCCGTCATGGTCTTTGTAATCTGAACCACCTCCC | Cloning 3XFLAG tagged constructs | This work |
| **pLVX constructs** | pcDNA3.4_N-3XFLAG_fwd | CGGACTCTAGAGGATCGAACCCTTGAATTCGCCGCCACCATGGATTACAAAGACCATGACGGCGACTATAAGGATCACGACATCGACTACAAGGATGATGACG | Cloning 3XFLAG tagged constructs | This work |
| **pLVX constructs** | pcDNA3.4_N-3XFLAG_rev | TGATTGTCGAGATATCAAACTCATTACTAACCGGTGGCTTCGGCCAGTAACGTTA | Cloning 3XFLAG tagged constructs | This work |
| **NSP1** | NSP1-V5-MS2BP_fwd | GGGGACAAGTTTGTACAAAAAAGCAGGCTTAGCCGCCACCATGGAG | Cloning ORF-V5-MS2BP constructs | This work |
| **NSP1** | NSP1-V5-MS2BP_rev | GGGGACCACTTTGTACAAGAAAGCTGGGTTGCCGCCGTTCAGTTCG | Cloning ORF-V5-MS2BP constructs | This work |
| **NSP2** | NSP2-V5-MS2BP_fwd | GGGGACAAGTTTGTACAAAAAAGCAGGCTTAGCCGCCACCATGGCC | Cloning ORF-V5-MS2BP constructs | This work |
| **NSP2** | NSP2-V5-MS2BP_rev | GGGGACCACTTTGTACAAGAAAGCTGGGTTGCCGCCCTTAAGGGTAAAGG | Cloning ORF-V5-MS2BP constructs | This work |
| **NSP3** | NSP3-V5-MS2BP_fwd | GGGGACAAGTTTGTACAAAAAAGCAGGCTTAGCCGCCACCATGgc | Cloning ORF-V5-MS2BP constructs | This work |
| **NSP3** | NSP3-V5-MS2BP_rev | GGGGACCACTTTGTACAAGAAAGCTGGGTTTCCGCCTTTGAGGGCAA | Cloning ORF-V5-MS2BP constructs | This work |
| **NSP5** | NSP5-V5-MS2BP_fwd | GGGGACAAGTTTGTACAAAAAAGCAGGCTTAGCCGCCACCATGTCTG | Cloning ORF-V5-MS2BP constructs | This work |
| **NSP5** | NSP5-V5-MS2BP_rev | GGGGACCACTTTGTACAAGAAAGCTGGGTTCTGGAAAGTGACCCCACTGC | Cloning ORF-V5-MS2BP constructs | This work |
| **NSP6** | NSP6-V5-MS2BP_fwd | GGGGACAAGTTTGTACAAAAAAGCAGGCTTAGCCGCCACCATGTCTGC | Cloning ORF-V5-MS2BP constructs | This work |
| **NSP6** | NSP6-V5-MS2BP_rev | GGGGACCACTTTGTACAAGAAAGCTGGGTTCTGCACTGTAGCCACTTTGATACA | Cloning ORF-V5-MS2BP constructs | This work |
| **NSP7** | NSP7-V5-MS2BP_fwd | GGGGACAAGTTTGTACAAAAAAGCAGGCTTAGCCGCCACCATGAGC | Cloning ORF-V5-MS2BP constructs | This work |
| **NSP7** | NSP7-V5-MS2BP_rev | GGGGACCACTTTGTACAAGAAAGCTGGGTTCTGCAATGTCGCCCGG | Cloning ORF-V5-MS2BP constructs | This work |
| **NSP9** | NSP9-V5-MS2BP_fwd | GGGGACAAGTTTGTACAAAAAAGCAGGCTTAGCCGCCACCATGAATAATGAG | Cloning ORF-V5-MS2BP constructs | This work |
| **NSP9** | NSP9-V5-MS2BP_rev | GGGGACCACTTTGTACAAGAAAGCTGGGTTTTGGAGTCGCACGGTCG | Cloning ORF-V5-MS2BP constructs | This work |
| **NSP12** | NSP12-V5-MS2BP_fwd | GGGGACAAGTTTGTACAAAAAAGCAGGCTTAGCCGCCACCATGTCAG | Cloning ORF-V5-MS2BP constructs | This work |
| **NSP12** | NSP12-V5-MS2BP_rev | GGGGACCACTTTGTACAAGAAAGCTGGGTTCTGCAGGACGGTGTGAGG | Cloning ORF-V5-MS2BP constructs | This work |
| **NSP14** | NSP14-V5-MS2BP_fwd | GGGGACAAGTTTGTACAAAAAAGCAGGCTTAGCCGCCACCATGGCCGAAAATGTGACCGG | Cloning ORF-V5-MS2BP constructs | This work |
| **NSP14** | NSP14-V5-MS2BP_rev | GGGGACCACTTTGTACAAGAAAGCTGGGTTCTGCAGTCTGGTGAAGGTGT | Cloning ORF-V5-MS2BP constructs | This work |
| **NSP16** | NSP16-V5-MS2BP_fwd | GGGGACAAGTTTGTACAAAAAAGCAGGCTTAGCCGCCACCATGAGTTCT | Cloning ORF-V5-MS2BP constructs | This work |
| **NSP16** | NSP16-V5-MS2BP_rev | GGGGACCACTTTGTACAAGAAAGCTGGGTTATTGTTGACCAGCACGTCG | Cloning ORF-V5-MS2BP constructs | This work |
| **ORF3b** | ORF3b-V5-MS2BP_fwd | GGGGACAAGTTTGTACAAAAAAGCAGGCTTAGCCGCCACCATGGCATACTGCTGGAGATG | Cloning ORF-V5-MS2BP constructs | This work |
| **ORF3b** | ORF3b-V5-MS2BP_rev | GGGGACCACTTTGTACAAGAAAGCTGGGTTCGGCCAGCAGCATCG | Cloning ORF-V5-MS2BP constructs | This work |
| **ORF7b** | ORF7b-V5-MS2BP_fwd | GGGGACAAGTTTGTACAAAAAAGCAGGCTTAGCCGCCACCATGATTGAGCTGTCTCTCAT | Cloning ORF-V5-MS2BP constructs | This work |
| **ORF7b** | ORF7b-V5-MS2BP_rev | GGGGACCACTTTGTACAAGAAAGCTGGGTTGGCGTGGCATGTCTCG | Cloning ORF-V5-MS2BP constructs | This work |
| **N** | N-V5-MS2BP_fwd | GGGGACAAGTTTGTACAAAAAAGCAGGCTTAGCCGCCACCATGAGC | Cloning ORF-V5-MS2BP constructs | This work |
| **N** | N-V5-MS2BP_rev | GGGGACCACTTTGTACAAGAAAGCTGGGTTCGCCTGAGTAGAATCGGCT | Cloning ORF-V5-MS2BP constructs | This work |
| **ORF9c** | ORF9c-V5-MS2BP_fwd | GGGGACAAGTTTGTACAAAAAAGCAGGCTTAGCCGCCACCATGCTTCAATCCTGCTATAACTTCTTGAAAGAGCAAC | Cloning ORF-V5-MS2BP constructs | This work |
| **ORF9c** | ORF9c-V5-MS2BP_rev | GGGGACCACTTTGTACAAGAAAGCTGGGTTATCCGTAAGACAGCAGCACA | Cloning ORF-V5-MS2BP constructs | This work |

**Supplementary Table 7.** Reference names and sequences of DsiRNAs from IDT

| **siRNA target gene** | **DsiRNA name** | **Sequence 5ʹ —> 3ʹ (+strand, –strand)#** |
| --- | --- | --- |
| **HSPA5** | hs.Ri.HSPA5.13.1 | rUrCrUrArCrArGrCrUrUrCrUrGrArUrArArUrCrArArCrCAA, rUrUrGrGrUrUrGrArUrUrArUrCrArGrArArGrCrUrGrUrArGrArArA |
| **NDUFA4** | hs.Ri.NDUFA4.13.1 | rArArArUrCrArUrGrUrUrGrGrArGrArUrCrUrCrUrArUrUGT, rArCrArArUrArGrArGrArUrCrUrCrCrArArCrArUrGrArUrUrUrCrA |
| **LAMP1** | hs.Ri.LAMP1.13.1 | rArArGrGrArArUrCrCrArGrUrUrGrArArUrArCrArArUrUCT, rArGrArArUrUrGrUrArUrUrCrArArCrUrGrGrArUrUrCrCrUrUrGrU |
| **LAPTM4A** | hs.Ri.LAPTM4A.13.1 | rArGrUrArUrGrArArGrUrCrArUrCrGrGrUrArArUrUrArCTA, rUrArGrUrArArUrUrArCrCrGrArUrGrArCrUrUrCrArUrArCrUrGrA |
| **PSMD13** | hs.Ri.PSMD13.13.3 | rCrGrGrUrUrUrGrArGrArArUrGrUrUrCrCrUrArUrArArUAA, rUrUrArUrUrArUrArGrGrArArCrArUrUrCrUrCrArArArCrCrGrCrA |
| **LDHB** | hs.Ri.LDHB.13.3 | rGrArGrCrCrUrUrUrArGrUrUrUrUrCrArUrCrCrArUrGrUAC, rGrUrArCrArUrGrGrArUrGrArArArArCrUrArArArGrGrCrUrCrGrA |
| **UGGT1** | hs.Ri.UGGT1.13 TriFECTa Kit DsiRNA | rArGrCrUrGrArGrArUrGrUrUrCrCrUrUrArGrUrArArUrCAT,  rArUrGrArUrUrArCrUrArArGrGrArArCrArUrCrUrCrArGrCrUrGrU; rGrGrCrUrCrArGrCrUrGrArUrArArArCrArUrGrArArUrCTG, rCrArGrArUrUrCrArUrGrUrUrUrArUrCrArGrCrUrGrArGrCrCrArA; rGrArArUrGrGrArArArUrGrUrArUrUrGrGrUrArArArGrCTA, rUrArGrCrUrUrUrArCrCrArArUrArCrArUrUrUrCrCrArUrUrCrCrC |
| **RPN1** | hs.Ri.RPN1.13.1 | rGrUrUrCrUrGrArArGrUrCrUrArArGrArUrArUrUrUrUrUCA, rUrGrArArArArArUrArUrCrUrUrArGrArCrUrUrCrArGrArArCrArG |
| **ACE2** | hs.Ri.ACE2.13.1 | rArGrUrGrArUrGrUrUrUrGrGrArArUrCrGrArUrCrArUrGCT,  rUrGrUrCrArCrUrArCrArArArCrCrUrUrArGrCrUrArGrUrArCrGrA |
| **Scrambled** | Scrambled Negative Control DsiRNA | /5Phos/rCrUrUrCrCrUrCrUrCrUrUrUrCrUrCrUrCrCrCrUrUrGrUGA,  rUrCrArCrArArGrGrGrArGrArGrArArArGrArGrArGrGrArArGrGrA |
| **Transfection Control** | TYE 563 Transfection Control DsiRNA | /5TYE563/T*CrCrUrUrCrCrUrCrUrCrUrUrUrCrUrCrUrCrCrCrUrUrGrUG*A,  /5TYE563/T*CrArCrArArGrGrGrArGrArGrArArArGrArGrArGrGrArArGG*A |

#where rN is a ribonucleoside for rN ={rA,rU,rC,rG}, and N without any r prepended is a deoxyribonucleoside for N={A,T,C,G}

**Supplementary Table 8.** Selected siRNA genes targeted by SARS-CoV-2 translation enhancing proteins with eCLIP peak -log10(p-values) indicated in a blue cell

| **siRNA Target** | **NSP2** | **NSP3** | **NSP12** | **ORF3b** | **ORF7b** | **ORF9c** |
| --- | --- | --- | --- | --- | --- | --- |
| **PSMD13** | [8.71] | [4.603] | [5.002] |  |  |  |
| **UGGT1** | [3.469] |  | [5.1, 5.985] |  |  |  |
| **RPN1** |  | [9.438] | [6.58] |  | [4.981] | [3.837] |
| **LAMP1** |  | [15.722] | [26.587] |  | [8.93] | [7.293] |
| **NDUFA4** |  |  | [18.694] |  |  |  |
| **LDHB** |  |  | [20.623] |  |  |  |
| **HSPA5** |  |  |  | [12.995] |  |  |

**Supplementary Figure 1**

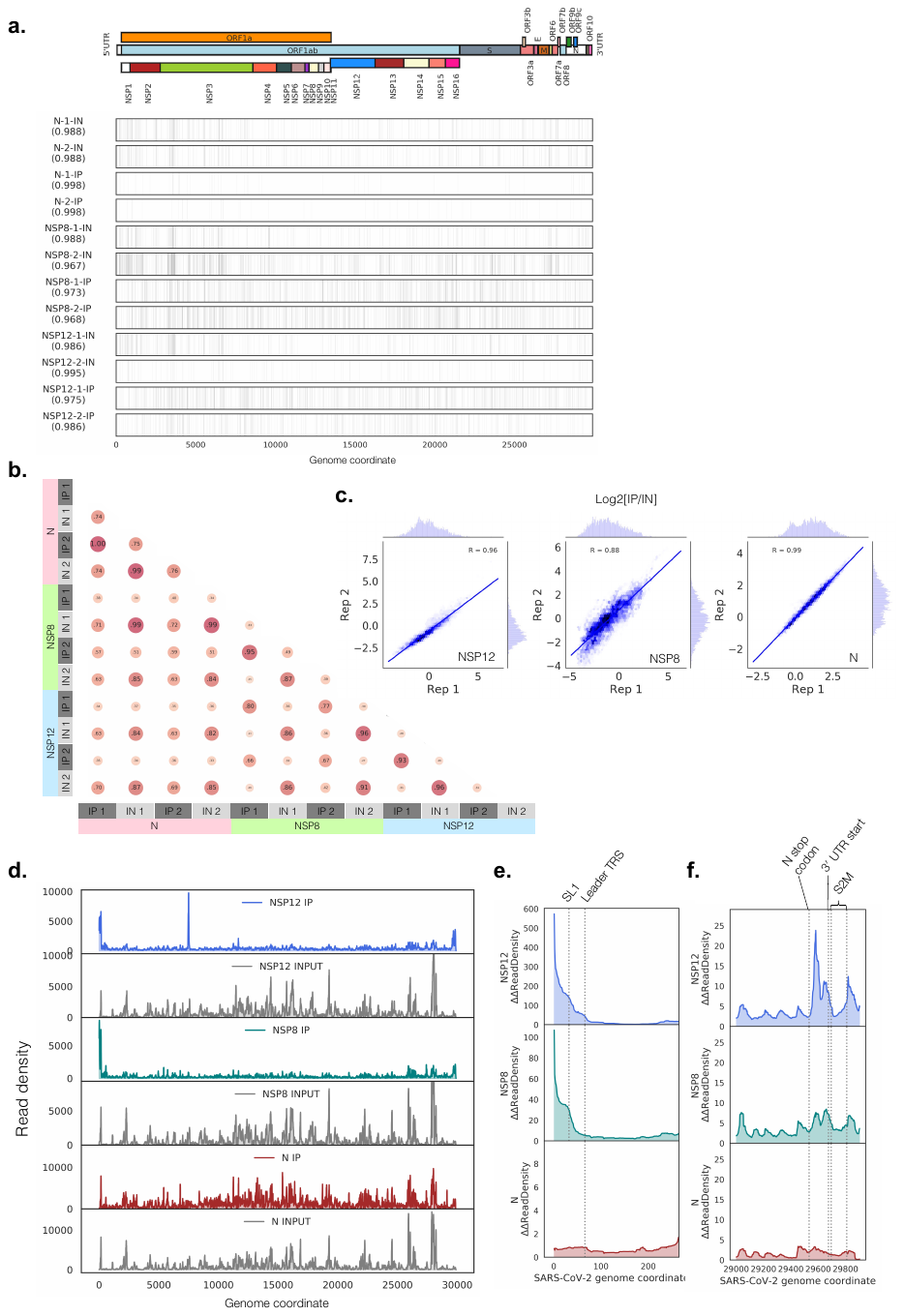
**Supplementary Figure 2**

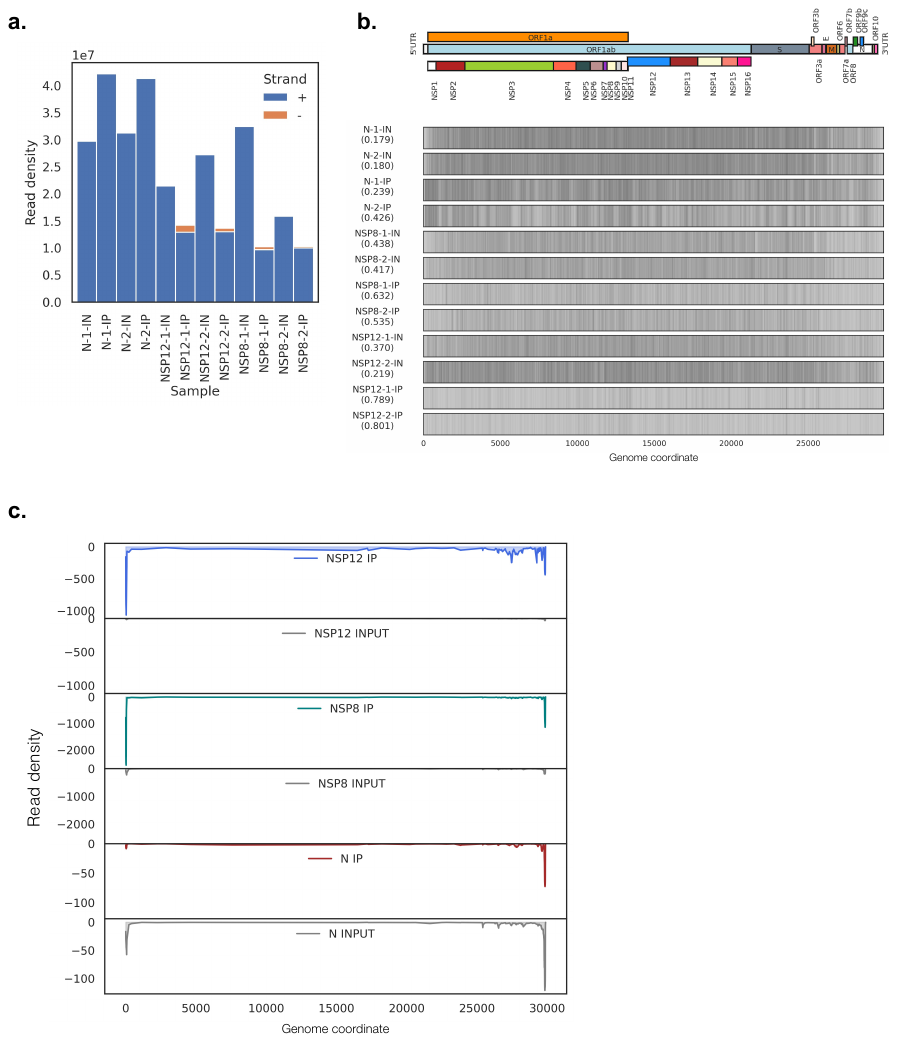

**Supplementary Figure 3**

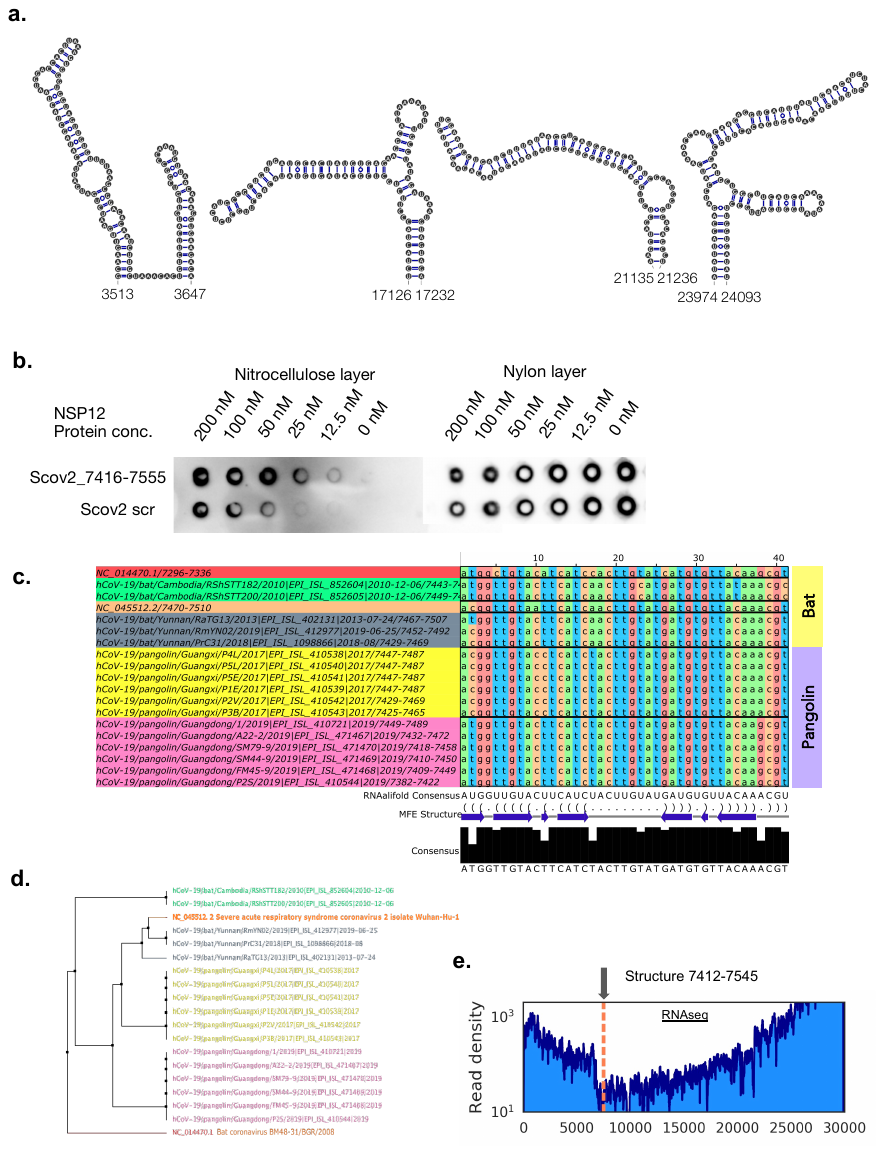

**Supplementary Figure 4**

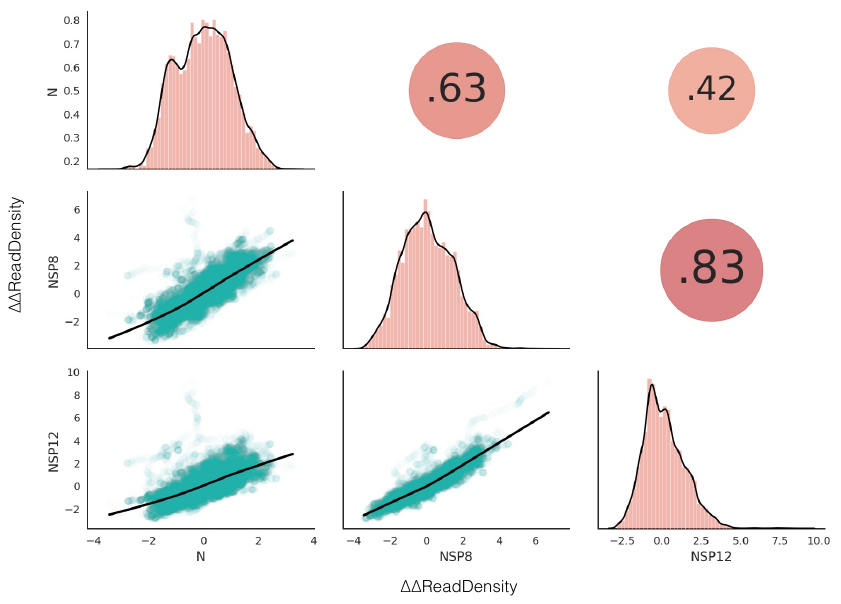
 **Supplementary Figure 5**

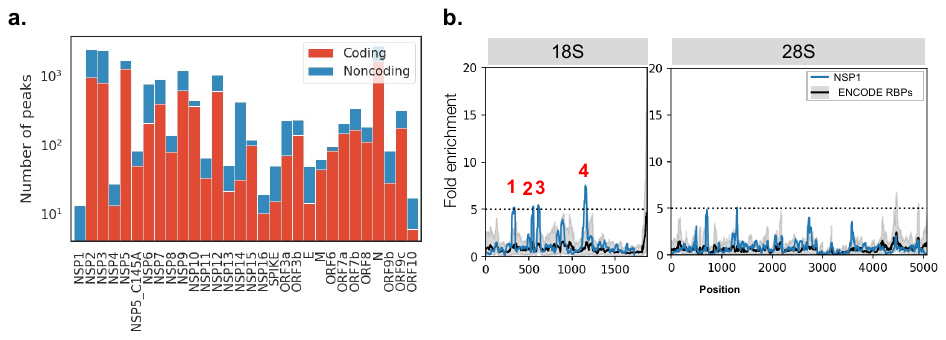

**Supplementary Figure 6**

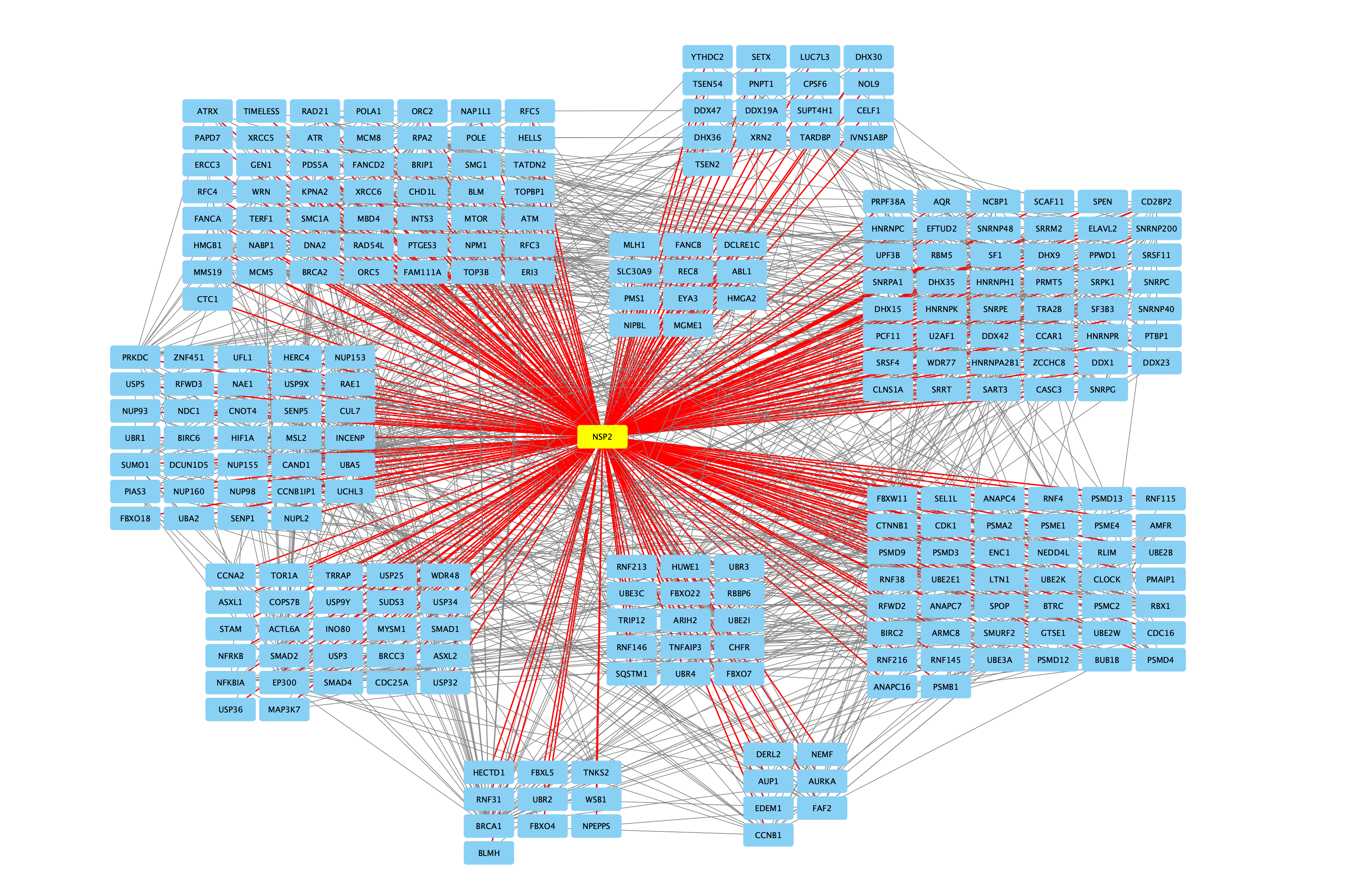

DNA metabolic process

RNA processing

mRNA splicing, via spliceosome

proteasomal protein catabolic process

proteasome-mediated ubiquitin-dependent protein catabolic process

protein polyubiquitination

protein modification by small protein removal

protein modification by small protein conjugation

ubiquitin-dependent protein catabolic process

DNA repair

NSP2

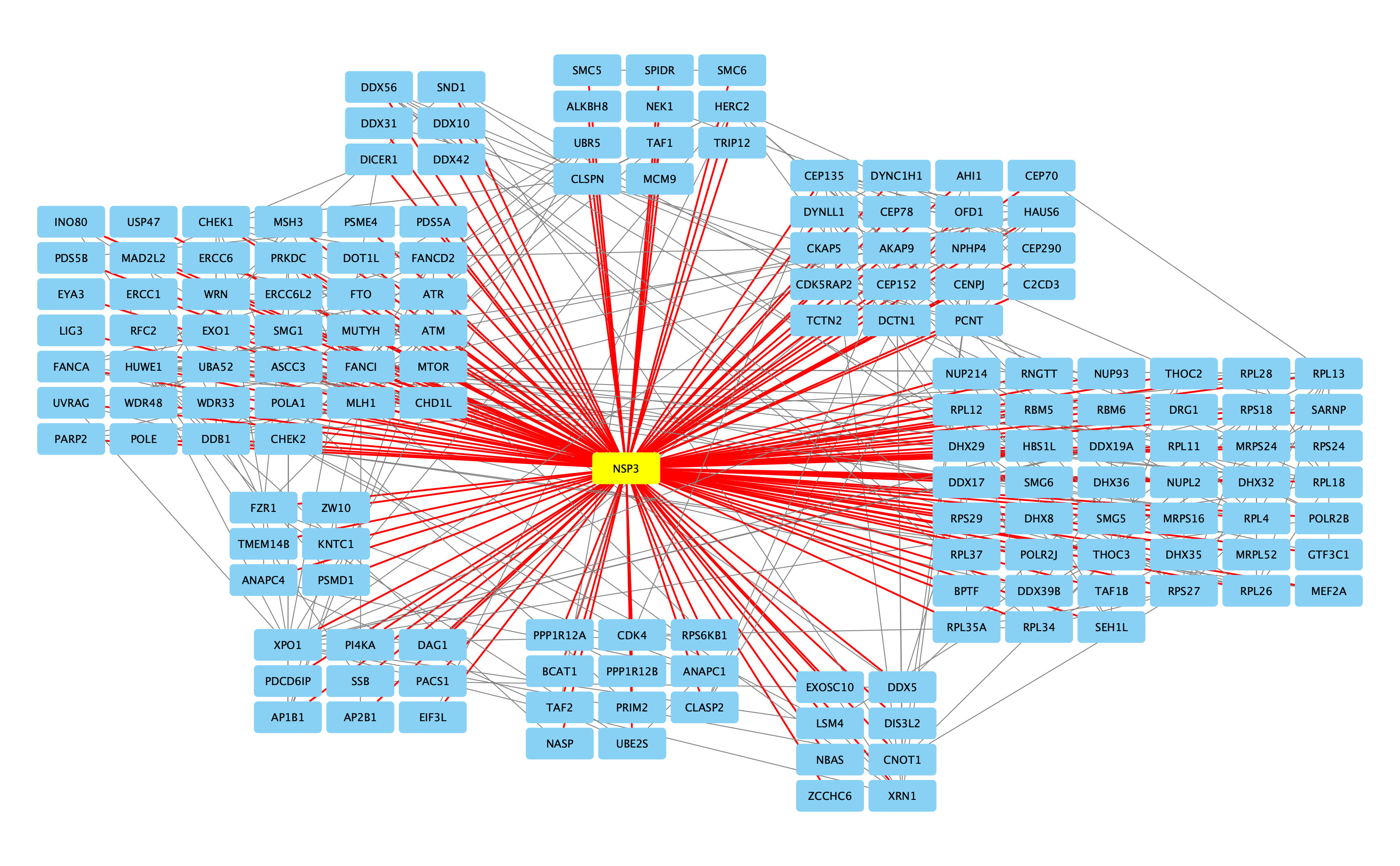

mitotic cell cycle phase transition

nuclear-transcribed mRNA catabolic process

viral process

regulation of mitotic cell cycle phase transition

gene expression

ciliary basal body-plasma membrane docking

RNA metabolic process

cellular macromolecule biosynthetic process

DNA repair

NSP3

**Supplementary Figure 6**

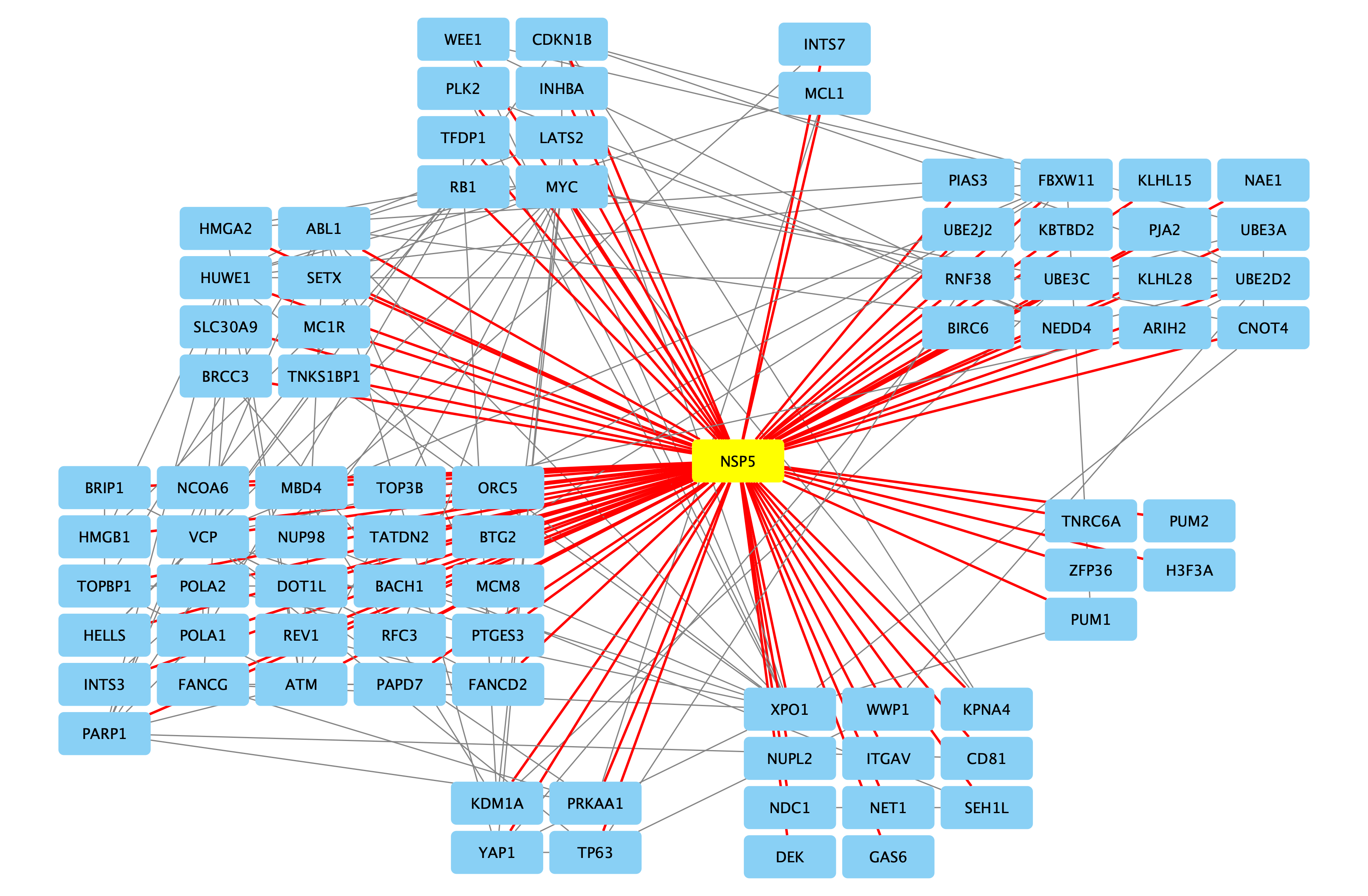

DNA metabolic process

DNA repair

protein modification by small protein conjugation

regulation of gene silencing by miRNA

response to gamma radiation

cellular response to DNA damage stimulus

G1/S transition of mitotic cell cycle

viral life cycle

NSP5

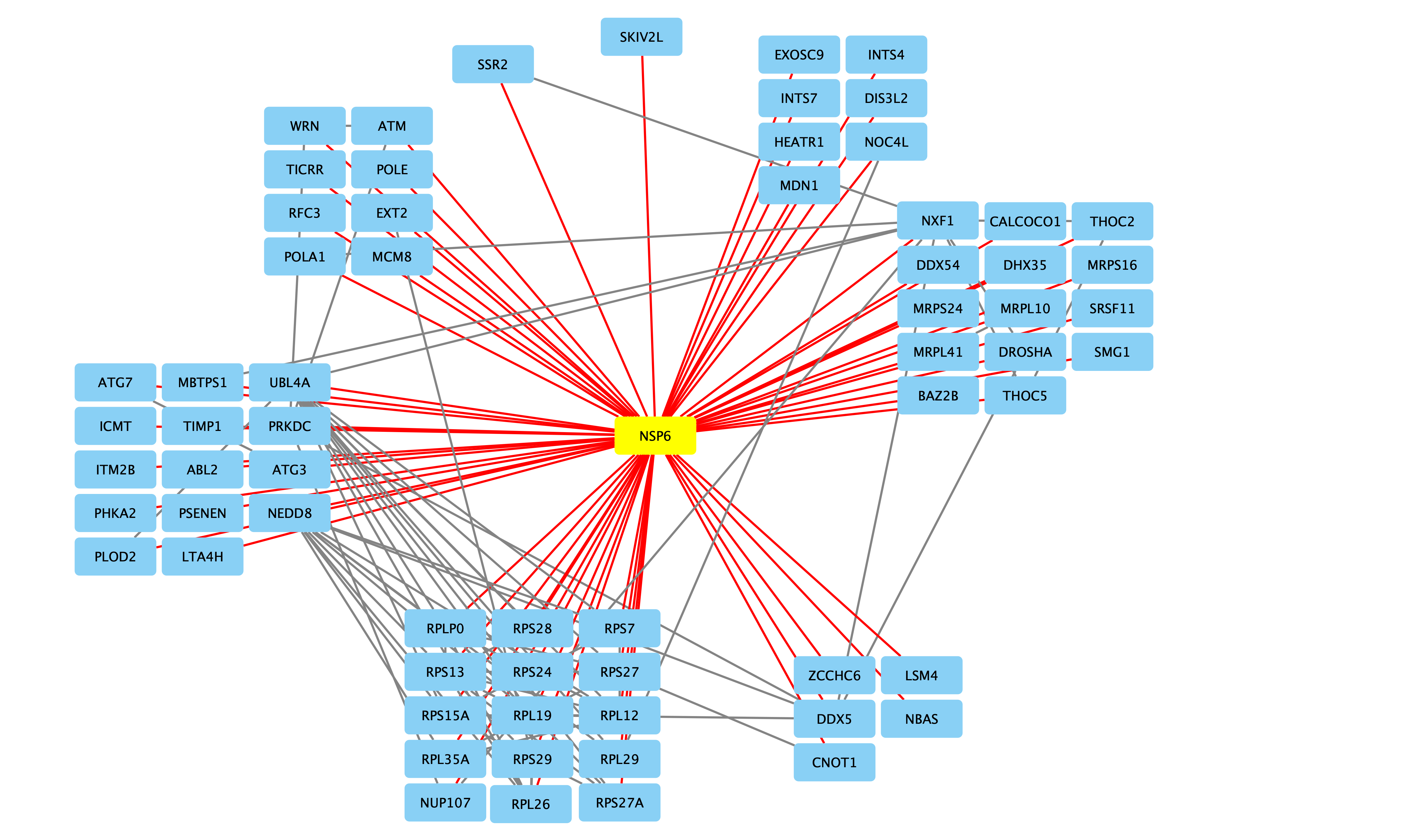

cellular protein metabolic process

cellular macromolecule biosynthetic process

cotranslational protein targeting to membrane

nuclear-transcribed mRNA catabolic process, nonsense-mediated decay

viral transcription

nuclear-transcribed mRNA catabolic process

gene expression

ncRNA processing

NSP6

**Supplementary Figure 6**

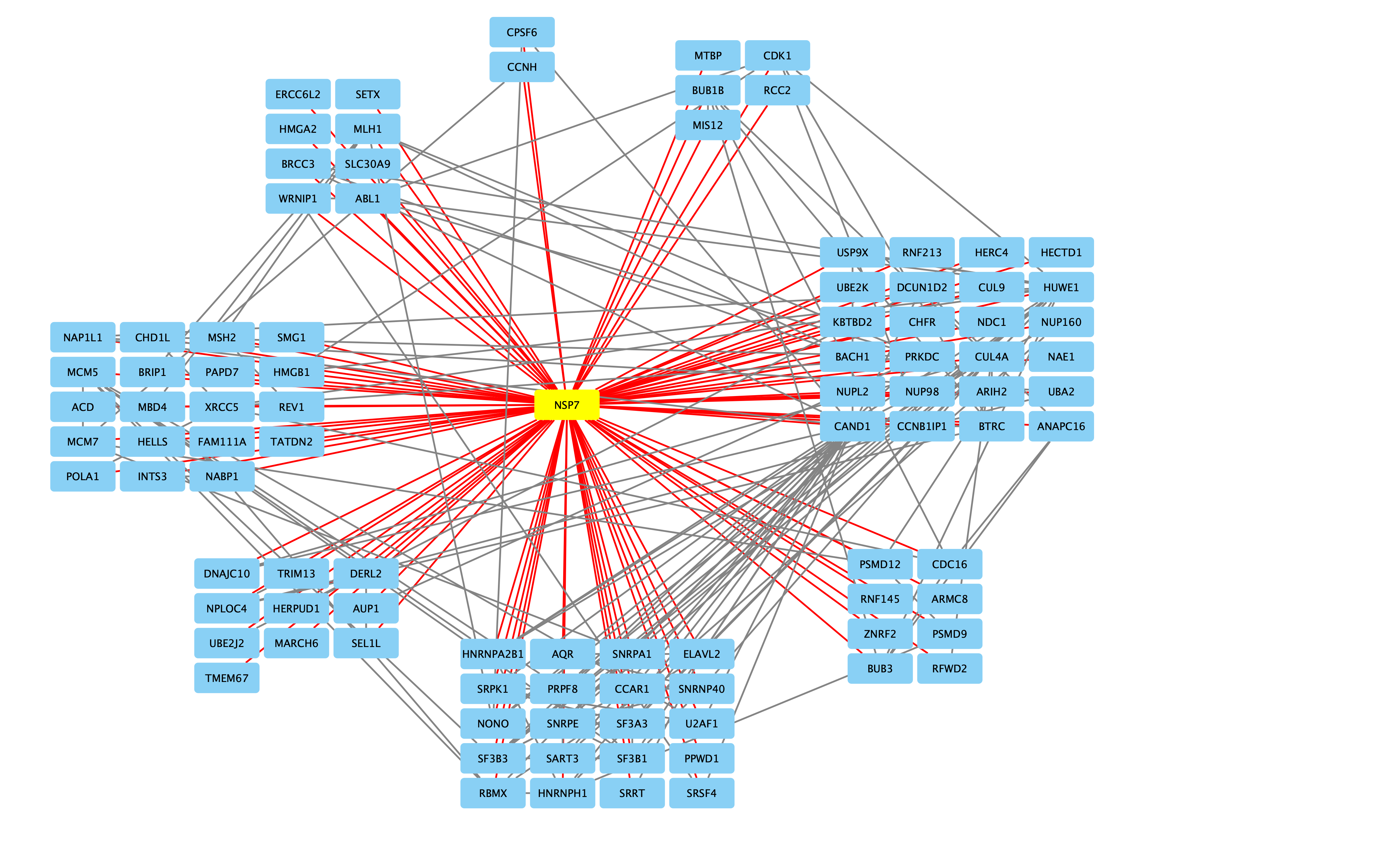

NSP7

DNA metabolic process

DNA repair

mRNA processing

protein localization to kinetochore

protein modification by small protein conjugation

proteasome-mediated ubiquitin-dependent protein catabolic process

mRNA splicing, via spliceosome

ERAD pathway

**Supplementary Figure 6**

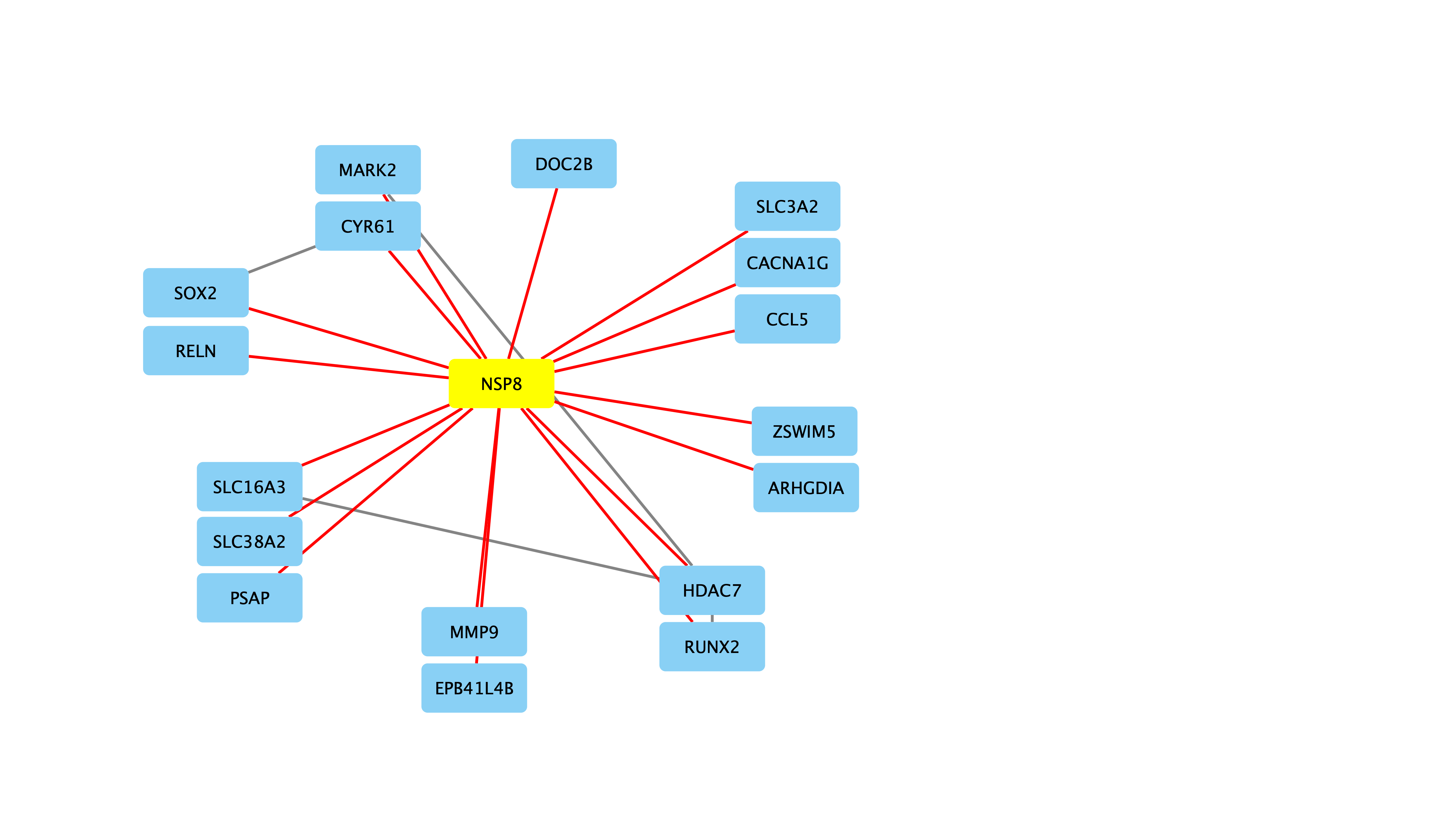

positive regulation of protein kinase activity

ATP-dependent chromatin remodeling

positive regulation of regulated secretory pathway

divalent metal ion transport

regulation of axonogenesis

regulation of osteoblast differentiation

carboxylic acid transport

regulation of keratinocyte migration

NSP8

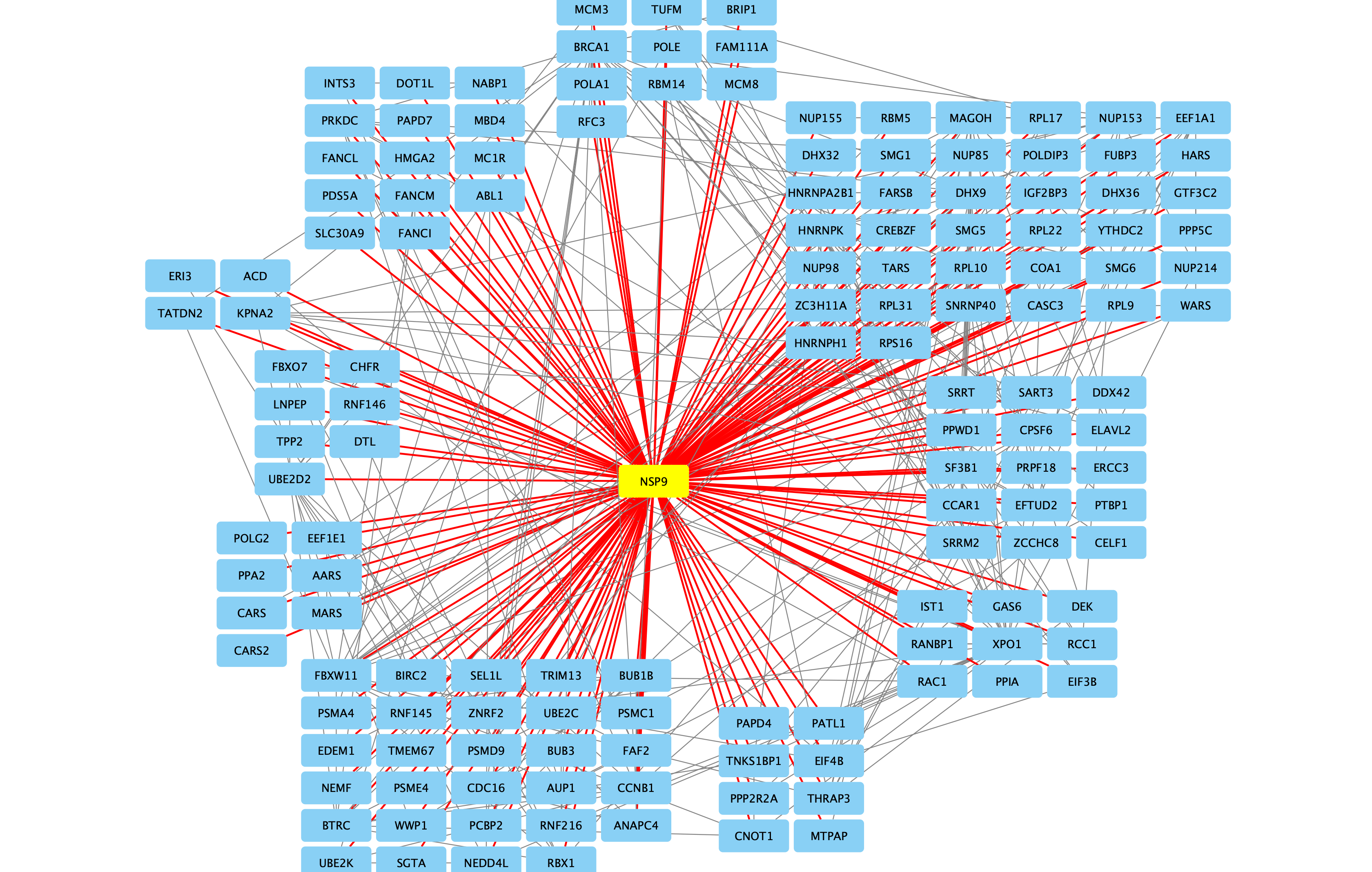

mRNA processing

nuclear-transcribed mRNA catabolic process

DNA metabolic process

gene expression

viral process

protein polyubiquitination

Cellular macromolecule biosynthetic process

DNA repair

tRNA aminoacylation for protein translation

proteasome-mediated ubiquitin-dependent protein catabolic process

NSP9

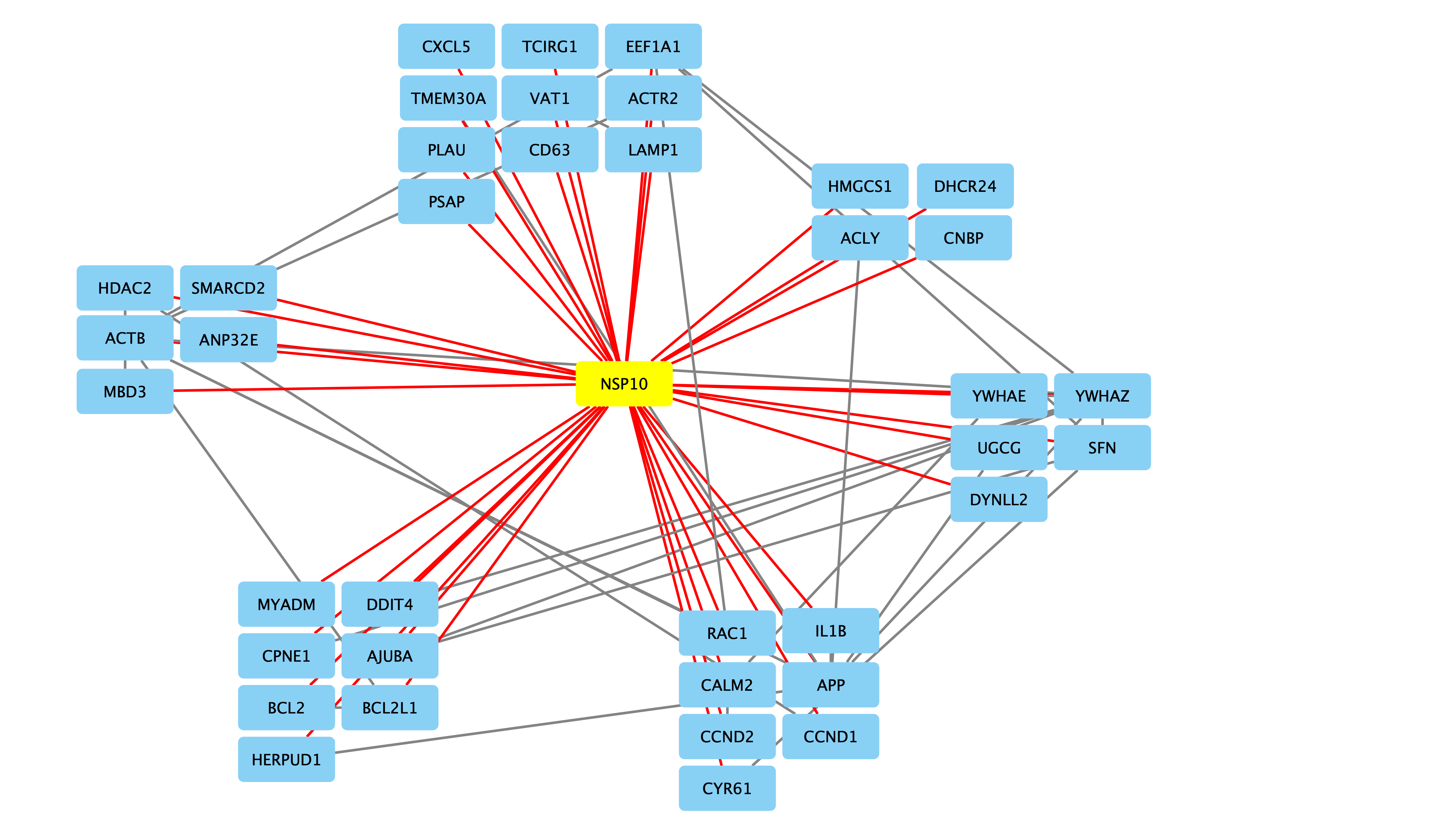

NSP10

neutrophil mediated immunity

ATP-dependent chromatin remodeling

negative regulation of intracellular signal transduction

positive regulation of protein modification process

positive regulation of establishment of protein localization to mitochondrion

secondary alcohol biosynthetic process

**Supplementary Figure 6**

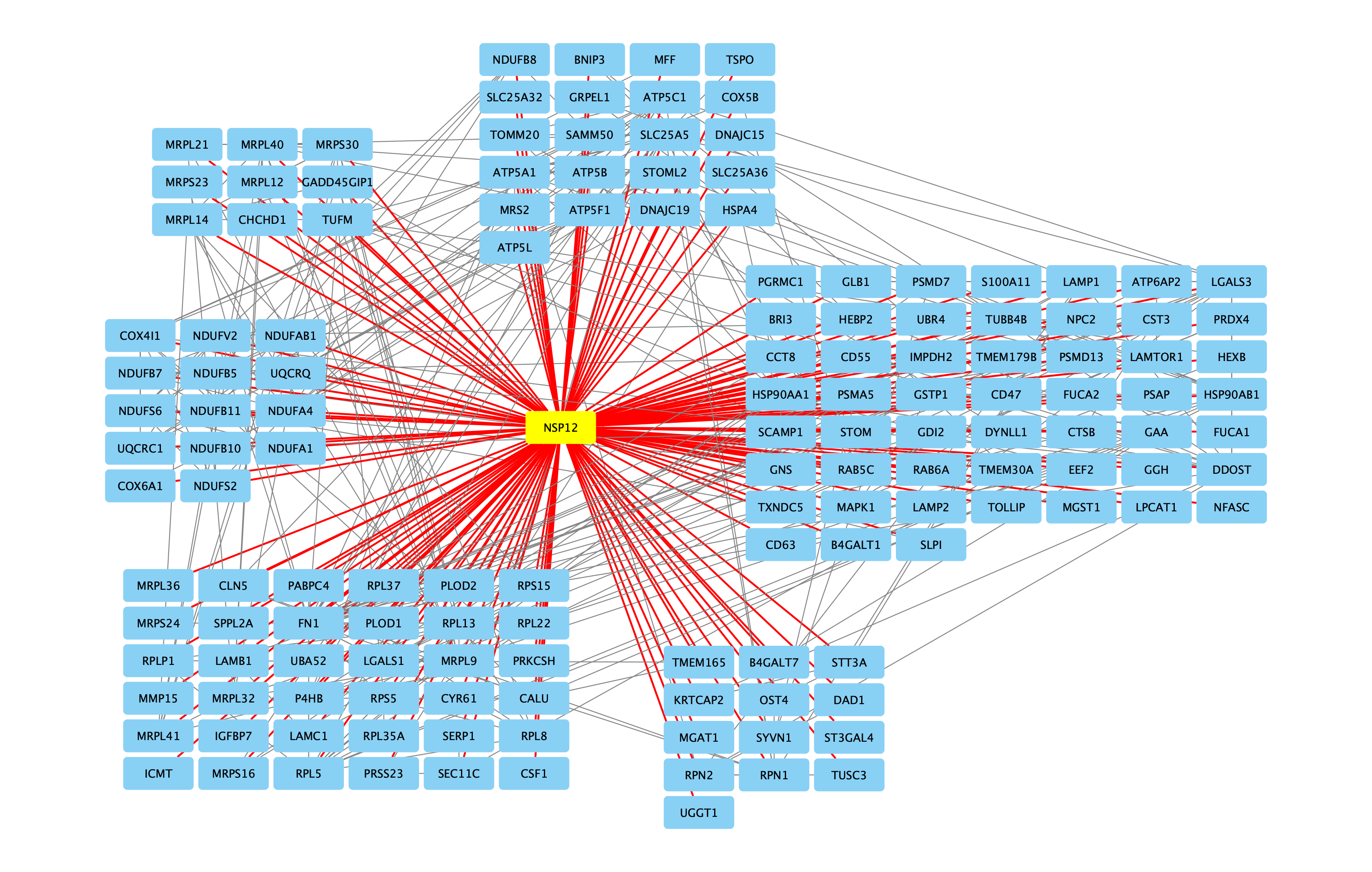

mitochondrial transport

mitochondrial translational elongation

neutrophil mediated immunity

protein N-linked glycosylation

mitochondrial ATP synthesis coupled electron transport

cellular protein metabolic process

NSP12

**Supplementary Figure 6**

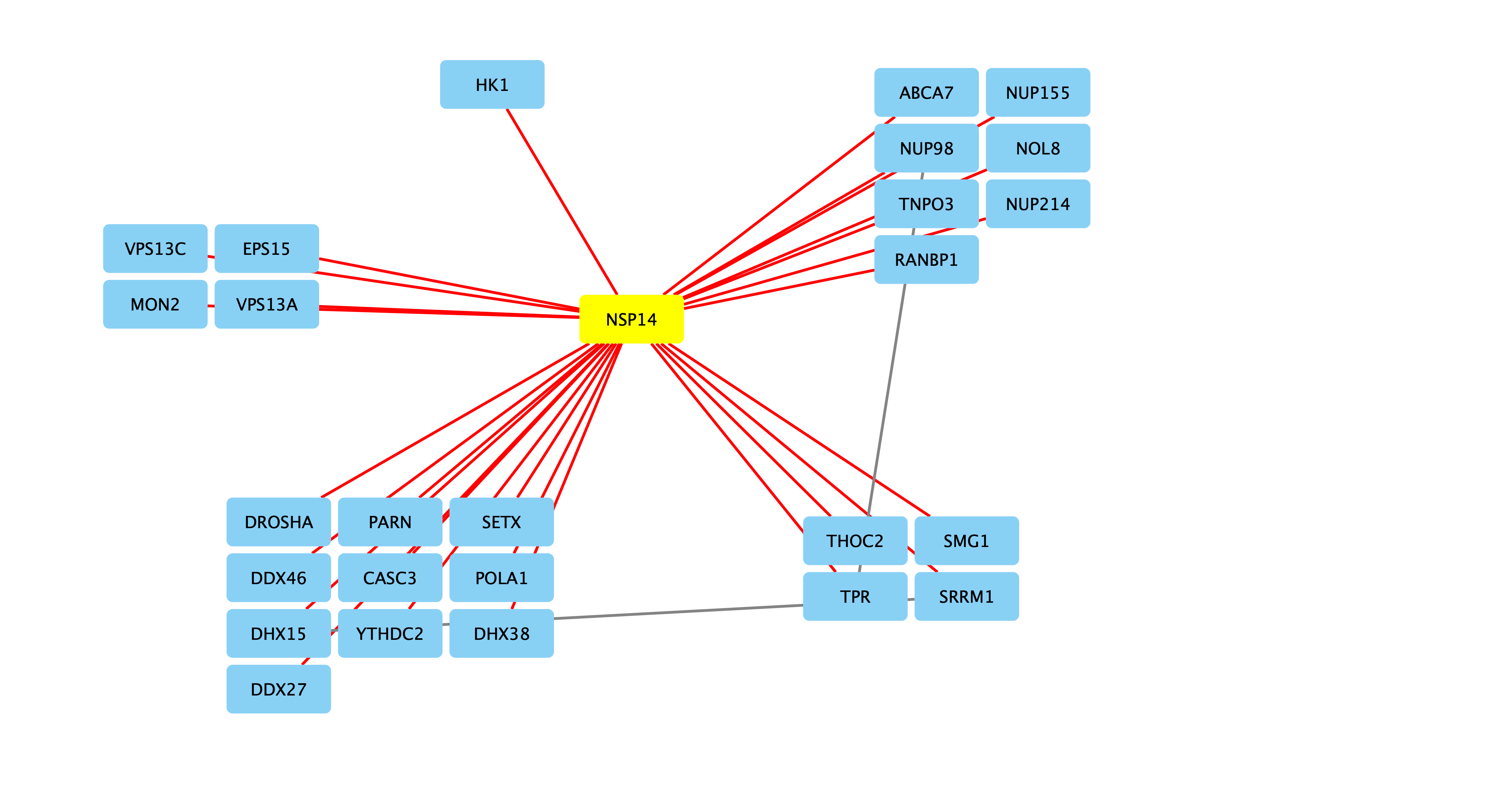

Golgi to endosome transport

establishment of protein localization to organelle

protein localization to nucleus

mRNA export from nucleus

RNA metabolic process

NSP14

**Supplementary Figure 6**

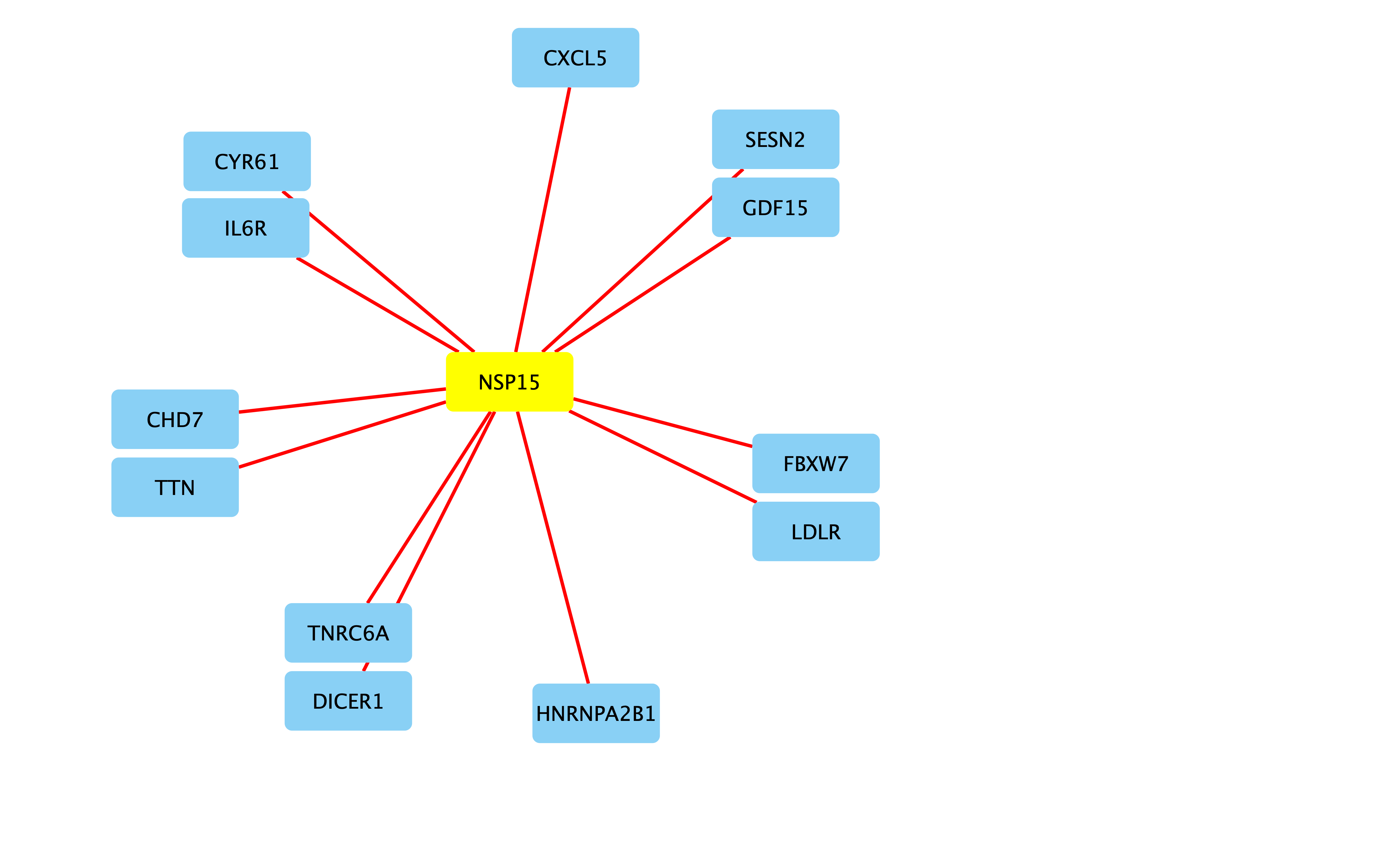

positive regulation of ossification

positive regulation of chemotaxis

regulation of protein phosphorylation

regulation of triglyceride biosynthetic process

production of miRNAs involved in gene silencing by miRNA

gene silencing by miRNA

heart morphogenesis

NSP15

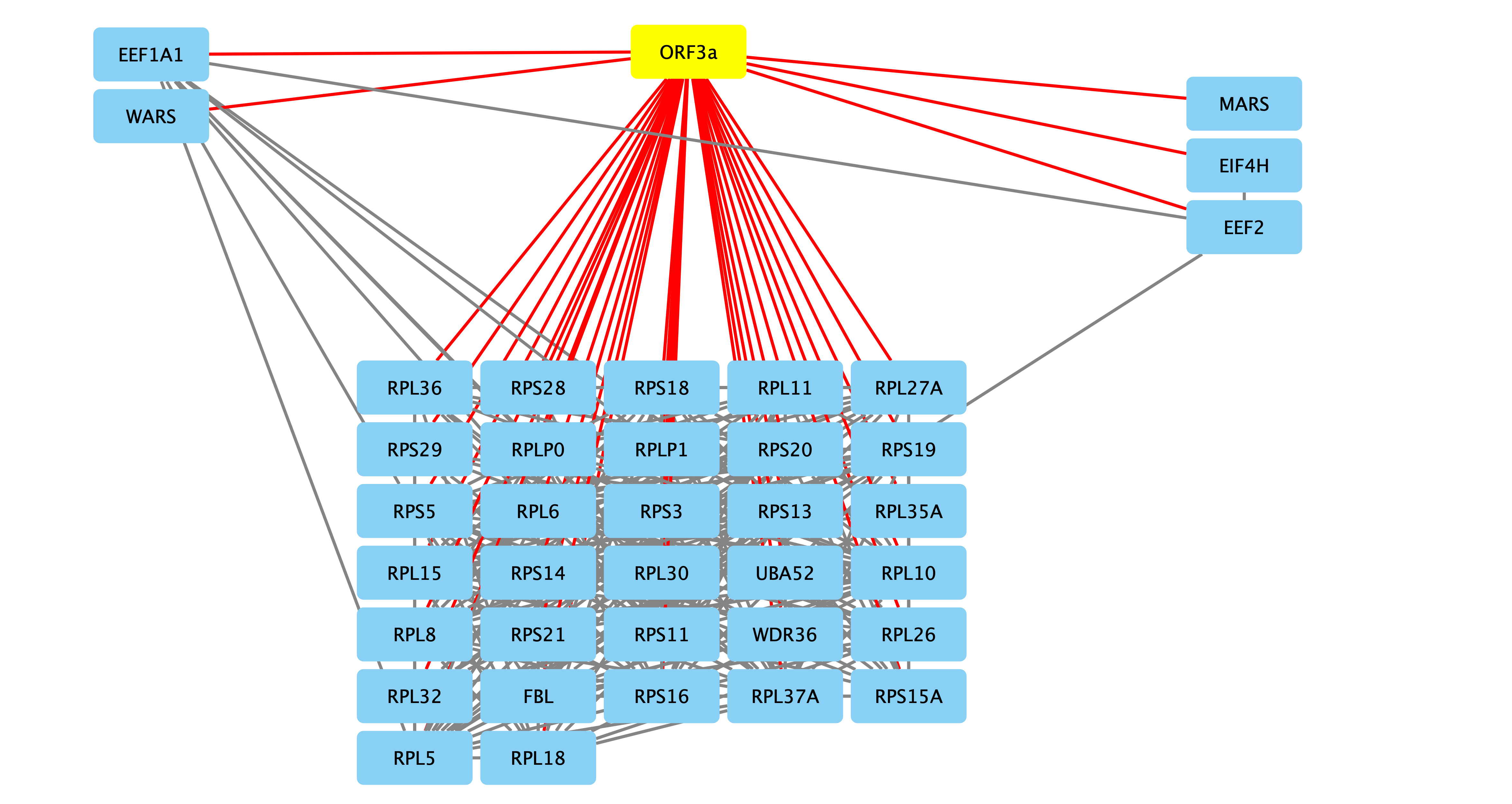

rRNA processing

translation

peptide biosynthetic process

ORF3a

**Supplementary Figure 6**

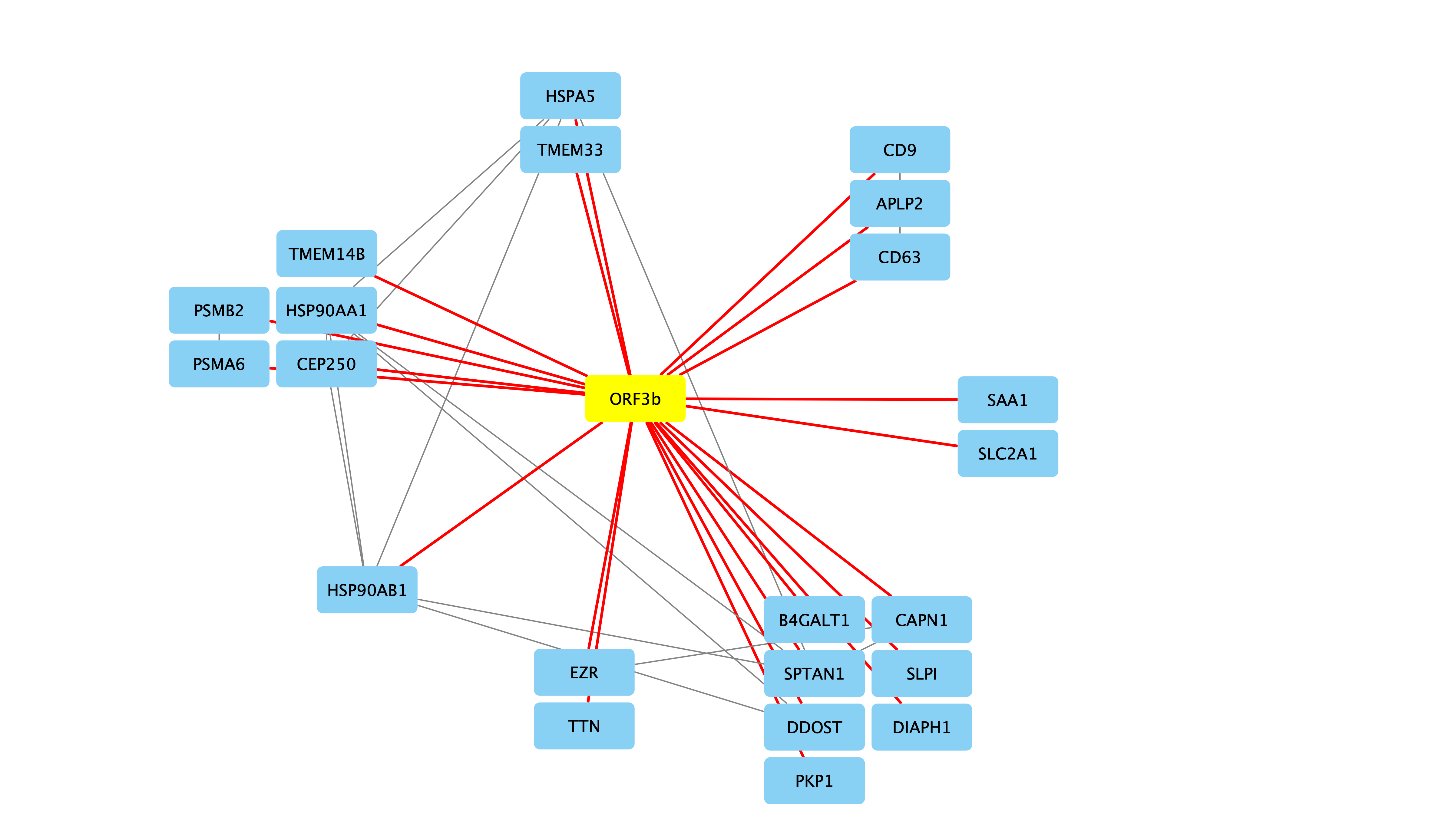

regulation of protein secretion

regulation of IRE1-mediated unfolded protein response

platelet degranulation

neutrophil mediated immunity

regulation of mitotic cell cycle phase transition

protein kinase A signaling

response to unfolded protein

ORF3b

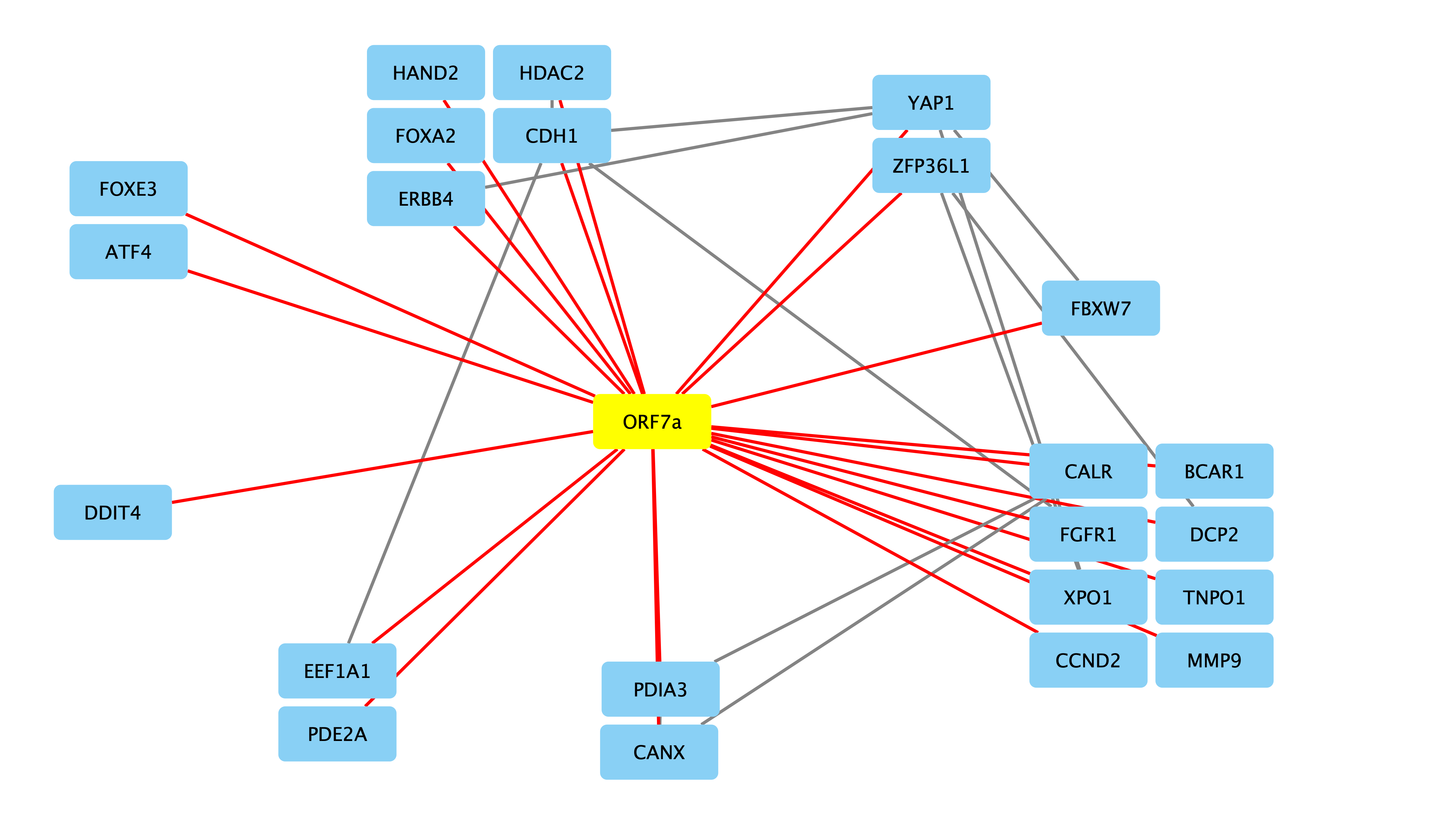

ORF7a

regulation of stem cell proliferation

negative regulation of multicellular organismal process

regulation of apoptotic process

antigen processing and presentation of peptide antigen via MHC class I

cellular response to growth factor stimulus

generation of neurons

mRNA transcription from RNA polymerase II promoter

positive regulation of nucleic acid-templated transcription

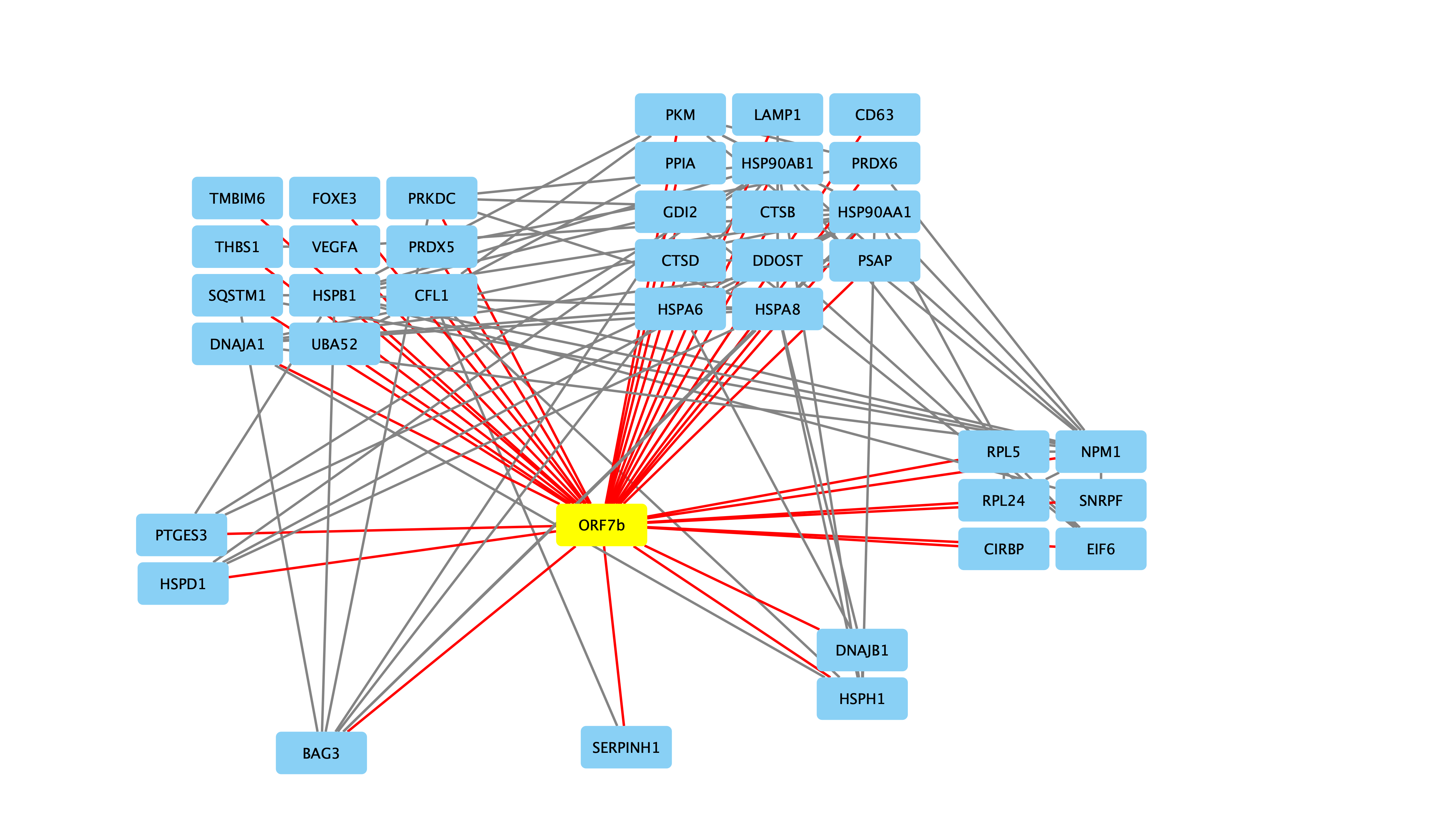

negative regulation of programmed cell death

neutrophil activation involved in immune response

ribonucleoprotein complex assembly

'de novo' posttranslational protein folding

response to unfolded protein

regulation of cellular response to heat

chaperone-mediated protein complex assembly

ORF7b

**Supplementary Figure 6**

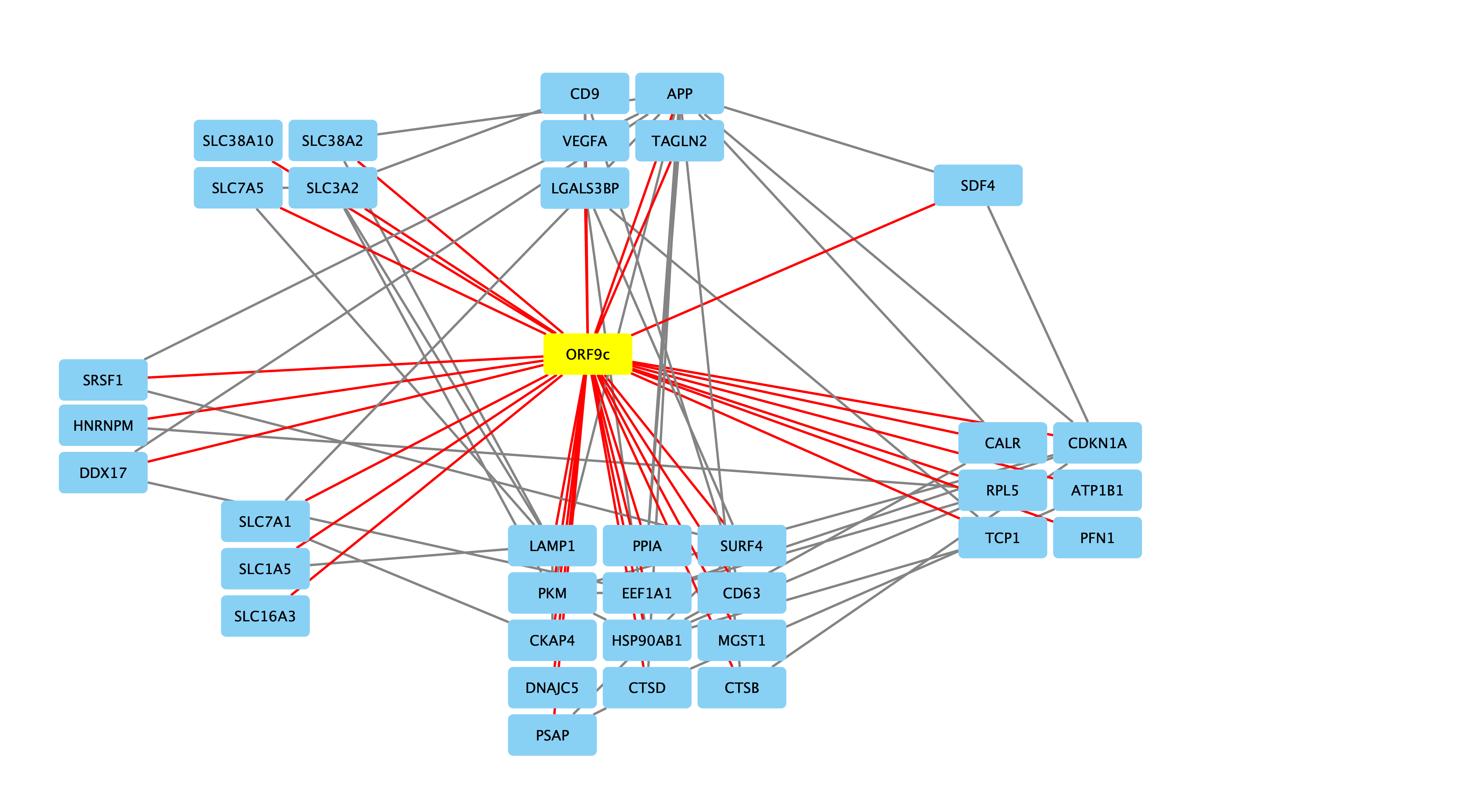

protein stabilization

carboxylic acid transport

alternative mRNA splicing, via spliceosome

amino acid transmembrane transport

platelet degranulation

regulated exocytosis

neutrophil mediated immunity

ORF9c

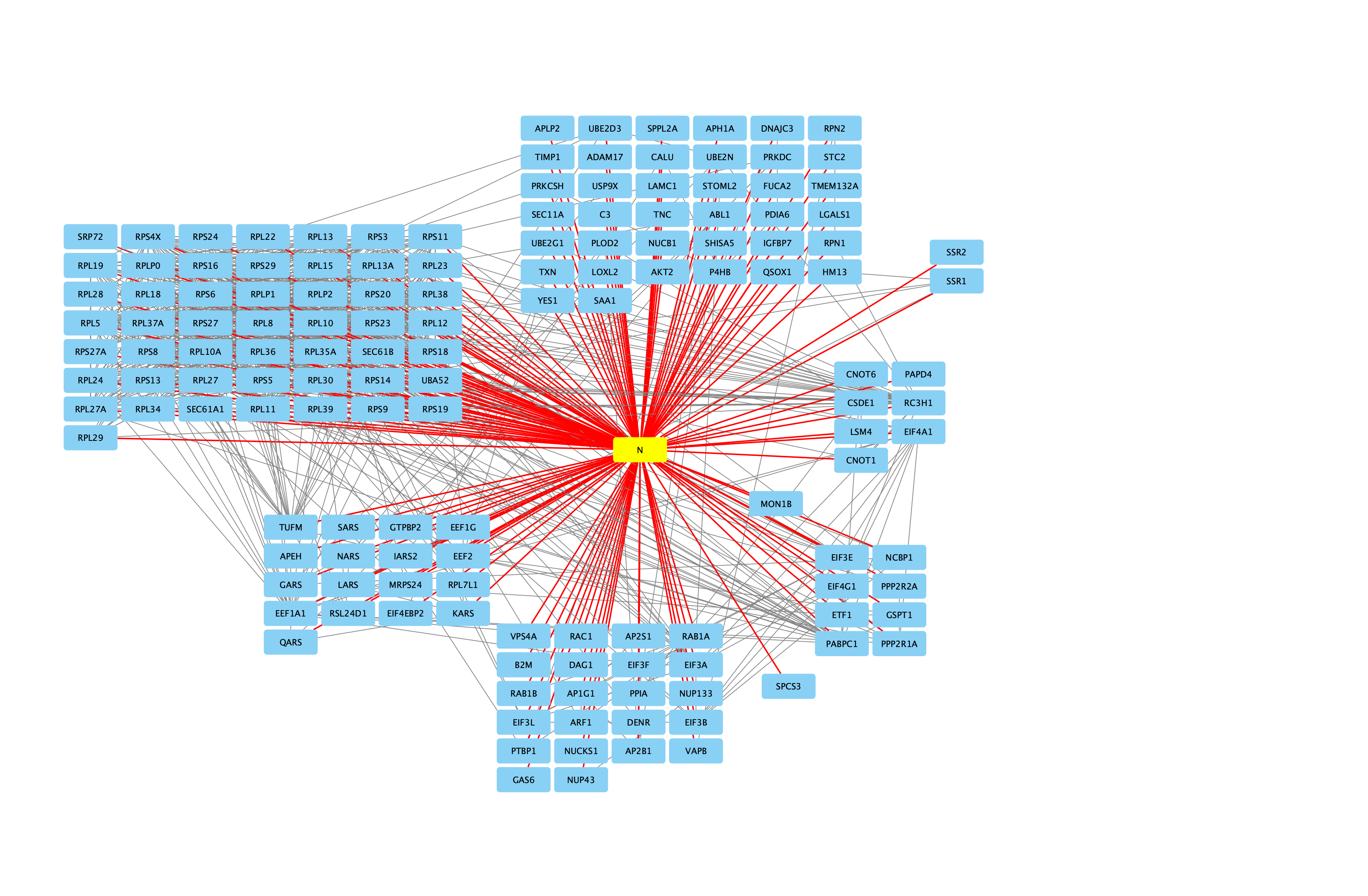

cellular protein metabolic process

viral process

viral transcription

translation

protein targeting to ER

cotranslational protein targeting to membrane

SRP-dependent cotranslational protein targeting to membrane

nonsense-mediated decay

nuclear transcribed mRNA catabolic process

N

**Supplementary Figure 7**

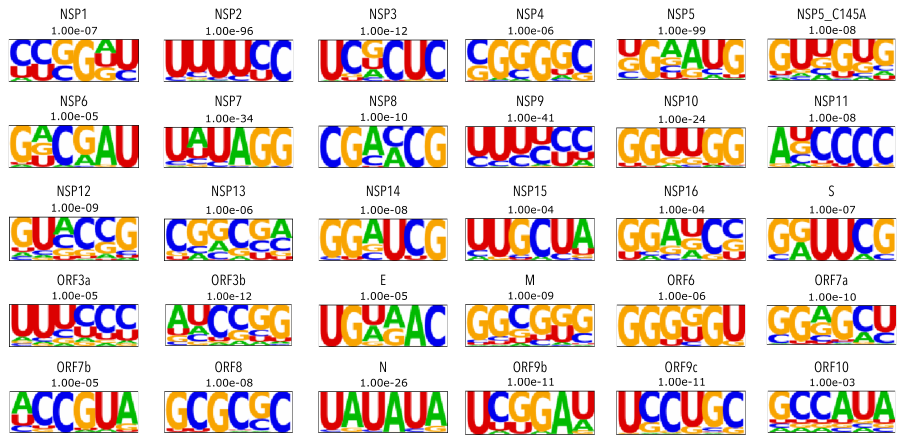

**Supplementary Figure 8**

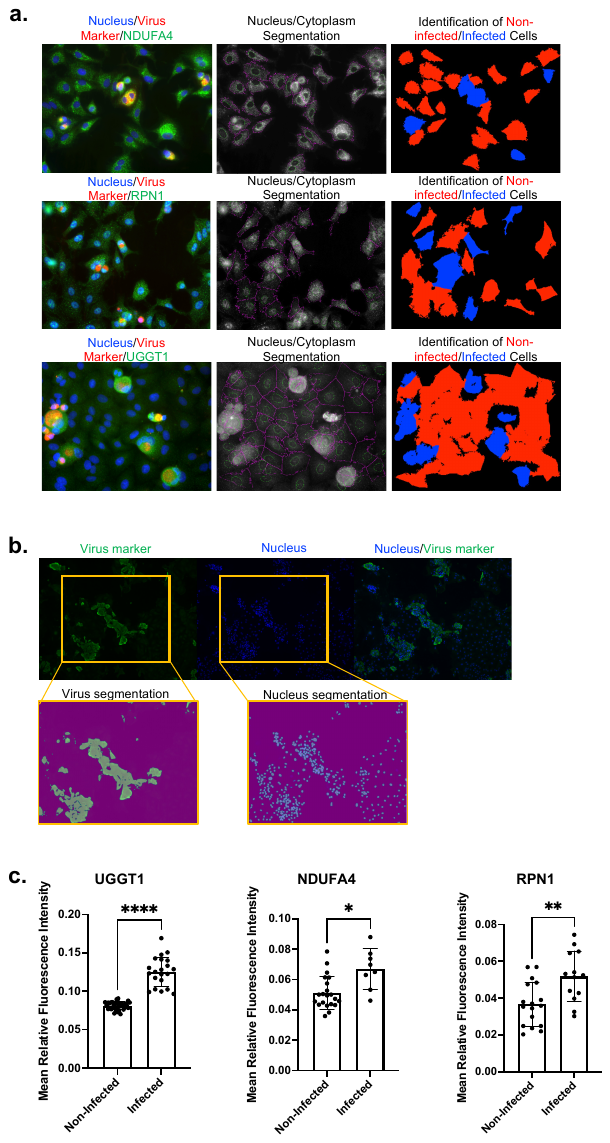

**Supplementary Figure 9**

^[1]^
